# Spatiotemporal transcriptomic dynamic across the auditory pathway from young to old age in 5xFAD Alzheimer disease mice

**DOI:** 10.64898/2026.10.17.759861

**Authors:** Yong Kong, Tian-Ying Zhai, Hong-Bo Zhao

## Abstract

Auditory dysfunction can precede clinical dementia, but the molecular progression of Alzheimer disease (AD) pathology from the sensory periphery to auditory cortex remains poorly resolved. We performed bulk RNA sequencing of auditory cortex (AC), inferior colliculus (IC), cochlear nucleus (CN), and cochlea from 5xFAD and wild-type mice at postnatal day 50 (P50), 8 months, 11 months, and 16 months. Age-specific genotype effects and Genotype×Age interactions were evaluated using DESeq2, followed by Hallmark, KEGG, and Gene Ontology gene-set enrichment analyses and a cross-region synthesis of interaction trajectories. At P50, AC, IC, and cochlea showed coordinated suppression of mitochondrial metabolism, translation, RNA processing, and proteostasis, whereas CN displayed the opposite metabolic and biosynthetic profile together with reduced synaptic and adhesion programs. At 8 months, inflammatory interactions converged across all four regions, including interferon, complement, IL6–JAK–STAT3, and broader immune-response pathways. This coordination fractured at 11 months: AC and IC retained inflammatory and negative synaptic shifts, whereas CN and cochlea frequently changed in the opposite direction, including transient positive shifts in synaptic programs. By 16 months, immune interactions converged again, while metabolic, proteostatic, and synaptic trajectories remained region dependent; CN opposed AC, IC, and cochlea in several major metabolic programs. These findings define a distributed, asynchronous response rather than a monotonic bottom-up or top-down progression. The resulting spatiotemporal map links early metabolic stress, intermediate central–peripheral decoupling, and late inflammatory convergence, providing a molecular framework for evaluating peripheral-to-central auditory gain and temporal-processing measures as candidate indicators of AD progression.

## Introduction

Alzheimer disease (AD) develops across a prolonged biological continuum in which amyloid accumulation, altered brain metabolism, synaptic dysfunction, and neuroinflammation precede dementia. In dominantly inherited AD, molecular and imaging abnormalities emerge years before expected symptom onset, motivating accessible measures of evolving neural dysfunction (Bateman et al., 2012). Auditory tests are attractive because they probe neural function noninvasively from the sensory periphery through the brainstem and cortex. The relevant phenotype may extend beyond reduced sound sensitivity to impaired temporal resolution, sensory gating, central gain control, and extraction of signals from noise.

The predictive power of auditory deficits in AD rests particularly on these central and integrative functions (Liu et al., 2020; Mei et al., 2021). Longitudinal studies found that impaired central auditory speech processing can precede probable AD and identify older adults at increased risk of dementia (Gates et al., 2002, 2011). Peripheral hearing loss is also associated with incident dementia (Lin et al., 2011), and poor speech-in-noise performance predicted greater dementia risk in UK Biobank (Stevenson et al., 2022). These associations do not establish causation or disease specificity. They instead suggest that auditory measures may capture distributed neural stress before cognitive symptoms dominate, especially when combined with audiometry, cognitive testing, and established AD biomarkers.

Hearing depends on a vertically organized and recurrent network (Fig. 1A&B). Signals transduced in the cochlea pass through the cochlear nucleus (CN) and inferior colliculus (IC) before reaching auditory cortex (AC), while local inhibitory circuits and descending projections continuously modify activity. A lesion at one node can therefore alter distant activity and homeostatic demand. Auditory brainstem response (ABR) wave I, later-wave-to-wave I relationships, gap detection, and speech-in-noise performance interrogate different levels of this network. Interpreting these measures in AD requires a molecular map that resolves where pathology emerges and whether peripheral and central compartments remain coupled over time (Liu et al., 2020; Mei et al., 2021).

**Figure 1.**
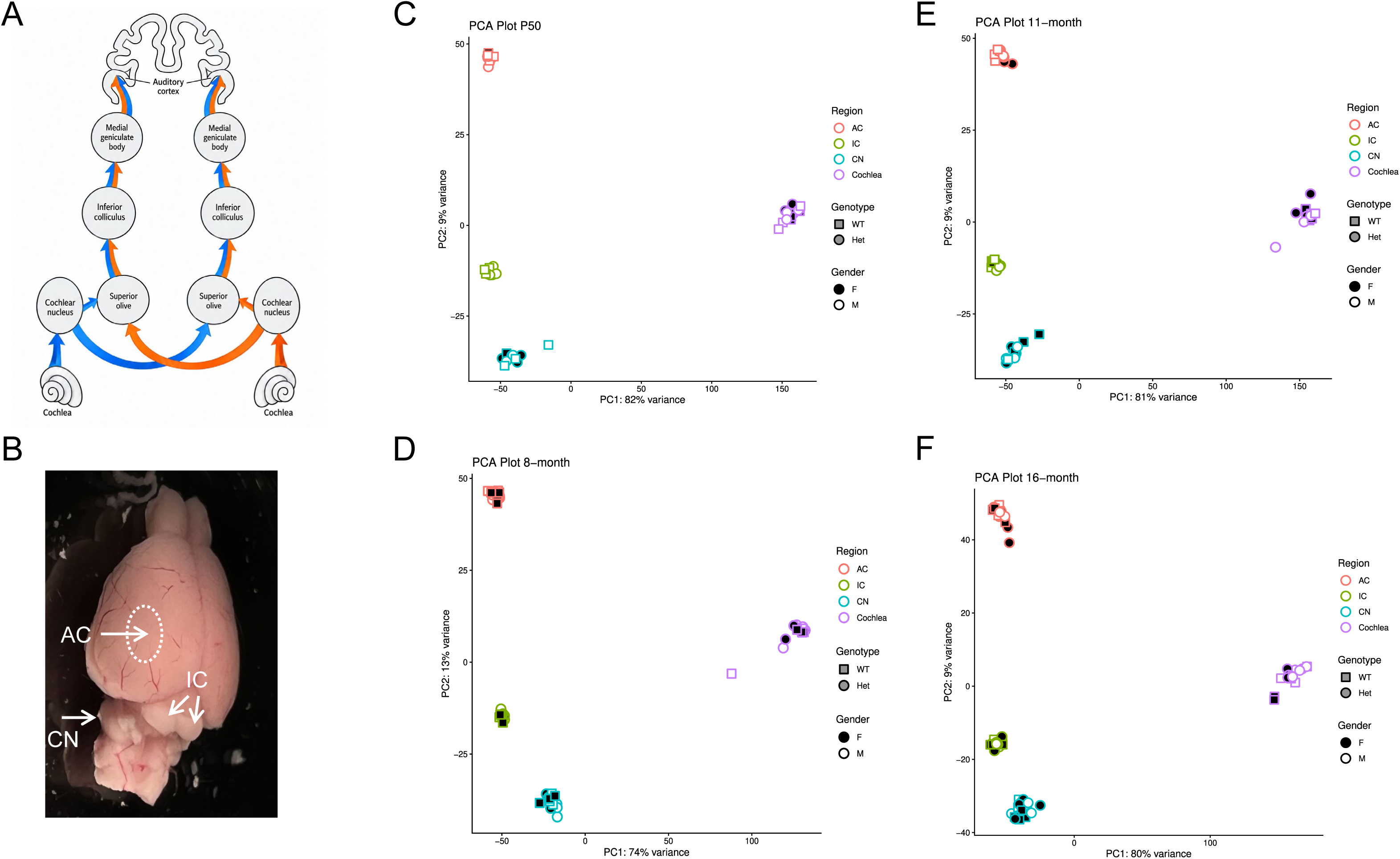
Auditory system. (A), mouse brain (B), and regional separation of auditory transcriptomes (C-F). The cerebellum above the cochlear nucleus (CN) was removed to show CN. AC: auditory cortex, IC: inferior colliculus. Principal-component analysis (PCA) shows clustering of samples according to auditory region.

The 5xFAD model enables such staging. This AD mouse strain co-expresses five familial AD mutations in human amyloid precursor protein and presenilin 1, accumulate abundant Aβ42, and develop rapid amyloid-dominant pathology, and is widely used in AD studies. Intraneuronal Aβ42 appears by approximately 1.5 months; plaques and gliosis begin near 2 months; synaptic degeneration and neuronal loss follow (Oakley et al., 2006). Auditory abnormalities also emerge early. Gap-detection and cortical gap-encoding deficits occur around 2 months (Kaylegian et al., 2019; Weible et al., 2020), auditory cortical Aβ42 is detectable by P21, and plaques later appear in AC and IC while early lower-brainstem plaque burden remains limited (Weible and Wehr, 2022). Increased central auditory gain may precede reduced auditory-nerve output and threshold sensitivity (Mei et al., 2021; Na et al., 2023), whereas later cohorts show peripheral hearing loss and reduced acoustic startle (Liu et al., 2020; O’Leary et al., 2017).

However, the molecular organization of these abnormalities across the auditory pathway remains unclear. We therefore used bulk RNA sequencing to compare 5xFAD and wild-type mice in AC, IC, CN, and cochlea at postnatal day 50 (P50), 8 months, 11 months, and 16 months. These sequential ages span early molecular or peri-plaque change, established plaque-associated disease, advanced synaptic pathology, and late neurodegeneration. We estimated age-specific genotype effects and Genotype×Age interactions and integrated Hallmark, KEGG, and Gene Ontology (GO) enrichment across regions. Evaluating four connected structures across four ages provides a comprehensive spatiotemporal map and tests whether auditory pathology follows an ascending, descending, or asynchronous course. This vertical map also supplies a mechanistic basis for evaluating specific auditory metrics as candidate indicators of systemic AD progression.

## Materials and Methods

### Animals

All procedures and experiments followed in the use of animals were approved by Yale University’s Institutional Animal Care & Use Committee (Yale: 2022-20463) and conformed to the standards of the NIH Guidelines for the Care and Use of Laboratory Animals.

5xFAD transgenic mice were purchased from Jackson Laboratory, United States (Stock No: 006554, mixed C57BL/6;C3H genetic background). Since the C57BL/6 genetic background is associated with age-related hearing loss, we crossed 5xFAD mice with CBA/CaJ mouse strain (Stock No: 00654, Jackson Laboratory, Bar Harbor, ME, United States), which has no apparent age-related hearing loss (Zheng et al., 1999), for at least 6 generations. The genotype of mice was detected by PCR amplification of tail genomic DNA. Both genders were used in experiments. WT littermates were used as controls. Mice were housed in a quiet individual room in basement with regular 12/12 light/dark cycle. The background noise level in the mouse room at mouse hearing range (4-70 kHz) was <30 dB SPL.

### RNAsample preparation and sequencing

The AC, IC, CN, and cochlear (Fig. 1A&B) were quickly isolated and quick-frozen in liquid nitrogen. RNAs were extracted by use of absolute RNA Miniprep Kit (#400800, Agilent, Santa Clara, CA, United States) following manufacture’s instruction. The extracted RNA was examined for quality and concentration by NanoDrop™ Spectrophotometer (ND-ONE-W, Thermo Fisher Scientific Inc.) and sent to Yale Center for Genome Analysis (YCGA) for Bulk Poly(A) RNA sequencing with NovaSeq paired-End 101 bp (Zhai et al., 2025).

### RNA-Sequencing data analysis

Bulk RNA-sequencing data from AC, IC, CN, and cochlea were analyzed independently. Low-quality terminal sequence was gently trimmed with Btrim (Kong, 2011), reads were aligned to the Mus musculus GRCm39 reference genome with STAR (Dobin et al., 2013), and GRCm39 annotations and identifier mappings were obtained from Ensembl BioMart (Durinck et al., 2009). Programs used default parameters unless otherwise stated. The animal cohort information is listed in Table 1.

**Table 1.** Composition of the 5xFAD auditory system RNA-sequencing cohort.

| Age | WT, n |  | 5xFAD, n |  | Total animals, n | Regional RNA-seq samples, n |
| --- | --- | --- | --- | --- | --- | --- |
|  | Female | Male | Female | Male |  |  |
| **P50** | 3 | 3 | 3 | 3 | 12 | 48 |
| **8 months** | 3 | 3 | 3 | 3 | 12 | 48 |
| **11 months** | 3 | 3 | 3 | 3 | 12 | 48 |
| **16 months** | 4 | 3 | 4 | 5 | 16 | 63 |
| **Total** | **13** | **12** | **13** | **14** | **52** | **207** |

**Note.** Counts in the genotype-by-sex columns represent unique animals. “5xFAD” denotes heterozygous transgenic animals. Each animal was sampled in the auditory cortex (AC), inferior colliculus (IC), cochlear nucleus (CN), and cochlea, except that one 16-month WT female lacked a cochlear sample. Accordingly, the regional totals were AC, $n = 52$ ; IC, $n = 52$ ; CN, $n = 52$ ; and cochlea, $n = 51$ .

Differential expression was analyzed with DESeq2 (Love et al., 2014). The model included biological sex, genotype, categorical age, and the Genotype×Age interaction: design = ∼ Gender + Genotype + Age_TS + Genotype:Age_TS. WT and P50 were reference levels. Wald contrasts estimated the 5xFAD-versus-WT effect within each age and the change in that genotype effect at 8, 11, or 16 months relative to P50. A likelihood-ratio test compared the full model with a reduced model lacking the interaction. Genes with Benjamini–Hochberg-adjusted P < 0.05 were significant.

Genes were ranked by the DESeq2 Wald statistic for clusterProfiler (Xu et al., 2024) GSEA against Hallmark, KEGG, and GO gene sets. Hallmark pathways were significant at BH-adjusted P < 0.05; KEGG and GO pathways were significant at q-value < 0.05. Positive age-specific NES denotes enrichment toward genes increased in 5xFAD relative to WT. Positive interaction NES denotes a positive shift in the genotype effect relative to P50 and does not necessarily indicate positive enrichment at the later age. Cross-region comparisons integrated independently fitted regional interactions descriptively; they were not formal Genotype×Age×Region tests. Detailed contrast construction, enrichment settings, GO redundancy reduction, and figure-selection procedures are provided in Supplementary Methods.

## Results

### P50 transcriptional states across the auditory pathway

#### Auditory cortex

At P50, AC contained 11 differentially expressed genes, including strong induction of *Cst7* and reductions in *Rps3a3* and *Resp18* (Figure 2 (a), 12). GSEA revealed a much broader phenotype than gene-level testing (Figure 2 (B-D), 13). Oxidative phosphorylation was strongly depleted (Hallmark NES = −2.765, adjusted P = 5.62 × 10⁻¹⁷), together with carbon and lipid metabolism, ribosome, spliceosome, proteasome, protein folding, and mTORC1-associated programs. In contrast, axonogenesis, synapse organization, cell adhesion, extracellular-matrix (ECM) interaction, membrane transport, and phosphatidylinositol signaling were positively enriched. These results define an early mismatch between reduced metabolic and proteostatic support and increased structural remodeling. Immune enrichment was selective: *Cst7* increased, but TNF–NF-κB, cytokine activity, and antigen-presentation pathways were depleted.

**Figure 2.**
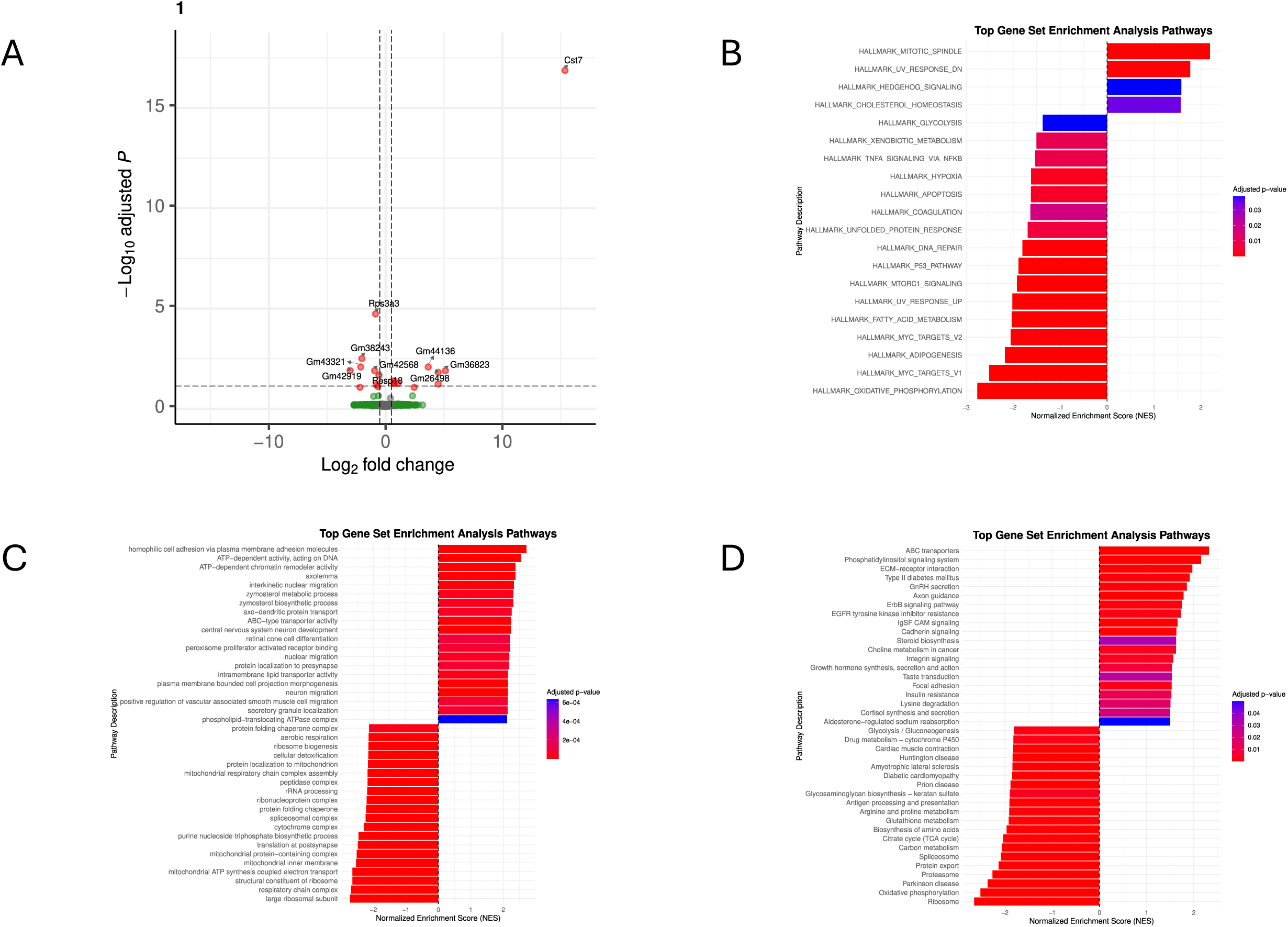
Age-specific transcriptomic alterations in the auditory cortex at P50. (A) Volcano plot showing gene-level differential expression between 5xFAD and wild-type (WT) mice. (B–D) Gene set enrichment analysis (GSEA) of (B) Hallmark, (C) Gene Ontology (GO), and (D) Kyoto Encyclopedia of Genes and Genomes (KEGG) pathways. For each gene-set collection, the 20 most positively enriched and 20 most negatively enriched pathways, ranked by normalized enrichment score (NES), are shown. Positive NES values indicate enrichment in 5xFAD mice, whereas negative NES values indicate enrichment in WT mice.

#### Inferior colliculus

IC showed only two significant genes, increased *Thy1* and decreased *Tmem184c*, but the strongest pathway-level metabolic suppression among the four regions (Figure 3). Hallmark oxidative phosphorylation reached NES = −3.383 (adjusted P = 1.72 × 10⁻³²), accompanied by depletion of the tricarboxylic-acid cycle, glycolysis, fatty-acid metabolism, ribosome, spliceosome, proteasome, myelin-associated genes, and MYC/mTORC1 programs. Synapse organization, neurotransmitter-receptor activity, axon guidance, ECM and adhesion pathways, calcium signaling, and membrane transport were positively enriched. Selected vesicle-trafficking processes remained negative, indicating structural and receptor remodeling rather than uniformly increased neurotransmission. As in AC, immune programs were heterogeneous rather than globally activated.

**Figure 3.**
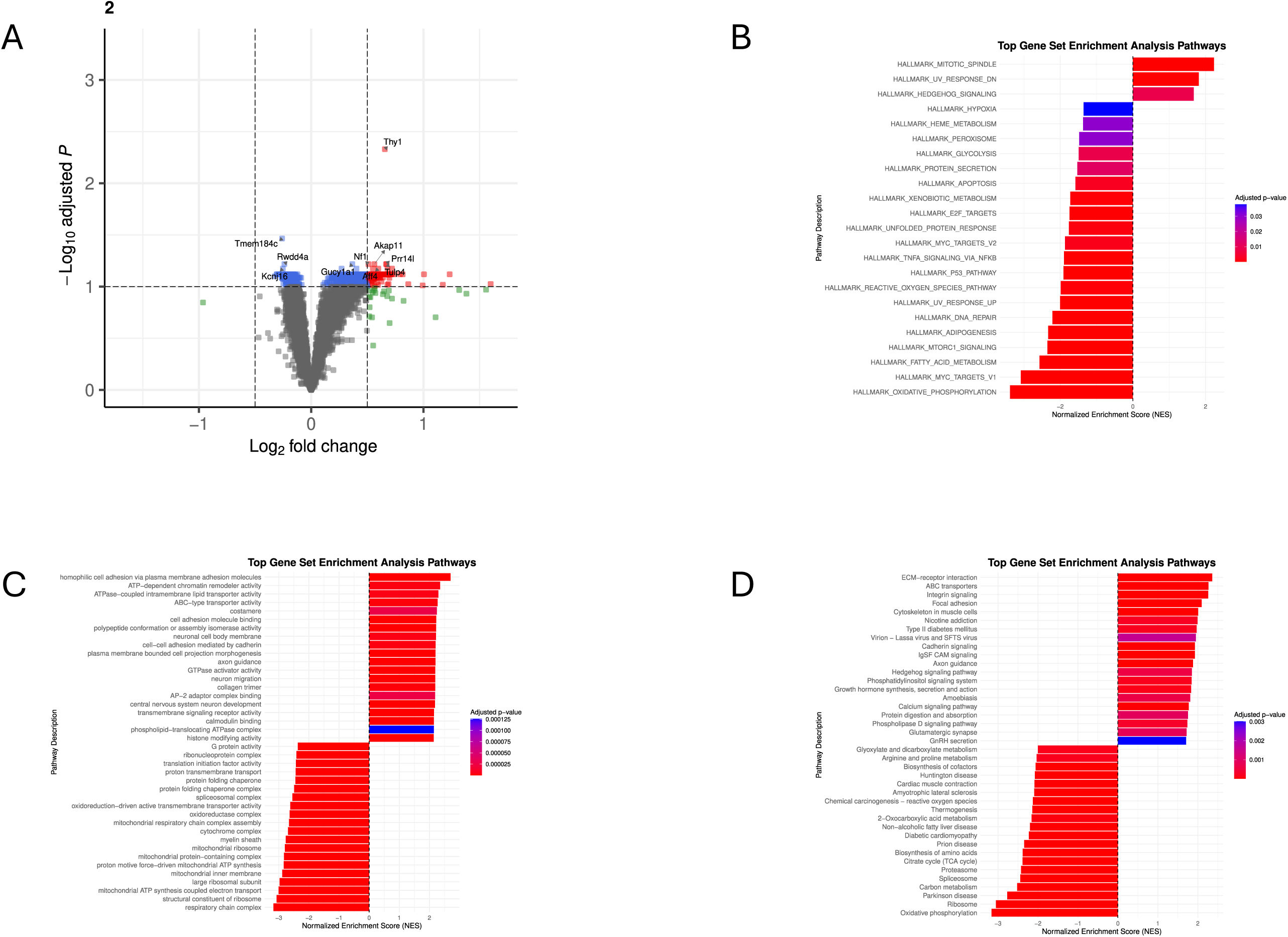
Age-specific transcriptomic alterations in the inferior colliculus at P50. (A) Volcano plot showing gene-level differential expression between 5xFAD and wild-type (WT) mice. (B–D) Gene set enrichment analysis (GSEA) of (B) Hallmark, (C) Gene Ontology (GO), and (D) Kyoto Encyclopedia of Genes and Genomes (KEGG) pathways. For each gene-set collection, the 20 most positively enriched and 20 most negatively enriched pathways, ranked by normalized enrichment score (NES), are shown. NES directionality is defined as in Figure 2.

#### Cochlear nucleus

CN showed two significant genes, increased *Rps3a1* and decreased *Aox4*, and a pathway configuration opposite to AC and IC (Figure 4). Oxidative phosphorylation (Hallmark NES = 2.355, adjusted P = 1.27 × 10⁻¹¹), carbon metabolism, glycolysis, ribosome, spliceosome, proteasome, MYC targets, and mTORC1 signaling were positively enriched. Synaptic transmission, axonal transport, receptor activity, cadherin/integrin signaling, focal adhesion, and inflammatory programs were depleted. The breadth of the positive metabolic and biosynthetic state is consistent with coordinated adaptation or altered cellular composition, although transcript abundance does not establish increased metabolic flux.

**Figure 4.**
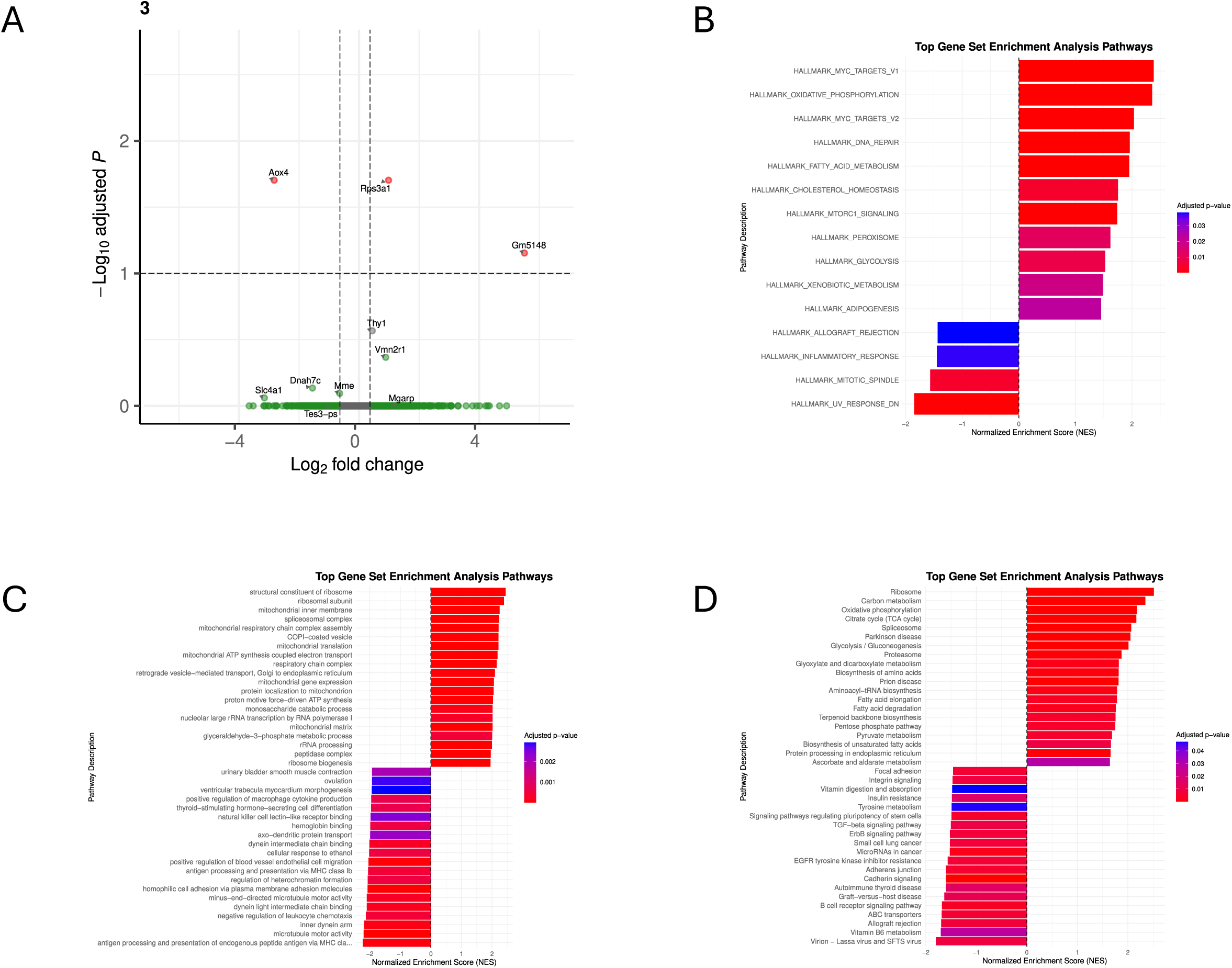
Age-specific transcriptomic alterations in the cochlear nucleus at P50. (A) Volcano plot showing gene-level differential expression between 5xFAD and wild-type (WT) mice. (B–D) Gene set enrichment analysis (GSEA) of (B) Hallmark, (C) Gene Ontology (GO), and (D) Kyoto Encyclopedia of Genes and Genomes (KEGG) pathways. For each gene-set collection, the 20 most positively enriched and 20 most negatively enriched pathways, ranked by normalized enrichment score (NES), are shown. NES directionality is defined as in Figure 2.

#### Cochlea

The P50 cochlea also showed two significant genes, increased *Ighv2-3* and *Gm37261* (Figure 5). Oxidative phosphorylation (Hallmark NES = −2.666, adjusted P = 6.73 × 10⁻¹⁶), carbon metabolism, ribosome, proteasome, synaptic-vesicle cycling, and glutamatergic and GABAergic programs were depleted. ECM, collagen, angiogenesis, cilium movement, stereocilium organization, and hair-cell differentiation were enriched. This combination indicates peripheral sensory and tissue remodeling in an energetically constrained state. The signatures do not establish hair-cell regeneration or increased cell number, and the mixed immune findings do not support broad cochlear inflammation at this age.

**Figure 5.**
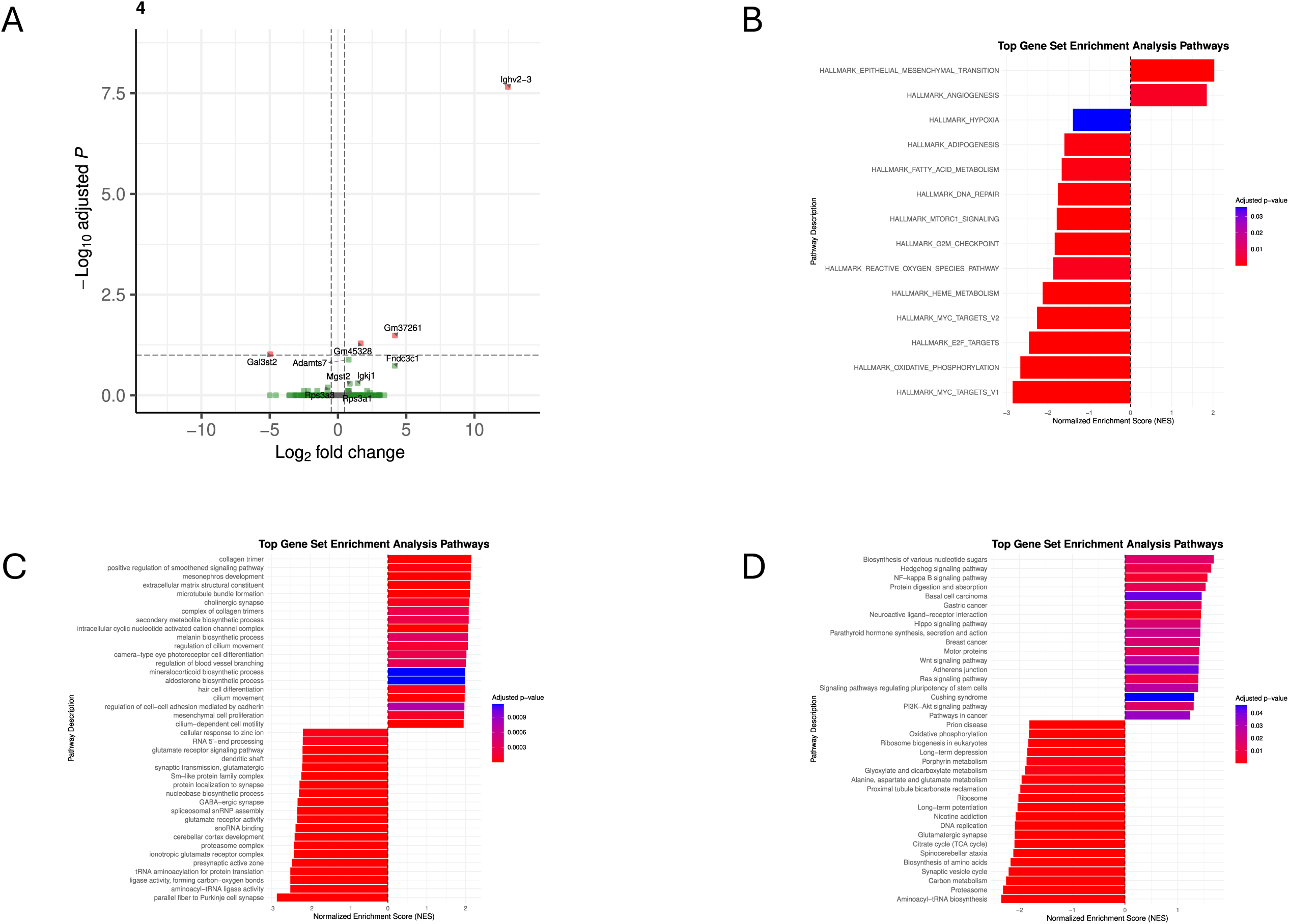
Age-specific transcriptomic alterations in the cochlea at P50. (A) Volcano plot showing gene-level differential expression between 5xFAD and wild-type (WT) mice. (B–D) Gene set enrichment analysis (GSEA) of (B) Hallmark, (C) Gene Ontology (GO), and (D) Kyoto Encyclopedia of Genes and Genomes (KEGG) pathways. For each gene-set collection, the 20 most positively enriched and 20 most negatively enriched pathways, ranked by normalized enrichment score (NES), are shown. NES directionality is defined as in Figure 2.

#### Cross-region synthesis at P50

Cross-region analysis identified two principal axes (Figure 6; Tables S1 and S2). AC, IC, and cochlea shared suppression of mitochondrial metabolism, translation, RNA processing, proteostasis, and myelin-associated genes, whereas CN showed coordinated positive enrichment of the same modules. Oxidative phosphorylation, carbon metabolism, ribosome, and proteasome all followed this three-versus-one configuration. Synaptic organization separated AC and IC from CN and cochlea: regulation of synapse structure or activity was positive in AC and IC but negative in CN and cochlea. Cell-adhesion pathways similarly increased in AC and IC and decreased in CN, while the cochlea showed a distinct ECM, vascular, and sensory-structure response.

**Figure 6.**
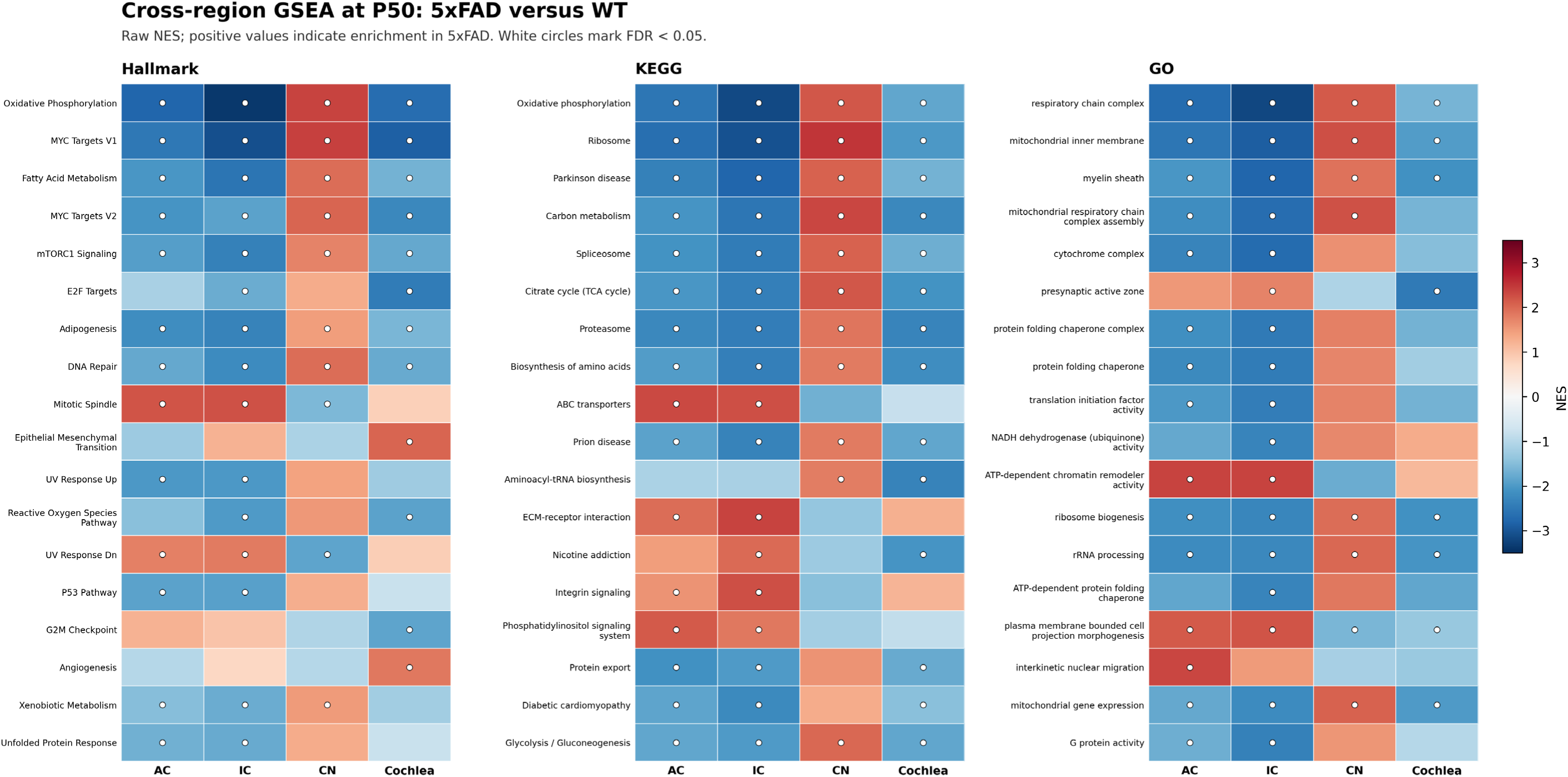
Cross-region gene-set enrichment at P50. Heatmaps show NES for 5xFAD versus WT in AC, IC, CN, and cochlea. Red denotes positive enrichment in 5xFAD and blue denotes negative enrichment; circles mark significant pathways. Main-figure pathways were significant in at least one region and had maximum |NES| ≥ 1.5. Significance was BH-adjusted P < 0.05 for Hallmark and q-value < 0.05 for KEGG and GO. GO terms were retained after independent redundancy reduction in all four regions and balanced across BP, CC, and MF.

These regional relationships argue against a uniform early pathway response. Higher central regions combined metabolic suppression with structural remodeling, CN combined metabolic activation with reduced synaptic and adhesion programs, and cochlea combined metabolic and synaptic suppression with peripheral tissue remodeling. Because regional models were fitted independently, these comparisons describe enrichment-profile differences rather than formal Region×Genotype interactions.

### Age dependent remodeling within each region

#### Auditory cortex

The AC transcriptional phenotype expanded from 11 differentially expressed genes at P50 to 527 at 8 months and 647 at 11 months, then contracted to 277 at 16 months (Figure 11 (a); see also Figure 2, S1, S5, 7, S9-12). By 8 months, interferon responses, complement, IL6–JAK–STAT3, TNF–NF-κB, antigen presentation, phagocytosis, lysosome, and leukocyte pathways were positively enriched. At 11 months, inflammatory and ECM-remodeling signals were broadest, accompanied by positive shifts in ribosomal, proteostatic, and stress-response programs relative to the strongly negative P50 state (Figure 11 (a)). Genes contributing to these responses included *Trem2*, *Itgax*, *Cd68*, *Mpeg1*, *Spp1*, *Tgfb1*, and collagen-associated transcripts.

**Figure 7.**
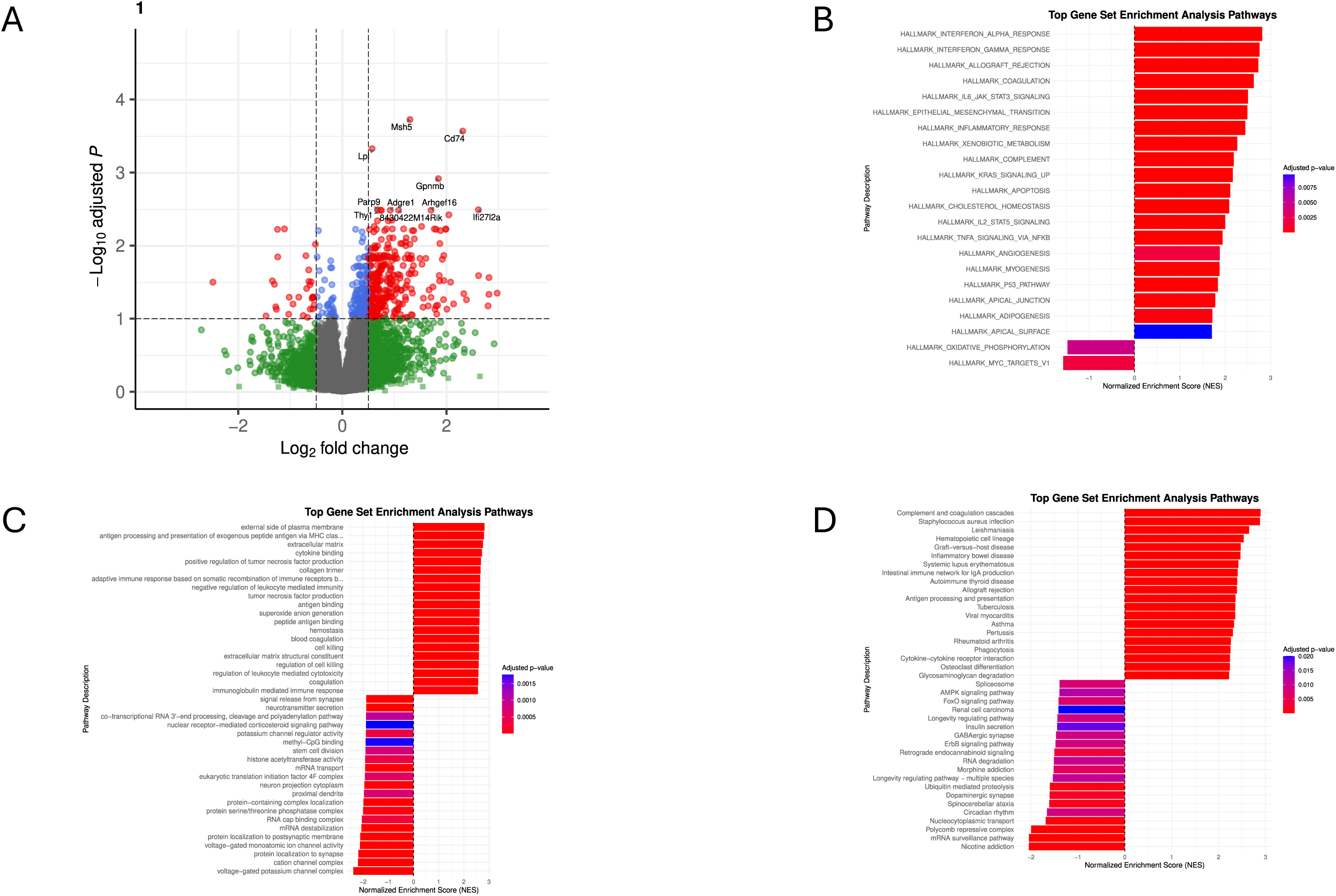
Age-specific transcriptomic alterations in the auditory cortex at 16-month. (A) Volcano plot showing gene-level differential expression between 5xFAD and wild-type (WT) mice. (B–D) Gene set enrichment analysis (GSEA) of (B) Hallmark, (C) Gene Ontology (GO), and (D) Kyoto Encyclopedia of Genes and Genomes (KEGG) pathways. For each gene-set collection, the 20 most positively enriched and 20 most negatively enriched pathways, ranked by normalized enrichment score (NES), are shown. NES directionality is defined as in Figure 2.

Neuronal signaling moved in the opposite direction (Figure 11 (a)). Axon guidance, phosphatidylinositol and ErbB signaling, glutamatergic and cholinergic synapses, synapse assembly, and ion-channel complexes shifted negatively after P50, most clearly at 11 months. At 16 months, inflammatory, phagolysosomal, antigen-presentation, and matrix programs remained enriched, while several metabolic pathways again showed negative age-specific NES. Their positive interaction NES values reflected attenuation of the deeper P50 deficit, illustrating why interaction direction cannot be equated with later-age activation. The omnibus likelihood-ratio test identified 193 genes with significant Genotype×Age effects, whereas only *Rps3a3* and *Cst7* passed the directional interaction threshold at 16 months, consistent with coordinated pathway changes distributed across many genes.

#### Inferior colliculus

IC showed 66 genes with significant Genotype×Age effects and its broadest pathway remodeling at 11 months (Figure 11 (b); see also Figure 3, S2, S6, 8, S13-16). Interferon-α, interferon-γ, IL6–JAK–STAT3, complement, antigen presentation, phagocytosis, and leukocyte-response pathways emerged by 8 months and persisted through 16 months. Age-dependent genes included *Trem2*, *Itgax*, *Tyrobp*, *Clec7a*, *Cd74*, and C1q components, linking the enrichment signal to coordinated myeloid, complement, and antigen-presentation programs.

**Figure 8.**
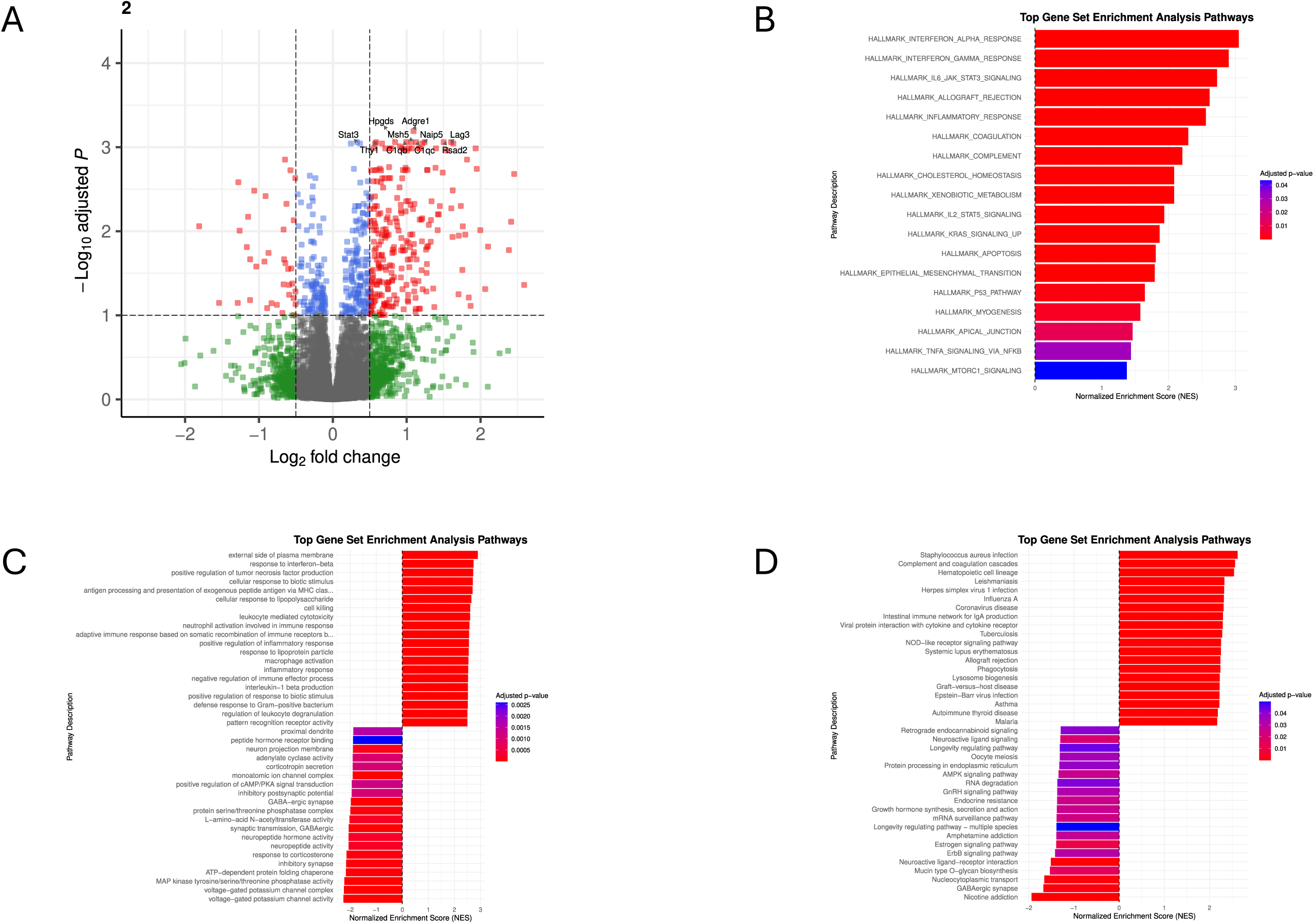
Age-specific transcriptomic alterations in the inferior colliculus at 16-month. (A) Volcano plot showing gene-level differential expression between 5xFAD and wild-type (WT) mice. (B–D) Gene set enrichment analysis (GSEA) of (B) Hallmark, (C) Gene Ontology (GO), and (D) Kyoto Encyclopedia of Genes and Genomes (KEGG) pathways. For each gene-set collection, the 20 most positively enriched and 20 most negatively enriched pathways, ranked by normalized enrichment score (NES), are shown. NES directionality is defined as in Figure 2.

Metabolic and biosynthetic pathways shifted positively relative to their pronounced P50 depletion. Oxidative phosphorylation moved from NES = −3.38 at P50 toward neutral or modestly positive values at later ages; ribosome and proteasome became positively enriched at 11 and 16 months. In contrast, GABAergic and glutamatergic synapses, synapse assembly, axon guidance, calcium signaling, and ciliary or axonemal programs shifted negatively (Figure 11 (b)). ECM pathways were reorganized rather than uniformly increased: coagulation and matrix-degradation programs rose, while ECM–receptor interaction weakened relative to its positive P50 state. Thus, IC progressed from early metabolic suppression with structural remodeling to sustained inflammation and negative synaptic interactions.

#### Cochlear nucleus

CN followed a strongly nonmonotonic trajectory (Figure 11 (c); see also Figure 4, S3, S7, 9, S17-20). At 8 months, its significant Hallmark interactions were uniformly positive and concentrated in interferon responses, complement, coagulation, IL6–JAK–STAT3, and inflammatory programs. The 11-month profile reversed: interferon, ECM-remodeling, oxidative-phosphorylation, and myogenesis interactions were negative, whereas neuroactive-ligand and several synaptic-organization programs shifted positively.

**Figure 9.**
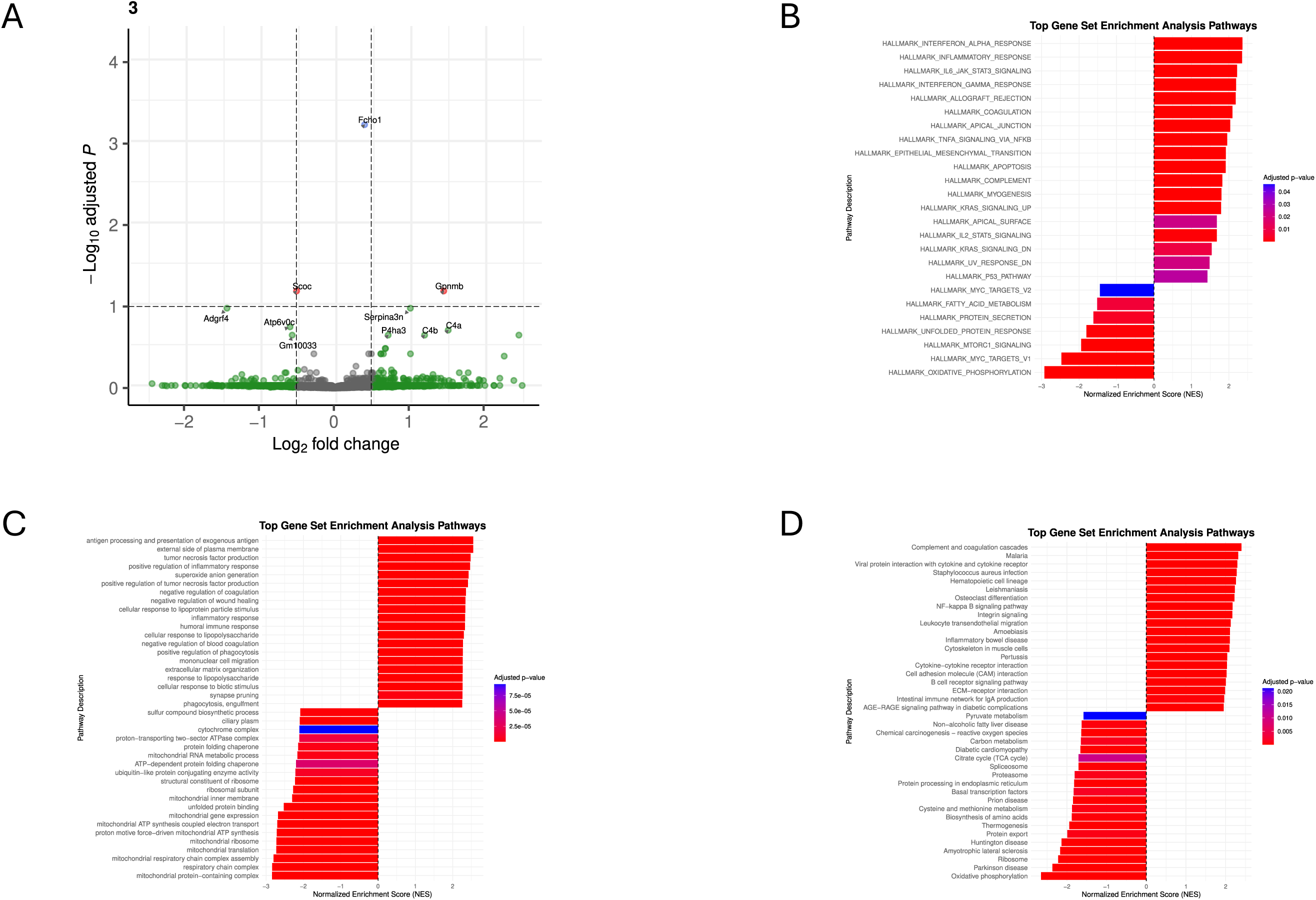
Age-specific transcriptomic alterations in the cochlear nucleus at 16-month. (A) Volcano plot showing gene-level differential expression between 5xFAD and wild-type (WT) mice. (B–D) Gene set enrichment analysis (GSEA) of (B) Hallmark, (C) Gene Ontology (GO), and (D) Kyoto Encyclopedia of Genes and Genomes (KEGG) pathways. For each gene-set collection, the 20 most positively enriched and 20 most negatively enriched pathways, ranked by normalized enrichment score (NES), are shown. NES directionality is defined as in Figure 2.

By 16 months, CN showed its broadest departure from P50. Immune, antigen-presentation, phagocytic, adhesion, TGF-β, and ECM programs shifted positively, while oxidative phosphorylation had the strongest negative Hallmark interaction (NES = −3.17). MYC targets, mTORC1, fatty-acid metabolism, ribosome, spliceosome, proteasome, and protein processing also shifted negatively, reversing the positive metabolic state observed at P50. Synaptic responses remained mixed. Only three genes, *Gm56815*, *Fcho1*, and *Scoc*, passed the omnibus Genotype×Age threshold, and none passed the directional interaction threshold, indicating that the CN trajectory was distributed across coordinated gene sets.

#### Cochlea

Cochlear remodeling was also nonmonotonic (Figure 11 (d); see also Figure 5, S4, S8, 10, S21-24). At 8 months, ECM, cytoskeletal, adhesion, myogenesis, complement, and coagulation programs strengthened relative to P50. At 11 months, inflammatory and matrix pathways shifted negatively, while glutamatergic, GABAergic, presynaptic, and neuroactive-ligand programs changed from negative enrichment at P50 or 8 months to positive enrichment. Cilium-movement and axonemal programs shifted strongly negative, separating the transient synaptic state from microtubule-based structural processes.

**Figure 10.**
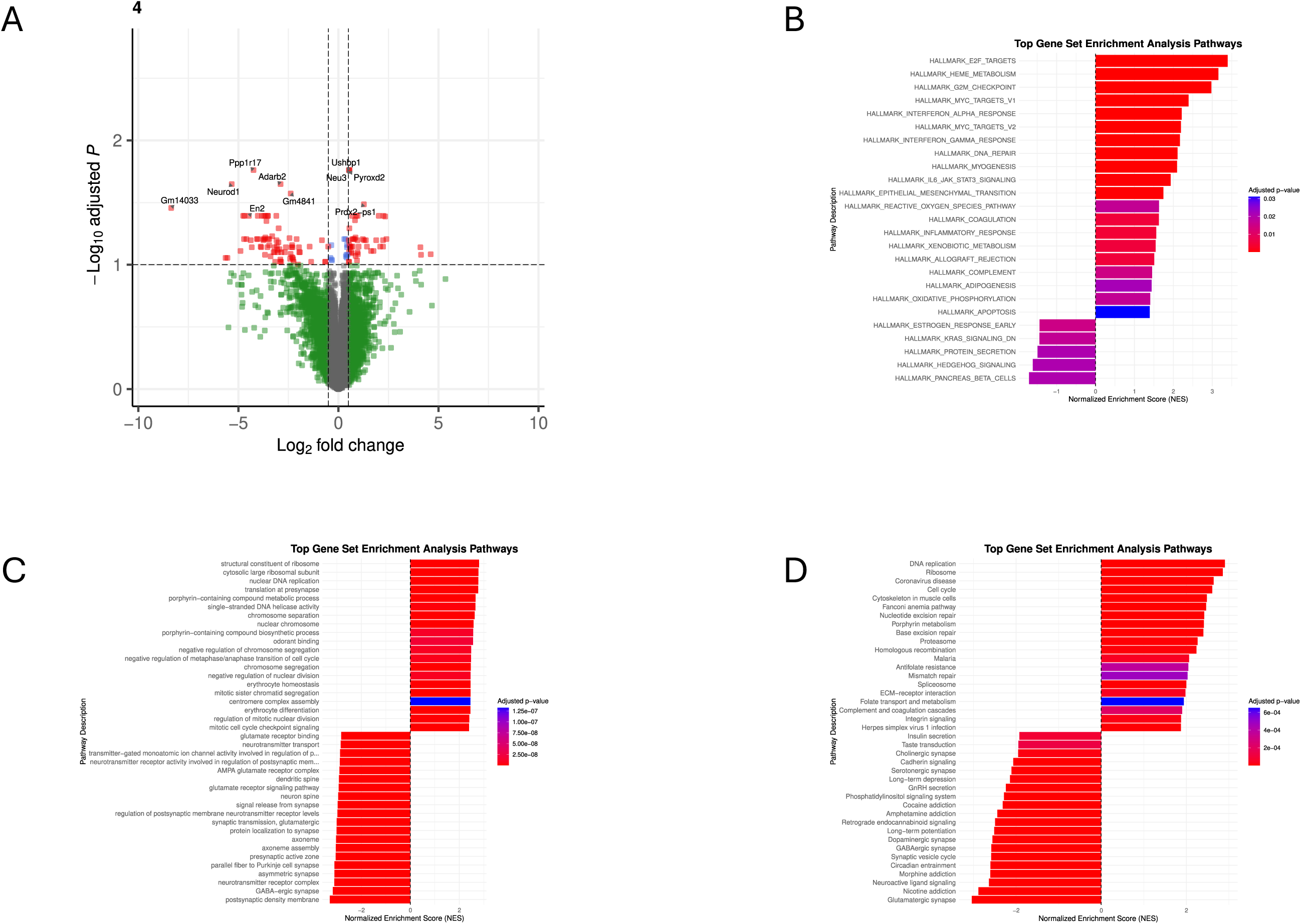
Age-specific transcriptomic alterations in the cochlea at 16-month. (A) Volcano plot showing gene-level differential expression between 5xFAD and wild-type (WT) mice. (B–D) Gene set enrichment analysis (GSEA) of (B) Hallmark, (C) Gene Ontology (GO), and (D) Kyoto Encyclopedia of Genes and Genomes (KEGG) pathways. For each gene-set collection, the 20 most positively enriched and 20 most negatively enriched pathways, ranked by normalized enrichment score (NES), are shown. NES directionality is defined as in Figure 2.

**Figure 11.**
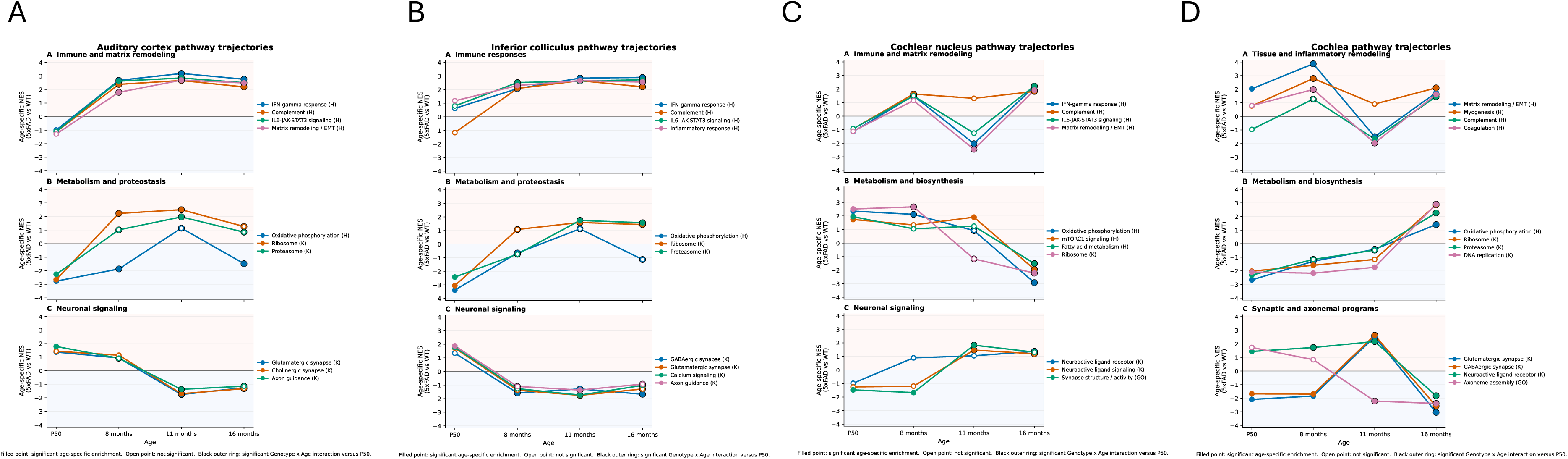
Representative age-dependent pathway trajectories in the four auditory regions. (A) Auditory cortex. Representative age-dependent pathway trajectories in the auditory cortex of 5xFAD mice. Lines show age-specific normalized enrichment scores (NES) for 5xFAD versus WT mice at P50, 8 months, 11 months, and 16 months. Representative pathways were selected from the statistically supported processes emphasized in the auditory-cortex Results and organized into (A) immune and matrix remodeling, (B) metabolism and proteostasis, and (C) neuronal signaling. Positive and negative NES indicate enrichment toward genes increased and decreased, respectively, in 5xFAD relat ive to WT mice. Filled points mark significant age-specific enrichment, whereas open points indicate nonsignificant enrichment. Black outer rings at 8, 11, or 16 months mark significant Genotyp e x Age interactions relative to P50. Significance was defined as BH-adjusted P < 0.05 for Hallmark pathways and q-value < 0.05 for KEGG pathways. H and K in pathway labels denote Hallmark and KEGG, respectively. Interaction significance indicates that the genotype effect changed relative to P50; it does not independently specify the direction of enrichment at the later age. (B) Inferior colliculus. Representative pathways were selected from the statistically supported processes emphasized in the inferior-colliculus Results and organized into (A) immune responses, (B) metabolism and proteostasis, and (C) neuronal signaling. (C) Cochlear nucleus. Representative pathways were selected from the statistically supported processes emphasized in the cochlear-nucleus Results and organized into (A) immune and matrix remodeling, (B) metabolism and biosynthesis, and (C) neuronal signaling. (D) Cochlea. Representative pathways were selected from the statistically supported processes emphasized in the cochlear Results and organized into (A) tissue and inflammatory remodeling, (B) metabolism and biosynthesis, and (C) synaptic and axonemal programs.

**Figure 12.**
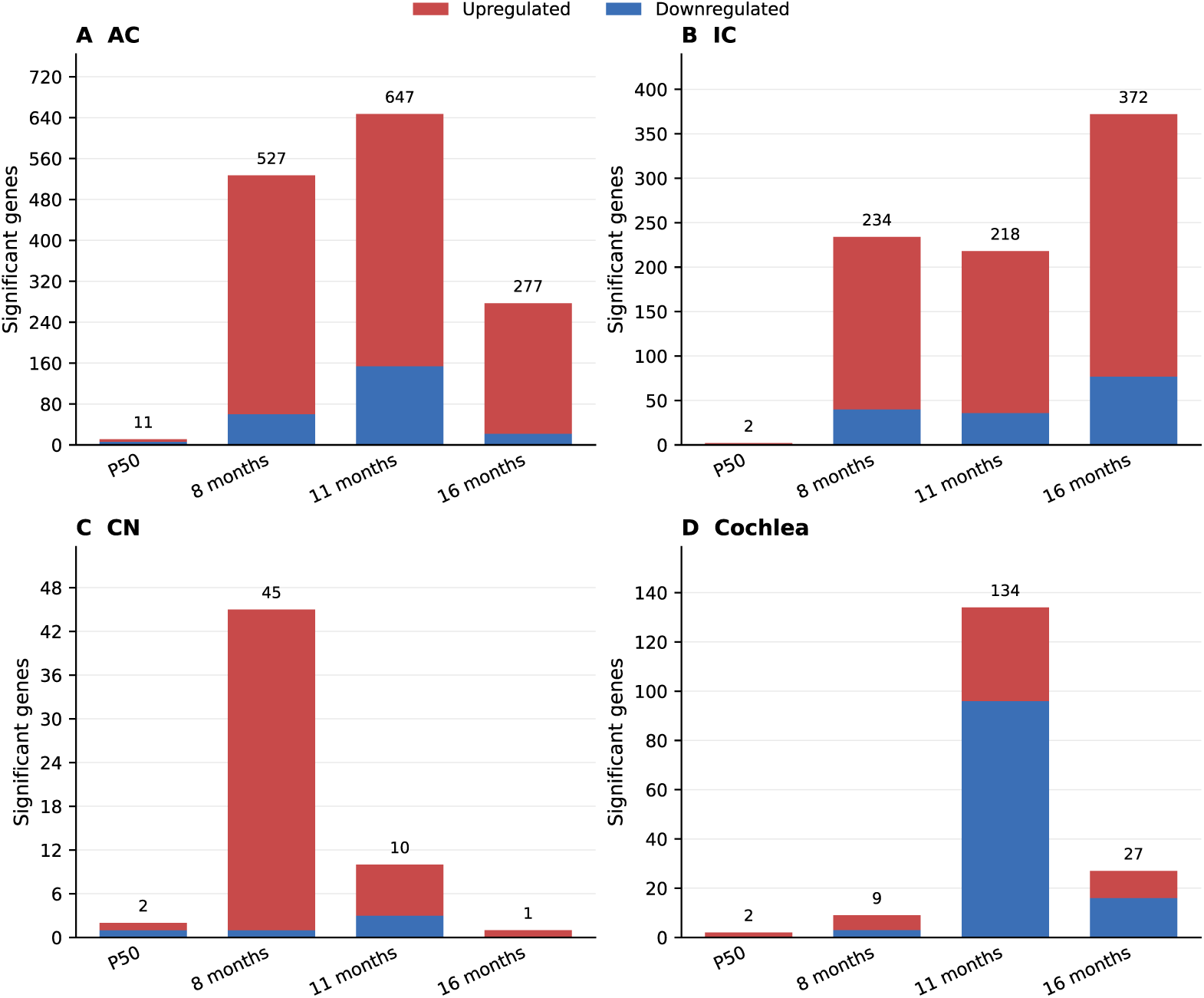
Age-dependent numbers of differentially expressed genes across the auditory pathway. Stacked bars show the numbers of genes that differed significantly between 5xFAD and WT mice within each age group in the auditory cortex (AC), inferior colliculus (IC), cochlear nucleus (CN), and cochlea. Differentially expressed genes were defined by a DESeq2 BH-adjusted P < 0.05. Red and blue segm ents indicate genes with positive and negative log2 fold changes, respectively, in 5xFAD relative to WT mice. Numbers above bars indicate total significant genes. The y-axis is scaled independently fo r each region to display regional temporal patterns clearly; therefore, bar heights should not be compared across panels without reference to the displayed counts. No fold-change threshold was applied.

**Figure 13.**
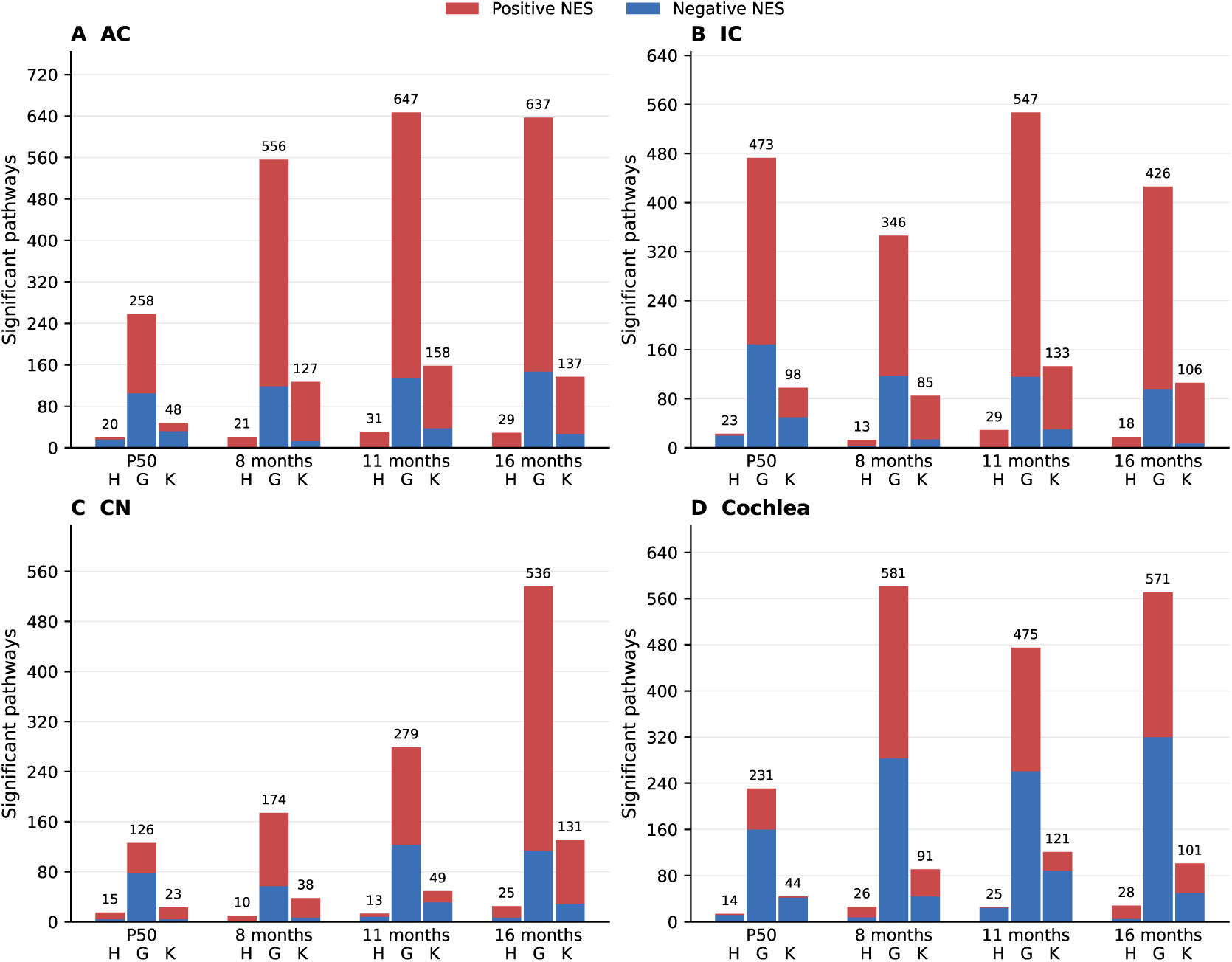
Age-dependent numbers of significantly enriched pathways across the auditory system. Stacked bars show the numbers of significant Hallmark, KEGG, and Gene Ontology (GO) gene sets in age-specific gene set enrichment analyses comparing 5xFAD with WT mice in the auditory cortex (AC), inferior colliculus (IC), cochlear nucleus (CN), and cochlea. Red and blue segments indicate positive and negative normalized enrichment scores (NES), respectively, in 5xFAD relative to WT mice. Numbers above bars indicate total significant pathways. Significance was defined as BH-adjusted P < 0.05 for Hallmark and q-value < 0.05 for KEGG and GO. A common y-axis scale is used across regions within each pathway database, whereas scales differ among Hallmark, KEGG, and GO because their tested gene-set universes differ substantially. GO counts reflect the independently redundancy-reduced GO results for each region and age and therefore summarize the retained terms rather than a fixed common GO universe.

At 16 months, ribosome, DNA replication, cell-cycle, proteasome, heme-metabolism, and oxidative-phosphorylation programs shifted strongly positive relative to P50, whereas synaptic pathways again became negative. Glutamatergic synapse changed from NES = 2.52 at 11 months to −3.04 at 16 months, and ciliary/axonemal suppression persisted. The likelihood-ratio test identified 15 genes with significant Genotype×Age effects, but only one, four, and one genes passed directional interaction testing at 8, 11, and 16 months. The late biosynthetic activation and transient midlife synaptic enrichment therefore reflect coordinated pathway-level changes rather than widespread large single-gene effects.

### Cross-region temporal synthesis

We integrated the 12 regional age contrasts to distinguish conserved from region-dependent trajectories (Figure 14; S25, Table S4). At 8 months, eight Hallmark pathways showed significant positive interactions in all four regions: allograft rejection, apoptosis, coagulation, complement, IL6–JAK–STAT3 signaling, inflammatory response, interferon-γ response, and oxidative phosphorylation. Fifteen Hallmark and 34 KEGG pathways were shared by at least three regions, and pairwise interaction-profile correlations were positive. These data identify a coordinated early shift in inflammatory, cell-survival, and bioenergetic programs throughout the auditory system.

**Figure 14.**
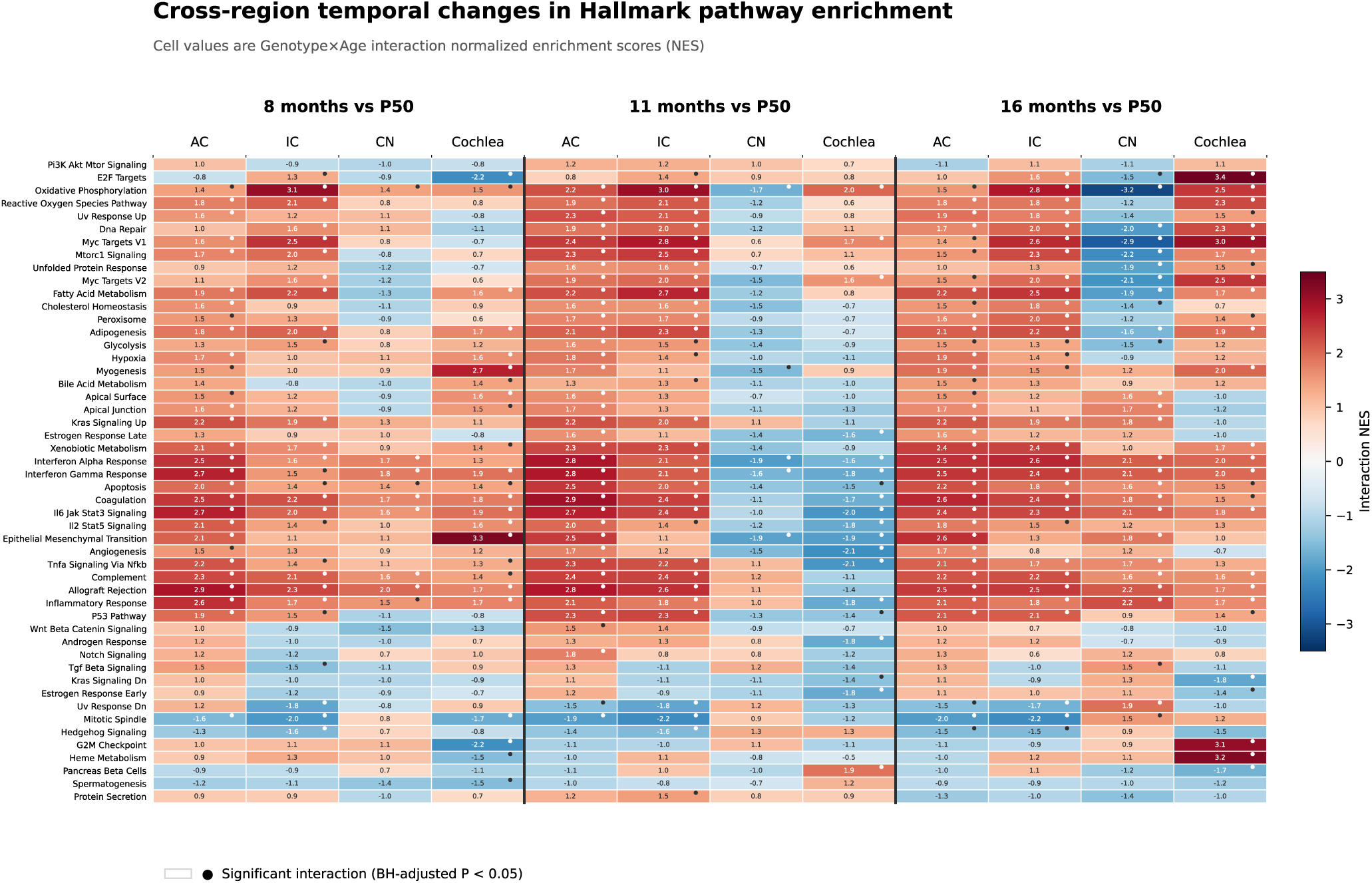
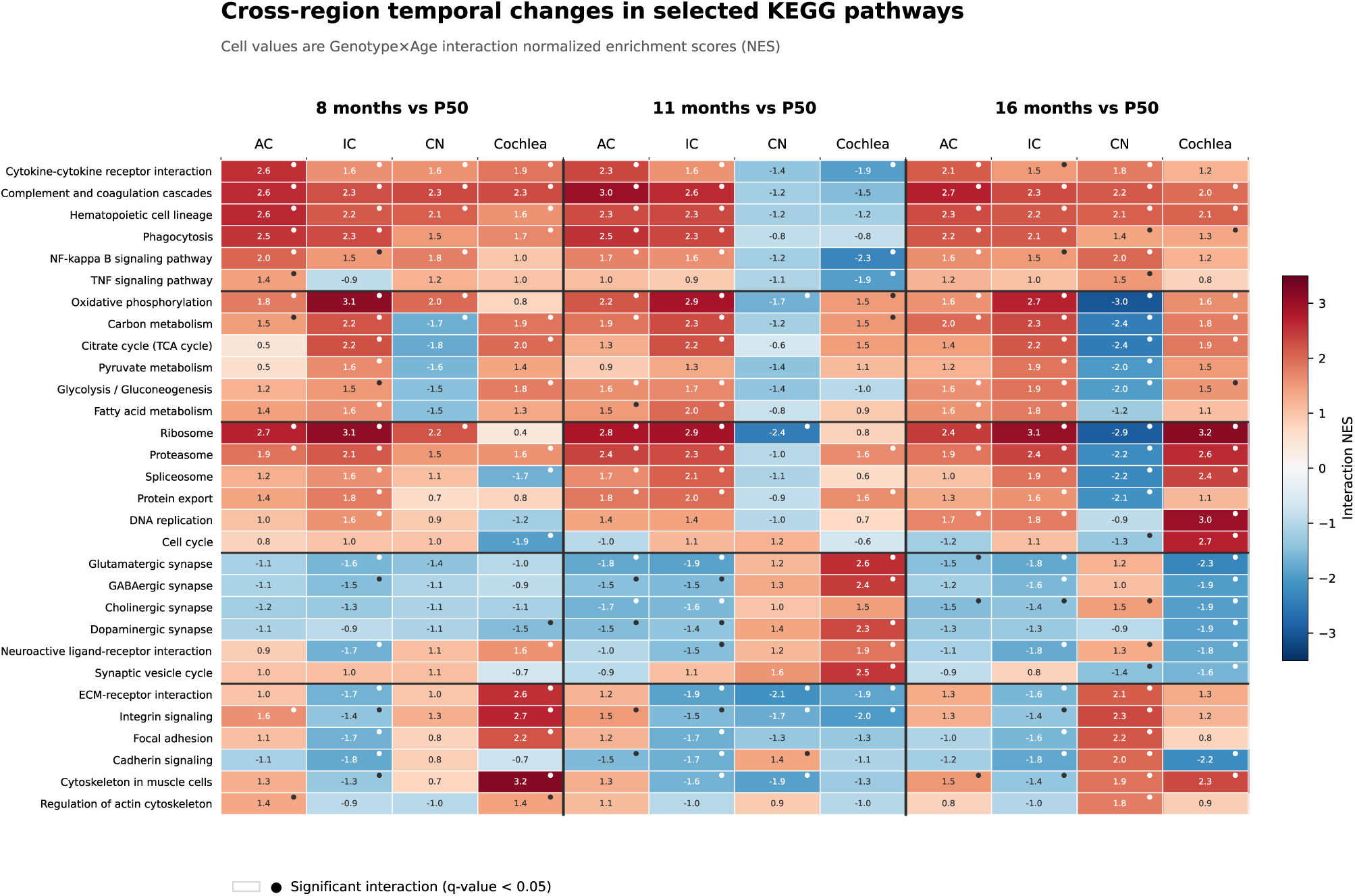
Cross-region temporal changes in pathway enrichment. Heatmaps show Genotype×Age interaction NES across AC, IC, CN, and cochlea at 8, 11, and 16 months relative to P50. (A) All 50 shared Hallmark pathways; (B) 30 prespecified KEGG pathways across the principal biological domains. Red and blue denote positive and negative shifts; dots mark BH-adjusted P < 0.05 for Hallmark or q-value < 0.05 for KEGG. Interaction direction describes change from P50, not absolute later-age enrichment.

Coordination fractured at 11 months. No Hallmark or KEGG pathway met the four-region conserved criterion, whereas 14 Hallmark and 62 KEGG pathways showed significant opposite-direction interactions. AC and IC remained closely correlated, but each was inversely related to CN. Interferon-γ shifted positively in AC and IC and negatively in CN and cochlea. Oxidative phosphorylation shifted positively in AC, IC, and cochlea but negatively in CN. Synaptic terms also divided anatomically: glutamatergic and GABAergic programs shifted negatively in AC and IC but positively in lower or peripheral regions.

At 16 months, immune interactions converged again. Eight Hallmark and 12 KEGG pathways showed conserved positive interactions, and the broadly shared set expanded to 23 Hallmark and 71 KEGG pathways. Metabolic, biosynthetic, and synaptic programs remained region dependent. Oxidative phosphorylation and ribosome shifted positively in AC, IC, and cochlea and negatively in CN; regulation of synapse structure or activity showed the opposite configuration. The temporal pattern therefore comprises early coordination, midlife regional divergence, and late inflammatory convergence superimposed on persistent local metabolic and neuronal states.

## Discussion

### Principal findings and central synthesis

By evaluating four anatomically distinct auditory structures across four sequential time points that capture early molecular or peri-plaque change, established plaque-associated disease, advanced synaptic pathology, and late neurodegeneration, this study provides a comprehensive spatiotemporal map of auditory-system involvement in 5xFAD mice. The response was distributed but asynchronous. At P50, AC, IC, and cochlea shared suppression of mitochondrial, biosynthetic, and proteostatic programs, while CN displayed the opposite metabolic state. At 8 months, inflammatory interactions converged across the pathway. Regional coordination then fractured at 11 months, particularly between AC/IC and CN/cochlea, before immune pathways converged again at 16 months. Metabolic and synaptic trajectories remained anatomically differentiated throughout. This organization does not support a monotonic pathological wave from cochlea to cortex or cortex to cochlea; it instead indicates stage-dependent coupling and decoupling among connected auditory compartments.

Pathway-level changes were extensive even where few individual genes passed genome-wide correction, indicating that the auditory phenotype is distributed across coordinated networks of moderate effects. The major cross-region relationships were also nonmonotonic. Inflammation followed a broadly shared trajectory, whereas energy production, protein homeostasis, and neuronal communication followed local trajectories.

### Chronological pathology in the 5xFAD model

P50 lies near an important transition. Intraneuronal Aβ42 begins around 1.5 months, while extracellular deposition and gliosis emerge near 2 months (Oakley et al., 2006). In the auditory system, intracellular Aβ42 is present in primary auditory cortical neurons by P21, whereas plaques become apparent later in AC and IC and remain limited in early lower-brainstem structures (Weible and Wehr, 2022). P50 is therefore early or peri-plaque for the auditory pathway, but it does not precede all Aβ-associated stress. Suppression of oxidative phosphorylation, translation, RNA processing, and proteostasis in AC, IC, and cochlea may reflect soluble or intraneuronal Aβ, altered activity, or transgene-associated cellular stress; RNA sequencing cannot distinguish these triggers or establish reduced metabolic flux.

By 8 months, the conserved interferon, complement, cytokine, phagocytic, and lysosomal interactions resembled the plaque-associated inflammatory phase described in 5xFAD cortex and hippocampus. *Trem2*-, *Itgax*-, *Tyrobp*-, *Clec7a*-, *Cst7*-, and antigen-presentation-associated signals in central regions are consistent with disease-associated myeloid programs reported in 5xFAD brain (Keren-Shaul et al., 2017; Zhou et al., 2020), although bulk tissue cannot separate activation from altered cell abundance.

The 11-month interval represented reorganization rather than simple amplification. AC and IC retained inflammation and acquired negative synaptic interactions, whereas CN and cochlea showed relative reductions in interferon-associated enrichment and positive shifts in selected synaptic programs. The cochlear synaptic reversal was transient and should not be interpreted as restored function. At 16 months, renewed inflammatory convergence was consistent with advanced amyloid-associated disease, yet CN remained metabolically distinct. The auditory system therefore mirrors the canonical progression from early cellular stress to plaque-associated inflammation and synaptic disturbance while adding prominent peripheral involvement and a nonmonotonic first-relay response.

### Spatial dynamics across the auditory pathway

The regional pattern does not support a strictly bottom-up sequence. Metabolic suppression was already prominent in AC and strongest in IC at P50, while CN showed positive enrichment of the same programs. This pattern agrees with reports that auditory Aβ pathology appears first in cortex and later in IC, with limited early CN plaque burden (Weible and Wehr, 2022). Positive enrichment of synaptic structure and axon-guidance programs in AC and IC may reflect remodeling under early energetic stress rather than enhanced transmission.

A purely top-down model is also insufficient because the P50 cochlea showed metabolic and synaptic suppression together with vascular, ECM, ciliary, and sensory-structure remodeling. CN did not occupy a simple intermediate state: its early metabolic activation accompanied negative synaptic and adhesion programs and later reversed. CN may initially increase cellular support in response to altered afferent input or network demand and later lose that capacity as inflammation expands, but cell-resolved measurements are required to test this hypothesis.

The 11-month split is best described as higher-central versus lower/peripheral decoupling. Extensive parallel projections and corticofugal feedback allow local pathology to alter distant activity in both directions. Early cortical dysfunction could change descending control of IC and CN, while cochlear or auditory-nerve dysfunction could drive central homeostatic gain. The increased central gain reported before later reductions in auditory-nerve output in 5xFAD mice supports this network-level interpretation (Na et al., 2023), although the present data do not establish causal direction.

### Mechanistic implications

The earliest shared mechanism involved the energetic and proteostatic systems needed for rapid auditory signaling. Coordinated depletion of oxidative phosphorylation, the tricarboxylic-acid cycle, mitochondrial translation, ribosome biogenesis, chaperone activity, and proteasomal function would reduce the transcriptional reserve supporting ion-gradient restoration, axonal transport, vesicle cycling, and protein quality control. In AC and IC, this state coincided with structural and synaptic-organization enrichment, suggesting that remodeling demand arose in cells with reduced molecular support. Layer 5 synaptic deficits precede overt structural decay in 5xFAD cortex, providing precedent for early physiological dysfunction (Buskila et al., 2013), but the present data do not directly measure ATP production or protein turnover.

Neuroimmune activation dominated after P50. Interferon, complement, antigen presentation, phagocytosis, lysosome, and cytokine pathways are consistent with engagement of microglia, astrocytes, and vascular-associated or infiltrating cells. *Comp*lement-dependent microglial engulfment can mediate synapse loss in amyloid models (Hong et al., 2016), providing a plausible link between C1q-associated expression, phagocytic programs, and declining synaptic gene sets. This mechanism remains inferential because complement deposition and synaptic engulfment were not measured. Disease- or infection-labeled KEGG pathways should likewise be interpreted as shared immune-gene modules rather than evidence of the named conditions.

Neuronal changes were related to but not reducible to inflammation. Early structural enrichment in AC and IC gave way to negative glutamatergic, GABAergic, calcium-signaling, synapse-assembly, and spine-related interactions. These signals may reflect synapse loss, altered excitation–inhibition balance, reduced neuronal representation, or transcriptional regulation within surviving neurons. The transient positive synaptic state in CN and cochlea suggests that compensation may occur at different ages in different compartments.

ECM, adhesion, vascular, and cytoskeletal pathways define an additional response axis. Their enrichment may arise from gliosis, perivascular remodeling, altered blood–tissue interfaces, or changes in cellular composition. The Hallmark epithelial–mesenchymal-transition set is therefore best interpreted here as tissue and matrix remodeling. E2F and G2M enrichment in aged cochlea need not indicate proliferation of postmitotic hair cells, and cochlear stereocilia are actin-based rather than motile cilia or microtubule axonemes. Spatial localization is required before assigning these gene sets to specific cochlear cell types.

### Implications for auditory biomarkers in Alzheimer disease

The early molecular phenotype strengthens the view that auditory dysfunction can accompany AD-related biology before extensive plaque deposition or neuronal loss. Gap-detection and cortical encoding deficits occur in 5xFAD mice at a similar age (Kaylegian et al., 2019; Weible et al., 2020). Because early changes occurred in both central and peripheral compartments and differed in direction, hearing should not be treated as a unitary phenotype. Threshold sensitivity, auditory-nerve output, brainstem gain, cortical temporal processing, and sensory gating may report distinct and temporally separable mechanisms.

Human longitudinal studies link central auditory processing and speech-in-noise impairment to subsequent dementia (Gates et al., 2002, 2011; Stevenson et al., 2022). Auditory tests are inexpensive and repeatable, but they are neither causal readouts nor AD-specific biomarkers. The present map identifies a rational measurement framework: gap detection and cortical temporal coding may probe early central dysfunction; ABR wave I and later-wave relationships may capture peripheral-to-central gain; and speech-in-noise performance may reflect distributed network stress. Prospective studies must determine whether these measures track amyloid, tau, cognition, and disease progression beyond age-related peripheral hearing loss.

### Links to previous electrophysiological findings and music-based interventions

The present molecular findings extend our previous electrophysiological evidence of early auditory dysfunction in AD mouse models. Using auditory brainstem response (ABR) recordings, we demonstrated that APP/PS1 mice developed high-frequency hearing loss as early as 2 months of age, before the reported onset of spatial learning deficits (Liu et al., 2020). We subsequently identified reduced early sensory auditory-evoked cortical potentials (AECPs) and a diminished later P3 component in APP/PS1 mice, with the latter differing from the pattern observed during normal aging (Mei et al., 2021). Consistent with our previous findings of early functional changes in the auditory system, the current transcriptomic analysis provides a complementary molecular framework: early disturbances in mitochondrial, biosynthetic, and proteostatic programs could compromise the cellular support required for synchronized auditory responses, whereas later immune activation and synaptic alterations could contribute to progressive electrophysiological dysfunction. Together, these studies identified abnormalities at both subcortical and cortical levels at the early AD stages.

These findings also motivate investigation of auditory stimulation as a potential intervention for Alzheimer disease and related dementias (AD/ADRD). A randomized study reported improvements in mood and selected cognitive domains following regular singing or music listening in people with dementia (Särkämö et al., 2014), although another multicenter trial found no significant additional benefit of individualized music listening or active music therapy over standard care for behavioral and psychological symptoms (Raglio et al., 2015).

Recent studies also have shown that gamma-frequency sensory stimulation, including auditory and visual stimulation, can reduce brain amyloid burden and improve memory in AD mouse models (Iaccarino et al., 2016; Adaikkan et al., 2019; Martorell et al., 2019; Wang et al., 2026). These results suggest that sensory-driven neural activity can influence AD-related pathology and cognitive outcomes, supporting a role for auditory function beyond disease detection. Our spatiotemporal molecular map raises the possibility that responses to auditory interventions depend on disease stage and the functional integrity of individual auditory compartments. Early metabolic and proteostatic disturbances may limit the capacity to sustain stimulus-driven activity, whereas later inflammatory and synaptic changes may alter neural synchronization and network responsiveness. Combining controlled auditory stimulation with ABR, AECP, cognitive testing, and molecular analyses could determine whether functional improvement accompanies changes in these disease-associated trajectories. Music listening offers an additional avenue for auditory engagement, although its effects should be evaluated separately from those of precisely timed gamma-frequency stimulation. Together, these approaches could help establish when and how auditory interventions influence AD/ADRD progression.

## Acknowledgement

This work was supported by NIH Grants R01 AG080587 and R01/RF1 AG082216 to HBZ.

## Author contributions

HBZ and YK designed experiments. YK, TYZ, and HBZ performed experiments. YK and HBZ analyzed data and wrote paper. YK, TYZ, and HBZ reviewed the paper.

## Data and materials availability

Data and materials availability.The raw RNA-sequencing data and processed gene-level count matrix generated in this study are publicly available through the NCBI Gene Expression Omnibus (GEO) under accession number GSE351085. All other data supporting the findings of this study are provided in the article and its Supplementary Information. Analysis code and additional materials are available from the corresponding author upon reasonable request.

## Supplementary information

### Supplementary Methods

#### Differential expression and interaction models

AC, IC, CN, and cochlea were analyzed independently in DESeq2. WT and P50 were the reference levels for genotype and age. Biological sex, stored as Gender in the sample metadata, was included as a covariate. The full design was:

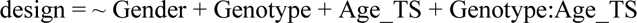

The main Genotype coefficient represented 5xFAD versus WT at P50. The genotype effect at each later age combined the main Genotype coefficient with the corresponding interaction coefficient. Each interaction represented (5xFAD − WT at the later age) − (5xFAD − WT at P50). Positive and negative coefficients therefore indicate positive and negative temporal shifts relative to P50; neither direction alone establishes the sign of differential expression at the later age.

An omnibus likelihood-ratio test identified genes whose genotype effect varied with age. The full model was compared with the reduced model:

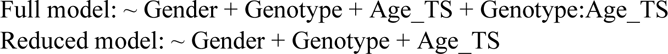

The likelihood-ratio test was fitted using dds_lrt <-DESeq(dds, test = “LRT”, reduced = ∼ Gender + Genotype + Age_TS). Wald tests estimated age-specific genotype contrasts and each later-age-versus-P50 interaction. DESeq2 normalization, dispersion estimation, model fitting, independent filtering, and other settings used defaults. Benjamini–Hochberg-adjusted P < 0.05 defined gene-level significance; genes without valid inferential statistics were treated as untested.

#### Gene set enrichment analysis

Genes were ranked by the DESeq2 Wald statistic for each age-specific genotype comparison and interaction contrast; genes without finite statistics were excluded. clusterProfiler analyzed Hallmark gene sets with GSEA, KEGG pathways with gseKEGG for Mus musculus, and GO biological process, cellular component, and molecular function terms with gseGO. Hallmark pathways were significant at BH-adjusted P < 0.05. KEGG and GO pathways were significant at q-value < 0.05.

Positive age-specific NES denotes enrichment toward genes increased in 5xFAD relative to WT, and negative NES denotes enrichment toward genes decreased in 5xFAD. For interactions, positive and negative NES denote positive and negative shifts in the genotype effect relative to P50. GO redundancy was reduced separately within every region and contrast using clusterProfiler “`simplify”` with default parameters; cross-analysis comparisons therefore used exact GO identifiers.

#### Regional heatmaps

Regional heatmaps displayed age-specific NES at P50, 8, 11, and 16 months. Dots marked significant age-specific enrichment, and black cell outlines marked significant Genotype×Age interactions. Hallmark panels included every pathway with a significant interaction at one or more later ages and age-specific results at all four ages. Each pathway was assigned to a positive- or negative-shift group according to its strongest significant interaction NES.

For KEGG and GO, eligible pathways had a significant interaction in at least one later-age contrast and age-specific results at all four ages. The 20 strongest predominantly positive and 20 strongest predominantly negative interaction shifts were selected. Positive and negative scores were the largest positive interaction NES and largest absolute negative interaction NES, respectively, across significant later-age contrasts.

#### Cross-region synthesis

The cross-region analysis integrated the 8-, 11-, and 16-month interactions relative to P50, producing 12 region-by-age profiles. Common universes contained 50 Hallmark pathways, 364 KEGG pathways, and 1,109 exact GO identifiers. A conserved pathway showed a significant same-direction interaction in all four regions at a given age. A broadly shared pathway showed a significant same-direction interaction in at least three regions and included conserved pathways. A divergent pathway showed at least one significant positive and one significant negative regional interaction.

The Hallmark heatmap contained all 50 shared pathways and used average-linkage hierarchical clustering of Euclidean distances after row standardization across the 12 interactions. The KEGG panel contained 30 prespecified pathways in immune/inflammatory, metabolic/mitochondrial, proteostatic/biosynthetic, synaptic, and ECM/adhesion/cytoskeletal domains. The supplementary GO panel contained the 20 strongest broadly shared and 20 strongest divergent exact identifiers. Terms were ranked by the number of significant regions multiplied by mean absolute NES among those regions. Spearman correlations compared regional profiles across each complete shared pathway universe. These analyses were descriptive and were not interpreted as formal Genotype×Age×Region tests.

#### Extended Results

##### P50 regional analyses

###### Auditory cortex

We first examined genotype-associated transcriptional changes in the auditory cortex of P50 mice. The analysis of the pooled four-age model identified 11 differentially expressed genes in the auditory cortex, comprising five increased and six decreased genes. *Cst7* showed the strongest genotype-associated increase (log₂ fold change = 15.358, adjusted P = 1.35 × 10⁻¹⁷), reflecting its near-absence in P50 WT samples and consistently detectable expression in most age-matched 5xFAD samples. The ribosomal pseudogene *Rps3a3* was decreased (log₂ fold change = −0.867, adjusted P = 1.85 × 10⁻⁵), as was *Resp18* (log₂ fold change = −0.569, adjusted P = 0.0211), whereas *Thy1* was modestly increased (log₂ fold change = 0.729, adjusted P = 0.0400). The remaining significant transcripts were uncharacterized or predicted genes. Despite the limited number of individually significant genes, GSEA identified extensive and highly significant changes across coordinated metabolic, proteostatic, and neuronal programs.

Mitochondrial energy metabolism was the most strongly suppressed biological program. Hallmark oxidative phosphorylation showed pronounced negative enrichment in 5xFAD mice (NES = −2.765, adjusted P = 5.62 × 10⁻¹⁷), consistent with KEGG oxidative phosphorylation (NES = −2.516, adjusted P = 1.00 × 10⁻¹⁰). GO terms associated with the respiratory-chain complex (NES = −2.689, adjusted P = 1.00 × 10⁻¹⁰), mitochondrial ATP synthesis coupled to electron transport (NES = −2.646, adjusted P = 1.00 × 10⁻¹⁰), and mitochondrial inner membrane (NES = −2.534, adjusted P = 1.00 × 10⁻¹⁰) were similarly depleted. Negative enrichment extended to KEGG carbon metabolism (NES = −2.063, adjusted P = 4.29 × 10⁻⁶), the citrate cycle (NES = −2.035, adjusted P = 2.56 × 10⁻⁴), glycolysis/gluconeogenesis (NES = −1.806, adjusted P = 0.00140), and fatty-acid degradation (NES = −1.794, adjusted P = 0.00235). These concordant results indicate coordinated suppression of mitochondrial respiration and metabolic substrate utilization in the P50 5xFAD auditory cortex.

RNA processing, protein synthesis, and protein quality-control pathways showed parallel reductions. KEGG ribosome produced the most negative KEGG enrichment score (NES = −2.650, adjusted P = 1.00 × 10⁻¹⁰), consistent with the reduction in *Rps3a3* and with negative GO enrichment for the large ribosomal subunit, structural constituents of the ribosome, rRNA processing, and ribosome biogenesis. The proteasome (NES = −2.261, adjusted P = 6.59 × 10⁻⁶), spliceosome (NES = −2.083, adjusted P = 1.79 × 10⁻⁷), protein export (NES = −2.131, adjusted P = 4.38 × 10⁻⁵), and protein processing in the endoplasmic reticulum (NES = −1.716, adjusted P = 2.65 × 10⁻⁴) were also negatively enriched. GO terms related to protein folding and molecular chaperones showed the same direction. Hallmark MYC targets V1 (NES = −2.509, adjusted P = 3.31 × 10⁻¹³), MYC targets V2 (NES = −2.051, adjusted P = 3.00 × 10⁻⁴), mTORC1 signaling (NES = −1.920, adjusted P = 7.13 × 10⁻⁶), and the unfolded-protein response (NES = −1.692, adjusted P = 0.00648) further support reduced biosynthetic and proteostatic activity.

In contrast, neuronal structural and connectivity programs were positively enriched. Homophilic cell adhesion through plasma-membrane adhesion molecules showed the strongest positive GO enrichment (NES = 2.713, adjusted P = 1.00 × 10⁻¹⁰). Other prominent terms included axodendritic protein transport (NES = 2.258, adjusted P = 5.50 × 10⁻⁵), central nervous system neuron development (NES = 2.234, adjusted P = 1.36 × 10⁻⁷), protein localization to the presynapse (NES = 2.174, adjusted P = 7.53 × 10⁻⁵), plasma-membrane-bounded cell-projection morphogenesis (NES = 2.145, adjusted P = 1.00 × 10⁻¹⁰), and axonogenesis (NES = 2.031, adjusted P = 1.00 × 10⁻¹⁰). Synapse assembly (NES = 1.978, adjusted P = 8.39 × 10⁻⁹), synapse organization (NES = 1.967, adjusted P = 1.00 × 10⁻¹⁰), and regulation of synapse structure or activity (NES = 1.864, adjusted P = 4.37 × 10⁻⁹) were also positively enriched.

KEGG analysis independently supported this structural-remodeling response. ABC transporters showed the strongest positive KEGG enrichment (NES = 2.322, adjusted P = 2.85 × 10⁻⁶), followed by phosphatidylinositol signaling (NES = 2.156, adjusted P = 6.86 × 10⁻⁷), ECM–receptor interaction (NES = 1.969, adjusted P = 5.25 × 10⁻⁵), axon guidance (NES = 1.785, adjusted P = 2.44 × 10⁻⁵), ErbB signaling (NES = 1.750, adjusted P = 8.53 × 10⁻⁴), cadherin signaling (NES = 1.632, adjusted P = 4.54 × 10⁻⁵), integrin signaling (NES = 1.569, adjusted P = 0.00336), and focal adhesion (NES = 1.533, adjusted P = 0.00191). Glutamatergic and cholinergic synapse pathways were positively enriched but showed smaller effect sizes and weaker statistical support than the structural pathways. The AC profile therefore primarily indicates remodeling of neurites, synapses, membrane transport, and cell–cell or cell–matrix interactions rather than a uniform increase in synaptic transmission.

Additional enrichment patterns suggested changes in cytoskeletal and stress-response regulation. Hallmark mitotic spindle was strongly positively enriched (NES = 2.190, adjusted P = 3.85 × 10⁻⁸), accompanied by positive UV-response-down, Hedgehog-signaling, and cholesterol-homeostasis gene sets. In contrast, Hallmark DNA repair, p53 signaling, apoptosis, hypoxia, and UV-response-up were negatively enriched. Because mature cortical neurons are predominantly postmitotic, the mitotic-spindle signature is more appropriately interpreted as altered microtubule and cytoskeletal regulation, or as a contribution from non-neuronal populations, rather than evidence of neuronal proliferation.

Immune-associated findings were selective rather than uniformly activated. Although *Cst7* was strongly increased, Hallmark TNFα signaling through NF-κB was negatively enriched (NES = −1.531, adjusted P = 0.00888), as were KEGG antigen processing and presentation (NES = −1.901, adjusted P = 2.14 × 10⁻⁴) and the GO molecular function cytokine activity (NES = −1.742, adjusted P = 3.78 × 10⁻⁴). Several GO terms related to negative regulation of microglial activation or neuroinflammatory responses were positively enriched. Thus, the AC data do not support broad inflammatory activation at P50; instead, they indicate a restricted immune-associated response superimposed on more extensive metabolic and neuronal remodeling.

Collectively, the analysis identifies a marked dissociation between metabolic capacity and neuronal structural remodeling in the young 5xFAD auditory cortex. Mitochondrial respiration, intermediary metabolism, translation, RNA processing, protein folding, and proteasomal function were coordinately suppressed, whereas axonal, synaptic, adhesion, extracellular-matrix, and membrane-signaling programs were positively enriched. This combination is consistent with an early remodeling state in which neuronal and cellular connectivity programs are altered despite reduced transcriptional support for energy production and proteostasis. Because these measurements were obtained from bulk tissue, the observed pathway shifts may reflect transcriptional regulation within resident cells, changes in cellular composition, or both.

###### Inferior colliculus

The P50 genotype contrast from the pooled four-age model identified two differentially expressed genes in the inferior colliculus. *Thy1* was increased in 5xFAD mice (log₂ fold change = 0.656, adjusted P = 0.00467), whereas *Tmem184c* was modestly decreased (log₂ fold change = −0.259, adjusted P = 0.0342). Despite the limited number of individually significant genes, GSEA revealed extensive and highly significant pathway-level changes, indicating that the IC response primarily involved coordinated shifts distributed across functionally related genes.

Mitochondrial energy metabolism was the most strongly suppressed program. Hallmark oxidative phosphorylation showed exceptionally strong negative enrichment in 5xFAD mice (NES = −3.383, adjusted P = 1.72 × 10⁻³²). This result was independently supported by KEGG oxidative phosphorylation (NES = −3.162, adjusted P = 1.00 × 10⁻¹⁰) and GO terms representing the respiratory-chain complex (NES = −3.176, adjusted P = 1.00 × 10⁻¹⁰), mitochondrial ATP synthesis coupled to electron transport (NES = −3.004, adjusted P = 1.00 × 10⁻¹⁰), mitochondrial inner membrane (NES = −2.885, adjusted P = 1.00 × 10⁻¹⁰), mitochondrial ribosome (NES = −2.797, adjusted P = 1.00 × 10⁻¹⁰), and respiratory-chain complex assembly (NES = −2.659, adjusted P = 1.00 × 10⁻¹⁰). These highly concordant results demonstrate broad suppression of the transcriptional machinery supporting mitochondrial respiration and ATP production.

Multiple pathways of intermediary metabolism were similarly reduced. KEGG carbon metabolism (NES = −2.520, adjusted P = 2.37 × 10⁻¹⁰), the citrate cycle (NES = −2.387, adjusted P = 4.76 × 10⁻⁷), glycolysis/gluconeogenesis (NES = −1.992, adjusted P = 1.60 × 10⁻⁴), fatty-acid metabolism (NES = −1.797, adjusted P = 7.71 × 10⁻⁴), and fatty-acid degradation (NES = −1.776, adjusted P = 0.00323) were negatively enriched. Hallmark fatty-acid metabolism showed an even stronger reduction (NES = −2.569, adjusted P = 2.41 × 10⁻¹²), accompanied by negative enrichment of glycolysis, peroxisome, heme metabolism, and the reactive oxygen species pathway. Thus, the mitochondrial phenotype was embedded within a broader reduction in glucose, lipid, amino-acid, and redox-associated metabolism.

Biosynthetic and protein-homeostasis pathways showed parallel suppression. KEGG ribosome was strongly negatively enriched (NES = −3.052, adjusted P = 1.00 × 10⁻¹⁰), consistent with negative GO enrichment for structural constituents of the ribosome (NES = −3.073, adjusted P = 1.00 × 10⁻¹⁰), the large ribosomal subunit, cytoplasmic translation, rRNA processing, and ribosome biogenesis. The spliceosome (NES = −2.441, adjusted P = 1.00 × 10⁻¹⁰), proteasome (NES = −2.425, adjusted P = 2.73 × 10⁻⁷), and protein processing in the endoplasmic reticulum (NES = −1.588, adjusted P = 0.00145) were also negatively enriched. GO terms related to protein folding, molecular chaperones, translation initiation, and the unfolded-protein response showed the same direction.

Hallmark analysis reinforced this coordinated reduction in biosynthetic capacity. MYC targets V1 was strongly negatively enriched (NES = −3.085, adjusted P = 7.59 × 10⁻²⁶), accompanied by reductions in MYC targets V2, mTORC1 signaling, DNA repair, E2F targets, the unfolded-protein response, and protein secretion. The convergence of mitochondrial, metabolic, translational, RNA-processing, and proteostatic pathways indicates a broad reduction in transcriptional programs required to sustain energy-intensive cellular functions in the 5xFAD inferior colliculus.

In contrast, neuronal structural and receptor-associated programs were positively enriched. GO analysis identified strong positive enrichment of axon guidance (NES = 2.200, adjusted P = 1.00 × 10⁻¹⁰), synapse assembly (NES = 2.090, adjusted P = 1.00 × 10⁻¹⁰), synapse organization (NES = 2.089, adjusted P = 1.00 × 10⁻¹⁰), postsynaptic membrane (NES = 2.024, adjusted P = 1.00 × 10⁻¹⁰), and neurotransmitter-receptor activity (NES = 2.019, adjusted P = 1.24 × 10⁻⁵). Synaptic transmission, glutamatergic (NES = 1.927, adjusted P = 2.81 × 10⁻⁵), GABAergic synapse (NES = 1.884, adjusted P = 5.21 × 10⁻⁶), neuron-projection organization (NES = 1.821, adjusted P = 5.30 × 10⁻⁵), dendritic spine, and presynaptic active-zone terms were also positively enriched.

KEGG analysis independently supported this neuronal-remodeling response. Axon guidance (NES = 1.878, adjusted P = 2.43 × 10⁻⁶), glutamatergic synapse (NES = 1.721, adjusted P = 8.20 × 10⁻⁴), cholinergic synapse (NES = 1.447, adjusted P = 0.0178), and serotonergic synapse (NES = 1.373, adjusted P = 0.0415) were positively enriched. These findings indicate increased representation of synaptic structures, neurotransmitter receptors, and axonal guidance programs despite the marked suppression of metabolic and proteostatic pathways. However, selected vesicle-trafficking processes, including vesicle fusion, synaptic-vesicle acidification, and COPI-coated vesicles, were negatively enriched. The synaptic response therefore appears to represent selective remodeling rather than a uniform increase in neurotransmission.

Cell-adhesion and extracellular-matrix programs formed a second major positively enriched axis. KEGG ECM–receptor interaction showed strong positive enrichment (NES = 2.359, adjusted P = 2.36 × 10⁻⁸), as did integrin signaling (NES = 2.259, adjusted P = 7.35 × 10⁻⁹), focal adhesion (NES = 2.096, adjusted P = 2.27 × 10⁻⁸), cadherin signaling (NES = 1.928, adjusted P = 1.52 × 10⁻⁸), and adherens junctions (NES = 1.613, adjusted P = 0.00303). GO analysis similarly identified positive enrichment of homophilic cell adhesion (NES = 2.701, adjusted P = 1.00 × 10⁻¹⁰), cell-adhesion molecule binding, collagen trimers, cell-junction assembly, cell–matrix adhesion, and extracellular-matrix organization. These changes connect the neuronal response to broader remodeling of cellular contacts and the extracellular environment.

Membrane transport and intracellular signaling were also enhanced. ABC transporters (NES = 2.265, adjusted P = 3.72 × 10⁻⁶), phosphatidylinositol signaling (NES = 1.844, adjusted P = 1.76 × 10⁻⁴), calcium signaling (NES = 1.773, adjusted P = 2.09 × 10⁻⁵), and phospholipase D signaling (NES = 1.739, adjusted P = 2.89 × 10⁻⁴) were positively enriched. Cilium movement (NES = 1.592, adjusted P = 4.22 × 10⁻⁴), cilium organization, and multiple microtubule-associated processes were also increased. Conversely, the myelin-sheath gene set was strongly negatively enriched (NES = −2.770, adjusted P = 1.00 × 10⁻¹⁰), indicating that positive neuronal and extracellular remodeling occurred alongside reduced expression of myelin-associated genes.

Hallmark mitotic spindle was strongly positively enriched (NES = 2.232, adjusted P = 2.57 × 10⁻¹⁰), together with UV-response-down and Hedgehog-signaling gene sets. In contrast, E2F targets and several DNA-damage and stress-response pathways were negatively enriched. In a predominantly postmitotic neural tissue, the mitotic-spindle signature is more likely to reflect microtubule and cytoskeletal remodeling or contributions from non-neuronal populations than neuronal proliferation.

Immune-associated pathways were heterogeneous. Hallmark TNFα signaling through NF-κB was negatively enriched (NES = −1.895, adjusted P = 4.11 × 10⁻⁶), and several GO terms related to T-cell cytokine production, interferon signaling, antigen presentation, and myeloid cytokine production were reduced. In contrast, immune-system development, cytokine binding, the NLRP3 inflammasome complex, and selected antigen-receptor regulatory terms were positively enriched. These results do not support broad inflammatory activation at P50 but instead indicate selective remodeling of immune-associated transcriptional programs.

Collectively, the IC analysis identifies a pronounced dissociation between cellular metabolic capacity and structural or synaptic remodeling. Mitochondrial respiration, intermediary metabolism, translation, RNA processing, protein folding, and proteasomal function were strongly suppressed, whereas synaptic organization, neurotransmitter-receptor pathways, axon guidance, cell adhesion, extracellular-matrix interactions, calcium signaling, and membrane transport were positively enriched. This combination is consistent with an early remodeling state in which neuronal and tissue-structural programs are altered despite reduced transcriptional support for the energy-intensive processes required to sustain them. Because the data were generated from bulk tissue, the observed changes may reflect transcriptional regulation within resident cells, shifts in cellular composition, or both.

###### Cochlear nucleus

The cochlear nucleus displayed limited gene-level differential expression but extensive coordinated pathway remodeling in P50 5xFAD mice. Within the top ranked genes, two genes met the multiple-testing threshold. The ribosomal protein gene *Rps3a1* was increased in 5xFAD mice (log₂ fold change = 0.982, adjusted P = 0.0198), whereas *Aox4* was decreased (log₂ fold change = −2.826, adjusted P = 0.0198). Thus, individual transcript-level changes were modest after multiple-testing correction, but gene-set enrichment analysis revealed broad and statistically robust shifts distributed across functionally related genes.

Mitochondrial metabolism represented the dominant positively enriched program. The Hallmark oxidative phosphorylation gene set showed strong positive enrichment in 5xFAD mice (NES = 2.355, adjusted P = 1.27 × 10⁻¹¹). This result was independently supported by KEGG oxidative phosphorylation (NES = 2.175, adjusted P = 6.72 × 10⁻⁸) and multiple GO terms related to the mitochondrial inner membrane, respiratory-chain assembly, mitochondrial translation, and ATP synthesis. Central carbon metabolism was also positively enriched, including KEGG carbon metabolism (NES = 2.346, adjusted P = 1.51 × 10⁻⁹), the citrate cycle (NES = 2.167, adjusted P = 1.94 × 10⁻⁵), and glycolysis/gluconeogenesis (NES = 2.011, adjusted P = 4.35 × 10⁻⁵). Hallmark fatty-acid metabolism (NES = 1.951, adjusted P = 2.46 × 10⁻⁵), glycolysis (NES = 1.525, adjusted P = 0.00824), and peroxisome genes (NES = 1.618, adjusted P = 0.0117) further indicated coordinated enhancement of energy-producing and substrate-utilization pathways.

Biosynthetic and protein-homeostasis pathways were similarly enriched. KEGG ribosome produced the strongest positive NES among the KEGG pathways (NES = 2.513, adjusted P = 1.00 × 10⁻¹⁰), consistent with the increase in *Rps3a1* and with positive GO enrichment for structural constituents of the ribosome (NES = 2.447, adjusted P = 1.00 × 10⁻¹⁰), ribosomal subunits, rRNA processing, and ribosome biogenesis. The spliceosome (NES = 2.065, adjusted P = 9.57 × 10⁻⁸), proteasome (NES = 1.875, adjusted P = 4.81 × 10⁻⁴), and protein processing in the endoplasmic reticulum (NES = 1.656, adjusted P = 4.50 × 10⁻⁴) were also positively enriched. Hallmark MYC targets V1 showed the highest Hallmark NES (NES = 2.383, adjusted P = 5.78 × 10⁻¹²), accompanied by MYC targets V2, DNA repair, and mTORC1 signaling. Together, these findings indicate coordinated enrichment of mitochondrial, translational, RNA-processing, and proteostatic programs in the 5xFAD cochlear nucleus.

In contrast, neuronal communication and structural-connectivity pathways were predominantly negatively enriched. GO analysis identified reductions in excitatory chemical synaptic transmission (NES = −1.886, adjusted P = 0.00474), protein localization to the presynapse (NES = −1.864, adjusted P = 0.00421), neurotransmitter-receptor activity (NES = −1.677, adjusted P = 0.00277), and synapse assembly (NES = −1.607, adjusted P = 2.72 × 10⁻⁵). KEGG axon guidance (NES = −1.404, adjusted P = 0.0122) and serotonergic synapse signaling (NES = −1.323, adjusted P = 0.0364) were also negatively enriched. These changes suggest that increased metabolic and biosynthetic pathway enrichment does not reflect a uniform elevation of neuronal function but instead coincides with reduced expression of gene programs supporting synaptic organization, neurotransmitter signaling, and axonal protein trafficking.

Cell-adhesion and junctional programs showed a parallel negative shift. Homophilic cell adhesion through plasma-membrane adhesion molecules was among the most strongly depleted GO processes (NES = −2.069, adjusted P = 6.27 × 10⁻⁸). KEGG cadherin signaling (NES = −1.602, adjusted P = 1.28 × 10⁻⁵), adherens junctions (NES = −1.601, adjusted P = 0.00557), integrin signaling (NES = −1.470, adjusted P = 0.00840), and focal adhesion (NES = −1.459, adjusted P = 0.00609) were likewise negatively enriched. These findings connect the synaptic changes to broader remodeling of cellular interactions and structural organization within the cochlear nucleus.

Immune-associated enrichment was predominantly negative, although individual GO terms indicated that this response was not uniform. Hallmark inflammatory response (NES = −1.447, adjusted P = 0.0375) and allograft rejection (NES = −1.434, adjusted P = 0.0383) were negatively enriched, as were KEGG B-cell receptor signaling (NES = −1.680, adjusted P = 0.00374) and GO pathways related to antigen presentation and leukocyte chemotaxis. Accordingly, the CN profile does not support broad inflammatory activation at P50; instead, it indicates selective suppression or redistribution of immune-associated transcriptional programs.

Collectively, the cochlear nucleus response in young 5xFAD mice was characterized by a pronounced dissociation between positively enriched metabolic and biosynthetic pathways and negatively enriched synaptic, adhesion, and immune-associated programs. The convergence of Hallmark, KEGG, and GO results supports a coordinated increase in mitochondrial energy metabolism, RNA processing, translation, and protein quality control, potentially reflecting an early adaptive or compensatory state. However, the simultaneous depletion of synaptic and structural-connectivity pathways suggests that this molecular response may occur alongside early disruption of neuronal communication. Because these data were generated from bulk tissue, contributions from altered cellular composition cannot be distinguished from transcriptional regulation within individual cell populations.

###### Cochlea

The analysis of P50 cochlea samples revealed limited gene-level differential expression but extensive pathway-level remodeling in the cochlea. Within the ranked top genes, two genes met the multiple-testing threshold, and both were increased in 5xFAD mice. *Ighv2-3* showed a particularly large estimated effect (log₂ fold change = 12.450, adjusted P = 2.21 × 10⁻⁸), while the uncharacterized transcript *Gm37261* was also increased (log₂ fold change = 4.178, adjusted P = 0.0328). In contrast to this sparse gene-level result, GSEA identified coordinated changes across numerous functional programs, indicating that the cochlear response primarily comprised distributed shifts among biologically related genes.

Suppression of energy metabolism was one of the strongest features of the cochlear transcriptome. Hallmark oxidative phosphorylation was markedly negatively enriched in 5xFAD mice (NES = −2.666, adjusted P = 6.73 × 10⁻¹⁶). This result was supported by negative enrichment of KEGG oxidative phosphorylation (NES = −1.821, adjusted P = 2.49 × 10⁻⁵), carbon metabolism (NES = −2.254, adjusted P = 2.05 × 10⁻⁸), the citrate cycle (NES = −2.094, adjusted P = 1.06 × 10⁻⁴), and glycolysis/gluconeogenesis (NES = −1.810, adjusted P = 0.00105). GO terms representing the mitochondrial matrix, mitochondrial inner membrane, mitochondrial gene expression, cellular respiration, and respiratory-chain assembly were similarly negatively enriched. Hallmark fatty-acid metabolism (NES = −1.664, adjusted P = 0.00103) and the reactive oxygen species pathway (NES = −1.868, adjusted P = 0.00205) were also reduced. Together, these results indicate broad transcriptional suppression of mitochondrial energy production and metabolic substrate utilization in the 5xFAD cochlea.

Biosynthetic and protein-homeostasis programs showed parallel reductions. Hallmark MYC targets V1 produced the most negative Hallmark enrichment score (NES = −2.856, adjusted P = 3.23 × 10⁻²¹), accompanied by negative enrichment of MYC targets V2, E2F targets, mTORC1 signaling, and DNA repair. KEGG aminoacyl-tRNA biosynthesis (NES = −2.346, adjusted P = 2.31 × 10⁻⁷), proteasome (NES = −2.311, adjusted P = 5.07 × 10⁻⁷), and ribosome (NES = −2.019, adjusted P = 4.93 × 10⁻⁸) were strongly negatively enriched. GO analysis independently identified reductions in tRNA aminoacylation, the proteasome complex, spliceosomal assembly, ribosome biogenesis, rRNA processing, and ribosomal subunits. These concordant findings suggest that impaired energy metabolism is accompanied by reduced transcriptional investment in RNA processing, protein synthesis, and protein quality control.

Neuronal and synaptic programs were also broadly suppressed. KEGG synaptic vesicle cycle (NES = −2.206, adjusted P = 6.04 × 10⁻⁷), glutamatergic synapse (NES = −2.092, adjusted P = 4.64 × 10⁻⁷), long-term potentiation (NES = −2.036, adjusted P = 4.20 × 10⁻⁵), and GABAergic synapse (NES = −1.681, adjusted P = 0.00142) were negatively enriched. Consistent GO results included the presynaptic active zone (NES = −2.471, adjusted P = 1.00 × 10⁻¹⁰), ionotropic glutamate receptor complex (NES = −2.418, adjusted P = 3.71 × 10⁻⁷), GABAergic synapse (NES = −2.322, adjusted P = 1.00 × 10⁻¹⁰), protein localization to the synapse (NES = −2.273, adjusted P = 5.76 × 10⁻⁹), and regulation of synaptic plasticity (NES = −2.163, adjusted P = 1.00 × 10⁻¹⁰). Thus, the metabolic and proteostatic changes coincided with reduced expression of gene programs supporting neurotransmission, synaptic-vesicle handling, and activity-dependent plasticity.

The synaptic response was nevertheless selective rather than uniform. Neuroactive ligand–receptor interaction was positively enriched in KEGG analysis (NES = 1.438, adjusted P = 0.00235), and the GO cholinergic synapse term was positively enriched (NES = 2.093, adjusted P = 7.05 × 10⁻⁵). These findings suggest remodeling of receptor and signaling composition against a broader background of reduced glutamatergic, GABAergic, and synaptic-vesicle programs.

In contrast to the metabolic and synaptic suppression, extracellular and sensory-structure programs were positively enriched. Hallmark epithelial–mesenchymal transition showed strong positive enrichment (NES = 2.029, adjusted P = 1.57 × 10⁻⁷), accompanied by Hallmark angiogenesis (NES = 1.846, adjusted P = 0.00276). GO analysis identified positive enrichment of collagen trimers (NES = 2.145, adjusted P = 4.83 × 10⁻⁷), extracellular matrix structural constituents (NES = 2.116, adjusted P = 1.27 × 10⁻⁷), mesenchymal-cell proliferation, blood-vessel branching, and endothelial-cell migration. These results indicate substantial remodeling of extracellular-matrix and vascular-associated transcriptional programs.

Cochlea-associated structural programs were also prominently enriched. Cilium movement (NES = 1.971, adjusted P = 2.72 × 10⁻⁸), cilium-dependent cell motility (NES = 1.947, adjusted P = 3.76 × 10⁻⁷), hair-cell differentiation (NES = 1.975, adjusted P = 6.54 × 10⁻⁵), stereocilium (NES = 1.920, adjusted P = 1.83 × 10⁻⁴), and stereocilium bundle (NES = 1.910, adjusted P = 5.99 × 10⁻⁵) were positively enriched in GO terms.

Positive KEGG enrichment of Hedgehog signaling (NES = 1.635, adjusted P = 0.00945), Wnt signaling (NES = 1.397, adjusted P = 0.0268), Ras signaling (NES = 1.389, adjusted P = 0.0107), and PI3K–Akt signaling (NES = 1.302, adjusted P = 0.0156) further supports altered developmental, structural, and tissue-maintenance signaling. These signatures are consistent with cochlear remodeling but do not, by themselves, establish increased hair-cell abundance or regenerative activity.

Immune-associated findings were heterogeneous. The marked increase in *Ighv2-3* and positive enrichment of KEGG NF-κB signaling (NES = 1.561, adjusted P = 0.00506) suggest a selective immune-related component. However, several GO terms related to acute inflammatory responses, myeloid immunity, mast-cell cytokine production, and leukocyte cytotoxicity were negatively enriched. The data therefore do not support uniform activation of cochlear inflammation at P50. Instead, they suggest selective changes in immune-cell-associated transcripts or immune signaling superimposed on broader metabolic and structural remodeling.

Collectively, the P50 5xFAD cochlea exhibited a pronounced separation between negatively enriched metabolic, proteostatic, and synaptic programs and positively enriched extracellular-matrix, vascular, ciliary, and hair-cell-associated programs. This pattern is consistent with early cochlear stress and tissue remodeling accompanied by reduced molecular support for energy-intensive biosynthesis and neurotransmission. Because the measurements were obtained from bulk cochlear tissue, the observed enrichment patterns may reflect transcriptional regulation within resident cells, changes in cellular composition, or both.

##### Age-dependent regional analyses

###### Auditory cortex

Age-specific differential expression revealed a marked expansion of the 5xFAD transcriptional phenotype after P50. DESeq2 identified 11 differentially expressed genes at P50, compared with 527 at 8 months, 647 at 11 months, and 277 at 16 months (FDR < 0.05). Consistent with this progression, the number of significantly enriched Hallmark pathways increased from 20 at P50 to 21, 31, and 29 at 8, 11, and 16 months, respectively. KEGG and simplified GO analyses showed the greatest breadth at 11 months, identifying 158 KEGG pathways and 647 GO terms at q < 0.05.

At P50, 5xFAD mice primarily exhibited suppression of biosynthetic and metabolic programs. Oxidative phosphorylation (NES = −2.76), MYC Targets V1 (NES = −2.51), fatty-acid metabolism, adipogenesis, mTORC1 signaling, DNA repair, and p53 signaling were negatively enriched. KEGG analysis independently identified negative enrichment of ribosome (NES = −2.65), proteasome (NES = −2.26), antigen processing and presentation (NES = −1.90), and phagocytosis (NES = −1.55). In contrast, mitotic spindle genes were positively enriched (NES = 2.19). Thus, the earliest detectable cortical phenotype involved coordinated metabolic and proteostatic disruption, with relatively limited gene-level differential expression.

By 8 months, the genotype-associated expression pattern had shifted toward a strong immune and inflammatory phenotype. Interferon-γ response (NES = 2.67), allograft rejection (NES = 2.82), inflammatory response (NES = 2.63), interferon-α response, IL6–JAK–STAT3 signaling, TNF signaling through NF-κB, complement, and coagulation were positively enriched. KEGG analysis similarly identified positive enrichment of cytokine–receptor interactions, antigen processing and presentation, phagocytosis, lysosome biogenesis, hematopoietic cell lineage, and leukocyte transendothelial migration. Genes contributing to these changes included *Fcrl2*, *Csf3r*, *Siglecf*, *Itgax*, *Ccl6*, *Ncf2*, *Mpeg1*, *Cd68*, and *Trem2*. These results support increased myeloid and antigen-processing activity, although bulk RNA sequencing cannot distinguish altered cellular composition from cell-intrinsic activation.

The immune phenotype was strongest and most broadly accompanied by metabolic and structural remodeling at 11 months. Interferon-γ and interferon-α responses reached NES values of 3.18, while coagulation, complement, IL6–JAK–STAT3 signaling, and the extracellular-matrix/remodeling signature represented by the Hallmark epithelial–mesenchymal transition gene set were also strongly enriched. The latter signature increased from NES = 1.79 at 8 months to 2.69 at 11 months and remained elevated at 16 months (NES = 2.49). Its leading-edge and differentially expressed genes included *Spp1*, *Tgfb1*, *Itgb5*, *Col1a1*, *Col3a1*, *Bgn*, *Emp3*, and *Serpine1*. In the brain, this signature is most appropriately interpreted as extracellular-matrix and tissue remodeling rather than epithelial transdifferentiation. Positive enrichment of ribosome, proteasome, MYC Targets V1, mTORC1 signaling, glycolysis, DNA repair, and ferroptosis-related genes further indicated broad reorganization of protein synthesis, bioenergetics, and cellular stress responses.

Neuronal signaling pathways changed in the opposite direction. Axon guidance and phosphatidylinositol and ErbB signaling were positively enriched at P50 but became negatively enriched or nonsignificant at later ages. At 11 months, glutamatergic synapse, cholinergic synapse, neuroactive ligand signaling, and axon guidance were all negatively enriched in 5xFAD mice. The formal interaction analyses confirmed negative shifts in axon guidance, phosphatidylinositol signaling, and ErbB signaling at 8, 11, and 16 months relative to P50.

Glutamatergic and cholinergic synapse interactions became significant at 11 and 16 months. GO analysis provided concordant evidence from negative shifts in synapse assembly, GABAergic synapse, axonogenesis, cation-channel complexes, and voltage-gated potassium-channel complexes. These changes link the evolving inflammatory phenotype to disruption of neuronal communication and synaptic organization in the auditory cortex.

At 16 months, inflammatory, phagolysosomal, antigen-presentation, and extracellular-matrix programs remained positively enriched, but the metabolic phenotype was not uniformly progressive. Oxidative phosphorylation and MYC Targets V1 were again negatively enriched, whereas ribosome and several stress-response pathways no longer reached age-specific significance. Nevertheless, their positive interaction NES values indicated that these effects remained shifted toward more positive values relative to the stronger P50 deficits. Therefore, positive interaction enrichment should not invariably be interpreted as pathway activation at the later age. Rather, it may represent attenuation of an early negative genotype effect.

The omnibus likelihood-ratio test identified 193 genes with significant Genotype×Age effects. Directional interaction tests identified 246 genes at 8 months and 274 genes at 11 months relative to P50, with positive coefficients predominating. At 16 months, only *Rps3a3* and *Cst7* passed the gene-level interaction FDR threshold, despite extensive pathway-level interactions. This difference indicates that the late-stage pathway signals arose from coordinated shifts across many genes rather than from large effects in numerous individual genes. Collectively, the results define a temporal transition from early metabolic and proteostatic abnormalities to sustained immune activation, phagolysosomal engagement, and extracellular-matrix remodeling, accompanied by reduced neuronal and synaptic pathway activity. The maximal breadth of these changes at 11 months and their partial persistence at 16 months support a dynamic, nonmonotonic evolution of auditory cortical pathology during 5xFAD disease progression.

###### Inferior colliculus

To characterize age-dependent changes in the 5xFAD transcriptional phenotype in the inferior colliculus (IC), we modeled the Genotype×Age interaction using P50 as the reference. The omnibus likelihood-ratio test identified 66 genes with significant Genotype×Age effects at FDR <0.05. Directional Wald contrasts identified 60 significant genes at 8 months relative to P50, including 53 positive and 7 negative interaction coefficients; 65 genes at 11 months, including 41 positive and 24 negative coefficients; and 9 genes at 16 months, including 5 positive and 4 negative coefficients. At the pathway level, the interaction analysis identified 27, 33, and 31 significant Hallmark pathways; 143, 199, and 173 KEGG pathways; and 504, 591, and 528 simplified GO terms at 8, 11, and 16 months, respectively. Thus, the 11-month comparison showed the greatest breadth of age-dependent pathway remodeling.

Immune and inflammatory programs emerged by 8 months and remained prominent through 16 months. Interferon-α response changed from a nonsignificant NES of 0.50 at P50 to significant positive enrichment at 8, 11, and 16 months (NES = 2.03, 2.64, and 3.05, respectively). Interferon-γ response showed a similar trajectory (NES = 0.62, 2.07, 2.85, and 2.90), as did IL6–JAK–STAT3 signaling (NES = 0.80, 2.52, 2.63, and 2.72). Inflammatory response and complement also became positively enriched at all three later ages, with significant positive interaction NES in each contrast. Concordant KEGG and GO results included antigen processing and presentation, phagocytosis, complement and coagulation cascades, hematopoietic cell lineage, inflammatory response, leukocyte activation, cell killing, Toll-like receptor signaling, and respiratory burst. Disease-named KEGG pathways, including allograft rejection and several infection-associated sets, reflected shared immune-response genes and do not indicate the corresponding diseases or infections.

Gene-level results supported the pathway-level immune transition. The omnibus LRT identified age-dependent expression profiles for *Lag3*, *Ccl6*, *Tyrobp*, *Mpeg1*, *Clec7a*, *Trem2*, *Itgax*, *Adgre1*, *Ctss*, *Cd74*, and the complement genes *C1qa*, *C1qb*, and *C1qc*. Positive interaction coefficients at 8 and/or 11 months were also detected for many of these genes. Leading-edge genes in the interferon, complement, and inflammatory pathways included *Cd74*, *H2-Aa*, *H2-Eb1*, *Irf8*, *Lcp2*, *Psmb8*, *Cd86*, *Fcgr1*, *Myd88*, and *Stat1*. Together, these results indicate that the age-dependent IC response included coordinated activation of antigen-presentation, innate immune, complement, and phagocytic transcriptional programs.

Bioenergetic and biosynthetic pathways exhibited a distinct temporal pattern. Hallmark oxidative phosphorylation was strongly negatively enriched at P50 (NES = −3.38), became neutral at 8 months (NES = −0.65), shifted modestly positive at 11 months (NES = 1.12), and was again negative but nonsignificant at 16 months (NES = −1.13). Nevertheless, all three interaction contrasts were strongly positive (NES = 3.06, 3.03, and 2.79), because each later genotype effect was more positive than the P50 effect. KEGG oxidative phosphorylation showed the same pattern. Hallmark MYC Targets V1, fatty-acid metabolism, adipogenesis, mTORC1 signaling, and reactive oxygen species response, together with KEGG ribosome, proteasome, carbon metabolism, and the tricarboxylic-acid cycle, also shifted positively relative to P50. Ribosome changed from strong negative enrichment at P50 (NES = −3.05) to positive enrichment at 11 and 16 months (NES = 1.59 and 1.44). Proteasome similarly changed from NES = −2.42 at P50 to significant positive enrichment at 11 and 16 months (NES = 1.74 and 1.57). These findings identify 11 months as the clearest transition toward increased enrichment of mitochondrial, translational, and proteostatic gene sets. The positive interactions at 16 months often reflected persistent attenuation of the strongly negative P50 phenotype rather than sustained positive enrichment.

Neuronal signaling and synaptic-organization pathways showed the reciprocal pattern. The GABAergic synapse GO term changed from significant positive enrichment at P50 (NES = 1.88) to negative enrichment at 8 months and significant negative enrichment at 11 and 16 months (NES = −1.24, −1.87, and −1.97). Synapse assembly changed from NES = 2.09 at P50 to −1.37, −2.01, and −1.49 at the three later ages. Regulation of synapse organization, neuron spine, dendritic spine, presynaptic active zone, and synaptic membrane adhesion showed concordant negative interaction effects. KEGG glutamatergic synapse, calcium signaling, axon guidance, neuroactive ligand–receptor interaction, and long-term potentiation also shifted negatively relative to P50. Representative negative gene-level interactions at 11 months included *Pcdhb16* and *Oprm1*, although most synaptic pathway signals arose from coordinated moderate changes across many genes rather than from large changes in individual transcripts.

Ciliary and axonemal programs also declined relative to P50. Cilium organization changed from significant positive enrichment at P50 (NES = 1.54) to significant negative enrichment at 8 and 11 months (NES = −2.04 and −1.58) and a weaker negative value at 16 months (NES = −1.07). The corresponding interaction NES values were negative at all three ages. Sperm flagellum and sperm axoneme assembly terms showed similar patterns; in neural tissue, these annotations represent shared motile-cilia and axonemal machinery rather than reproductive processes. *Cfap53* showed a significant omnibus Genotype×Age effect, and *Dnah6* had a significant negative interaction coefficient at 11 months, providing gene-level support for altered ciliary or axonemal organization.

Extracellular-matrix-related pathways showed remodeling rather than uniform activation. Coagulation changed from a negative value at P50 to significant positive enrichment at all later ages, while collagen catabolism and glycosaminoglycan degradation increased most clearly at 11 months. In contrast, KEGG ECM–receptor interaction remained positively enriched at several ages but showed significant negative interaction effects because its enrichment was weaker than at P50. Age-dependent expression of *Postn* and the positive 11-month interaction of *Serpinf2* further supported changes in matrix-associated and coagulation-related transcription. These opposing pathway trajectories indicate reorganization of extracellular-matrix processes rather than a simple global increase in matrix production.

Although only nine individual genes passed the Wald FDR threshold at 16 months, hundreds of pathways retained significant interaction signals. This difference is consistent with coordinated but individually modest changes distributed across many genes, which GSEA can detect even when few transcripts pass genome-wide correction. Overall, the IC transcriptional phenotype shifted from P50 enrichment patterns dominated by synaptic, neuronal-signaling, cytoskeletal, and ciliary programs toward immune, phagocytic, complement, and inflammatory programs at later ages. Mitochondrial, translational, and proteostatic pathways underwent their strongest transition at 11 months, whereas immune activation and negative synaptic interactions persisted through 16 months.

###### Cochlear nucleus

Genotype×Age analysis identified a strongly non-monotonic transcriptional trajectory in the cochlear nucleus (CN). Relative to P50, the number of significant interaction pathways increased from 9 Hallmark, 18 KEGG, and 96 GO terms at 8 months to 27 Hallmark, 165 KEGG, and 512 GO terms at 16 months. The 11-month contrast showed an intermediate number of significant pathways but frequently differed in direction from both 8 and 16 months. These results indicate that the 5xFAD-associated transcriptional state does not progress monotonically across age.

At 8 months, significant interactions predominantly favored immune and inflammatory programs. All nine significant Hallmark interactions had positive NES values and included allograft rejection (NES = 1.98), interferon-γ response (NES = 1.78), interferon-α response (NES = 1.73), coagulation (NES = 1.67), complement (NES = 1.59), IL6–JAK–STAT3 signaling (NES = 1.56), and inflammatory response (NES = 1.54). KEGG analysis independently identified positive interactions for complement and coagulation cascades, NF-κB signaling, cytokine–receptor interactions, hematopoietic cell lineage, and pathways containing antibacterial and antiviral response genes. GO terms further implicated immunoglobulin binding, complement binding, antigen processing and presentation, neutrophil chemotaxis, B-cell-mediated immunity, and leukocyte proliferation. Thus, the early CN response involved coordinated engagement of innate and adaptive immune-associated transcriptional programs rather than isolated changes in individual inflammatory genes. Consistent with this interpretation, several genes that were differentially expressed between 5xFAD and WT at 8 months also contributed to the leading edges of significant interaction pathways, including *Ccl4*, *Lgals3*, *C1qa*, and *Timp1*.

The 11-month profile differed markedly from the 8-month response. All five significant Hallmark interactions were negative and comprised interferon-α response (NES = −1.94), epithelial–mesenchymal transition (NES = −1.88), oxidative phosphorylation (NES = −1.67), interferon-γ response (NES = −1.62), and myogenesis (NES = −1.54). The epithelial–mesenchymal transition signature in neural tissue is best interpreted as an extracellular-matrix and tissue-remodeling program. Concordantly, KEGG identified negative interactions for ECM–receptor interaction, integrin signaling, ribosome, oxidative phosphorylation, glutathione metabolism, and several amino-acid and xenobiotic metabolic pathways. In contrast, neuroactive ligand signaling showed a positive interaction (NES = 1.78), while GO analysis identified positive interactions for neuron projection organization, synapse assembly, postsynaptic membrane-potential regulation, AMPA receptor clustering, synaptic vesicle transport, and GABAergic synapse. These findings define 11 months as a distinct intermediate state characterized by reduced interferon, matrix, translational, and mitochondrial interaction signals together with relative enrichment of synaptic-organization programs. The pathway changes do not establish improved neuronal function but indicate a redistribution of the genotype-associated transcriptional effect toward genes involved in synaptic structure and signaling.

By 16 months, the CN displayed its broadest transcriptional divergence from P50. Positive Hallmark interactions included inflammatory response (NES = 2.25), allograft rejection (NES = 2.16), interferon-γ response (NES = 2.11), interferon-α response (NES = 2.11), IL6–JAK–STAT3 signaling (NES = 2.07), coagulation, complement, TNF–NF-κB signaling, epithelial–mesenchymal transition, apical junction, and TGF-β signaling. KEGG and GO analyses reinforced this pattern through positive interactions involving B-cell receptor signaling, cytokine signaling, antigen processing and presentation, Fcγ-receptor-mediated phagosome formation, extracellular-matrix organization, integrin signaling, focal adhesion, and cell-adhesion molecules.

Recurrent leading-edge genes included *Csf1*, *Ripk2*, *Cxcl10*, *Ccl2*, *Tap1*, *Tnf*, and *Cd74* in immune pathways and *Serpine1*, *Fbn1*, *Thbs1*, *Fn1*, *Tgfb1*, and *Timp1* in matrix-remodeling pathways. These coordinated changes support a late inflammatory and tissue-remodeling state but do not, by themselves, distinguish altered cell abundance from transcriptional activation within resident cells.

The late immune and matrix response coincided with strong negative interactions in bioenergetic and biosynthetic pathways. Oxidative phosphorylation showed the strongest negative Hallmark interaction at 16 months (NES = −3.17), accompanied by MYC targets V1 (NES = −2.91), mTORC1 signaling (NES = −2.18), MYC targets V2 (NES = −2.10), DNA repair (NES = −1.99), fatty-acid metabolism (NES = −1.93), unfolded-protein response (NES = −1.87), and glycolysis (NES = −1.49). KEGG analysis similarly identified negative interactions for oxidative phosphorylation, ribosome, the tricarboxylic-acid cycle, carbon metabolism, amino-acid biosynthesis, spliceosome, proteasome, and protein processing in the endoplasmic reticulum. GO terms localized these changes to the respiratory-chain complex, mitochondrial translation and gene expression, ATP synthesis, ribosome biogenesis, protein folding, and chaperone complexes. Age-specific GSEA confirmed negative enrichment of oxidative phosphorylation and ribosome pathways in 5xFAD relative to WT at 16 months, contrasting with their positive enrichment at P50. Collectively, these results indicate that late immune and extracellular-matrix remodeling in the CN accompanies a pronounced shift away from mitochondrial energy production, protein synthesis, and proteostasis.

Synaptic changes at 16 months were heterogeneous rather than uniformly suppressive. Axon guidance, synapse organization, and synapse assembly showed positive interactions, whereas the synaptic vesicle cycle and several vesicle-maturation or acidification terms showed negative interactions. The late CN phenotype therefore combines immune activation and matrix remodeling with disruption of specific neuronal and metabolic modules, rather than reflecting a generalized loss of all neuronal transcriptional programs.

At the individual-gene level, the omnibus likelihood-ratio test identified only three genes with significant Genotype×Age effects: *Gm56815*, *Fcho1*, and *Scoc*. None of the three directional age-versus-P50 interaction contrasts contained an individual gene significant at FDR < 0.05. The detection of extensive pathway-level interactions despite few individually significant genes indicates that the CN response is distributed across coordinated groups of moderately changing genes. This pattern is consistent with the greater sensitivity of rank-based enrichment analysis for detecting concerted biological shifts that do not depend on large effects at single loci.

###### Cochlea

The cochlea exhibited extensive and strongly age-dependent pathway remodeling. Using P50 as the reference, the interaction GSEA identified 24, 19, and 28 significant Hallmark pathways at 8, 11, and 16 months, respectively. The corresponding analyses identified 75, 85, and 103 significant KEGG pathways and 328, 467, and 522 significant GO terms. Hallmark significance was defined by BH-adjusted P < 0.05, whereas KEGG and GO significance was defined by q-value < 0.05. A positive interaction NES indicates that the 5xFAD-versus-WT effect shifted positively relative to P50; it does not necessarily indicate positive enrichment in 5xFAD mice at the later age.

At 8 months, the predominant changes involved extracellular-matrix organization, cytoskeletal structure, cell adhesion, and inflammatory signaling. The Hallmark epithelial–mesenchymal transition signature increased from an age-specific NES of 2.03 at P50 to 3.87 at 8 months, with a strong positive interaction relative to P50 (interaction NES = 3.26, adjusted P = 5.3 × 10⁻³¹). Myogenesis showed a similarly positive interaction (NES = 2.72, adjusted P = 3.9 × 10⁻¹⁹). KEGG cytoskeleton in muscle cells increased from NES = 1.12 at P50 to NES = 3.24 at 8 months (interaction NES = 3.19, q-value = 9.2 × 10⁻⁹). Integrin signaling increased from NES = 1.19 to 3.01 (interaction NES = 2.66, q-value = 9.2 × 10⁻⁹), and ECM–receptor interaction increased from NES = 1.24 to 2.86 (interaction NES = 2.56, q-value = 2.4 × 10⁻⁸). *Comp*lement and coagulation cascades also shifted positively (interaction NES = 2.29, q-value = 1.8 × 10⁻⁵). These coordinated changes indicate enhanced matrix remodeling, adhesion, and tissue-response programs at 8 months.

The transcriptional state changed markedly at 11 months. Several inflammatory Hallmark signatures that were positively enriched at 8 months became negatively enriched at 11 months. Inflammatory response changed from NES = 1.59 at 8 months to NES = −2.13 at 11 months (interaction NES = −1.76, adjusted P = 1.0 × 10⁻⁴). IL6–JAK–STAT3 signaling changed from NES = 1.87 to −2.53 (interaction NES = −2.03, adjusted P = 5.6 × 10⁻⁵), and interferon-γ response changed from NES = 1.88 to −2.11 (interaction NES = −1.77, adjusted P = 1.0 × 10⁻⁴). Extracellular-matrix and adhesion pathways displayed similar negative shifts, including the Hallmark epithelial–mesenchymal transition signature (interaction NES = −1.92, adjusted P = 3.7 × 10⁻⁶).

In contrast, neuronal and synaptic pathways shifted strongly toward positive enrichment at 11 months. KEGG glutamatergic synapse changed from NES = −2.09 at P50 and −1.84 at 8 months to NES = 2.52 at 11 months (interaction NES = 2.58, q-value = 1.1 × 10⁻⁸). GABAergic synapse changed from NES = −1.68 at P50 and −1.70 at 8 months to NES = 2.63 at 11 months (interaction NES = 2.44, q-value = 1.1 × 10⁻⁷). Synaptic vesicle cycle, dopaminergic synapse, neuroactive ligand signaling, long-term potentiation, and long-term depression showed concordant positive interactions. GO analysis supported this pattern: presynaptic active zone reached an age-specific NES of 2.97 at 11 months (interaction NES = 3.12, q-value = 8.5 × 10⁻⁹), while neurotransmitter receptor complex, glutamatergic transmission, asymmetric synapse, and signal release from synapse also shifted positively. These results identify a pronounced but transient enhancement of synaptic gene-set enrichment at 11 months.

Cilium- and axoneme-associated programs moved in the opposite direction at 11 months. GO cilium movement was strongly negatively enriched at this age (age-specific NES = −2.99; interaction NES = −3.09, q-value = 8.5 × 10⁻⁹). Microtubule bundle formation, flagellated sperm motility, and other axonemal terms showed concordant negative interactions. In the cochlea, these gene sets likely reflect motile-cilia and microtubule-based structural programs; their coordinated suppression indicates disruption of ciliary and axonemal organization rather than a generalized reduction in all cellular structures.

By 16 months, the cochlear profile shifted toward strong activation of biosynthetic, cell-cycle, mitochondrial, and proteostasis-related pathways. Hallmark E2F targets changed from NES = −2.46 at P50 to NES = 3.39 at 16 months (interaction NES = 3.41, adjusted P = 8.7 × 10⁻⁴²). Heme metabolism changed from NES = −2.13 to 3.16 (interaction NES = 3.21, adjusted P = 3.3 × 10⁻³³), and G2M checkpoint changed from NES = −1.84 to 2.98 (interaction NES = 3.08, adjusted P = 5.3 × 10⁻³⁰). Oxidative phosphorylation shifted progressively from NES = −2.67 at P50 to −1.30 at 8 months, −0.41 at 11 months, and 1.41 at 16 months. Its interaction was significant at every later age and strongest at 16 months (interaction NES = 2.48, adjusted P = 1.6 × 10⁻¹⁴).

KEGG analysis confirmed the late activation of biosynthetic and maintenance programs. Ribosome changed from NES = −2.02 at P50 to NES = 2.85 at 16 months (interaction NES = 3.17, q-value = 3.6 × 10⁻⁹). DNA replication changed from NES = −2.09 to 2.90 (interaction NES = 3.03, q-value = 3.6 × 10⁻⁹), cell cycle changed from NES = −1.31 to 2.61 (interaction NES = 2.69, q-value = 3.6 × 10⁻⁹), and proteasome changed from NES = −2.31 to 2.27 (interaction NES = 2.61, q-value = 3.8 × 10⁻⁸). DNA-repair, spliceosomal, nucleocytoplasmic-transport, and porphyrin-metabolism pathways showed concordant positive shifts. Carbon metabolism and proteasome were the only KEGG pathways with significant interactions in the same positive direction at all three later ages, indicating persistent strengthening of metabolic and protein-quality-control programs relative to P50.

The synaptic enrichment observed at 11 months was not maintained at 16 months. Glutamatergic synapse changed from NES = 2.52 at 11 months to NES = −3.04 at 16 months (interaction NES = −2.28, q-value = 6.3 × 10⁻⁷). Neuroactive ligand signaling similarly changed from NES = 2.54 to −2.64 (interaction NES = −2.22, q-value = 3.2 × 10⁻⁸). GABAergic, dopaminergic, and cholinergic synapse pathways also became negatively enriched. GO terms related to presynaptic organization, neurotransmitter receptors, dendritic structure, and synaptic transmission showed parallel negative enrichment. Thus, the 11-month increase in synaptic gene-set enrichment represented a temporally restricted state rather than a sustained progression.

Ciliary and axonemal suppression persisted at 16 months. Axoneme assembly showed the strongest negative GO interaction (NES = −3.20, q-value = 6.6 × 10⁻⁹), followed by axoneme (NES = −3.12, q-value = 6.6 × 10⁻⁹), microtubule bundle formation (NES = −3.05, q-value = 6.6 × 10⁻⁹), and cilium movement (NES = −3.03, q-value = 6.6 × 10⁻⁹). These changes occurred alongside negative shifts in extracellular transport and transmitter-gated ion-channel activity, linking altered ciliary and microtubule organization with broader disruption of cochlear transport and signaling functions.

At the gene level, the omnibus likelihood-ratio test identified 15 genes with significant Genotype×Age effects at FDR < 0.05. These included *Egr3*, *Slain1*, *Sstr3*, *Adarb2*, *B3gnt3*, *Comp*, *Ppp1r17*, *Myh11*, *Shisa6*, *Caly*, and *Tmem132b*. The directional interaction tests identified only one significant gene at 8 months, four at 11 months, and one at 16 months. *Ighv2-3* showed a negative interaction at all three later ages, whereas *Gm14033* and *Tdo2* showed positive interactions specifically at 11 months; *Ighv12-3* showed a negative interaction at 11 months. The limited number of individually significant genes, despite extensive pathway-level enrichment, indicates that the age-dependent cochlear response was distributed across coordinated sets of moderately changing genes rather than being driven primarily by large effects in numerous individual genes.

Collectively, these findings reveal a nonmonotonic cochlear response to 5xFAD genotype. Extracellular-matrix, adhesion, and inflammatory programs predominated at 8 months; synaptic pathways shifted transiently toward positive enrichment at 11 months; and biosynthetic, mitochondrial, cell-cycle, and proteostasis programs became strongly activated at 16 months. In contrast, ciliary and axonemal pathways shifted negatively at both 11 and 16 months. This temporal sequence indicates that cochlear pathology does not progress through a single uniformly increasing transcriptional program but instead involves distinct, age-specific states affecting tissue remodeling, neuronal signaling, energy metabolism, protein homeostasis, and ciliary structure.

### Supplementary Tables

**Table S1.** Principal P50 transcriptional responses across auditory regions.

| Region | Significant DEGs | Dominant negatively enriched programs | Dominant positively enriched programs | Distinguishing regional feature |
| --- | --- | --- | --- | --- |
| Auditory cortex | 11 | Oxidative phosphorylation, carbon and lipid metabolism, ribosome, spliceosome, proteasome, protein folding | Synapse organization, neuronal projections, axon guidance, adhesion, ECM interaction, ABC transporters | Metabolic suppression coupled to neuronal and structural remodeling; strong Cst7 induction |
| Inferior colliculus | 2 | Oxidative phosphorylation, TCA cycle, glycolysis, fatty-acid metabolism, ribosome, spliceosome, proteasome, myelin-associated genes | Synapse organization, glutamatergic signaling, axon guidance, ECM, integrin and cadherin signaling, calcium signaling | Strongest metabolic suppression with extensive synaptic and extracellular remodeling |
| Cochlear nucleus | 2 | Synapse organization, axonal transport, receptor activity, cadherin, integrin and focal-adhesion signaling, inflammatory programs | Oxidative phosphorylation, TCA cycle, glycolysis, fatty-acid metabolism, ribosome, spliceosome, proteasome, MYC and mTORC1 programs | Coordinated reversal of the metabolic, biosynthetic, proteostatic, and myelin-associated programs |
| Cochlea | 2 | Oxidative phosphorylation, carbon metabolism, ribosome, proteasome, synaptic-vesicle cycling, glutamatergic and GABAergic pathways | ECM, collagen, angiogenesis, cilium and stereocilium organization, hair-cell differentiation, developmental signaling | Peripheral sensory-structure and vascular remodeling accompanying metabolic and synaptic suppression |

**Table S2.** Representative P50 pathway enrichment across auditory regions. Values are NES with adjusted P values in parentheses. A dash indicates that the pathway was absent from thesignificant results.

| Pathway | Auditory cortex | Inferior colliculus | Cochlear nucleus | Cochlea |
| --- | --- | --- | --- | --- |
| Hallmark oxidative phosphorylation | -2.765 ($5.62 \times 10^{-17}$) | -3.383 ($1.72 \times 10^{-32}$) | 2.355 ($1.27 \times 10^{-11}$) | -2.666 ($6.73 \times 10^{-16}$) |
| Hallmark MYC targets V1 | -2.509 ($3.31 \times 10^{-13}$) | -3.085 ($7.59 \times 10^{-26}$) | 2.383 ($5.78 \times 10^{-12}$) | -2.856 ($3.23 \times 10^{-21}$) |
| KEGG carbon metabolism | -2.063 ($4.29 \times 10^{-6}$) | -2.520 ($2.37 \times 10^{-10}$) | 2.346 ($1.51 \times 10^{-9}$) | -2.254 ($2.05 \times 10^{-8}$) |
| KEGG ribosome | -2.650 ($1.00 \times 10^{-10}$) | -3.052 ($1.00 \times 10^{-10}$) | 2.513 ($1.00 \times 10^{-10}$) | -2.019 ($4.93 \times 10^{-8}$) |
| KEGG proteasome | -2.261 ($6.59 \times 10^{-6}$) | -2.425 ($2.73 \times 10^{-7}$) | 1.875 ($4.81 \times 10^{-4}$) | -2.311 ($5.07 \times 10^{-7}$) |
| GO regulation of synapse structure or activity | 1.864 ($4.37 \times 10^{-9}$) | 1.934 ($1.90 \times 10^{-10}$) | -1.470 ($2.78 \times 10^{-4}$) | -1.519 ($2.28 \times 10^{-5}$) |
| GO neuron-projection organization | 1.958 ($1.20 \times 10^{-5}$) | 1.821 ($5.30 \times 10^{-5}$) | -1.374 (0.0287) | -1.748 ($2.73 \times 10^{-4}$) |
| KEGG axon guidance | 1.785 ($2.44 \times 10^{-5}$) | 1.878 ($2.43 \times 10^{-6}$) | -1.404 (0.0122) | — |
| GO homophilic cell adhesion | 2.713 ($1.00 \times 10^{-10}$) | 2.701 ($1.00 \times 10^{-10}$) | -2.069 ($6.27 \times 10^{-8}$) | — |
| GO myelin sheath | -2.035 ($3.01 \times 10^{-8}$) | -2.770 ($1.00 \times 10^{-10}$) | 1.895 ($7.37 \times 10^{-7}$) | -2.117 ($6.32 \times 10^{-10}$) |
| GO cilium movement | — | 1.592 ($4.22 \times 10^{-4}$) | -1.410 (0.00401) | 1.971 ($2.72 \times 10^{-8}$) |

**Table S3.** Representative auditory-cortex age-specific and interaction NES values. Asterisks mark BH-adjusted P < 0.05 for Hallmark or q-value < 0.05 for KEGG and GO. Interaction NES describes the change in the genotype effect relative to P50.

| Pathway | P50 | 8 months | 11 months | 16 months | 8 months vs P50 | 11 months vs P50 | 16 months vs P50 |
| --- | --- | --- | --- | --- | --- | --- | --- |
| Interferon-γ response | -0.99 | +2.67* | +3.18* | +2.76* | +2.68* | +2.80* | +2.50* |
| Inflammatory response | -0.83 | +2.63* | +2.51* | +2.44* | +2.56* | +2.13* | +2.09* |
| ECM/remodeling (EMT gene set) | -1.28 | +1.79* | +2.69* | +2.49* | +2.07* | +2.46* | +2.55* |
| Oxidative phosphorylation | -2.76* | -1.86* | +1.14 | -1.47* | +1.44* | +2.23* | +1.52* |
| MYC Targets V1 | -2.51* | -0.81 | +1.92* | -1.57* | +1.63* | +2.43* | +1.38* |
| Mitotic spindle | +2.19* | +1.22 | -1.20 | +0.69 | -1.57* | -1.93* | -1.96* |
| Ribosome | -2.65* | +2.23* | +2.50* | +1.27 | +2.70* | +2.85* | +2.44* |
| Phagocytosis | -1.55* | +2.31* | +2.56* | +2.25* | +2.50* | +2.54* | +2.17* |
| Phosphatidylinositol signaling | +2.16* | -1.04 | -1.65* | -1.25 | -1.90* | -2.05* | -2.12* |
| Axon guidance | +1.79* | +0.92 | -1.37* | -1.14 | -1.33* | -1.67* | -1.72* |
| Glutamatergic synapse | +1.37 | +0.94 | -1.76* | -1.25 | -1.15 | -1.78* | -1.47* |

**Table S4.** Cross-region classification summary. Broadly shared includes conserved pathways. . Divergent indicates at least one significant positive and one significant negative interaction among regions at the same age.

| Database | Age contrast | Conserved | Broadly shared | Divergent |
| --- | --- | --- | --- | --- |
| Hallmark | 8 months | 8 | 15 | 1 |
| Hallmark | 11 months | 0 | 3 | 14 |
| Hallmark | 16 months | 8 | 23 | 13 |
| KEGG | 8 months | 7 | 34 | 11 |
| KEGG | 11 months | 0 | 10 | 62 |
| KEGG | 16 months | 12 | 71 | 70 |
| GO | 8 months | 2 | 16 | 2 |
| GO | 11 months | 0 | 7 | 42 |
| GO | 16 months | 6 | 34 | 40 |

### Supplementary Figure Caption

**Figure S1.**
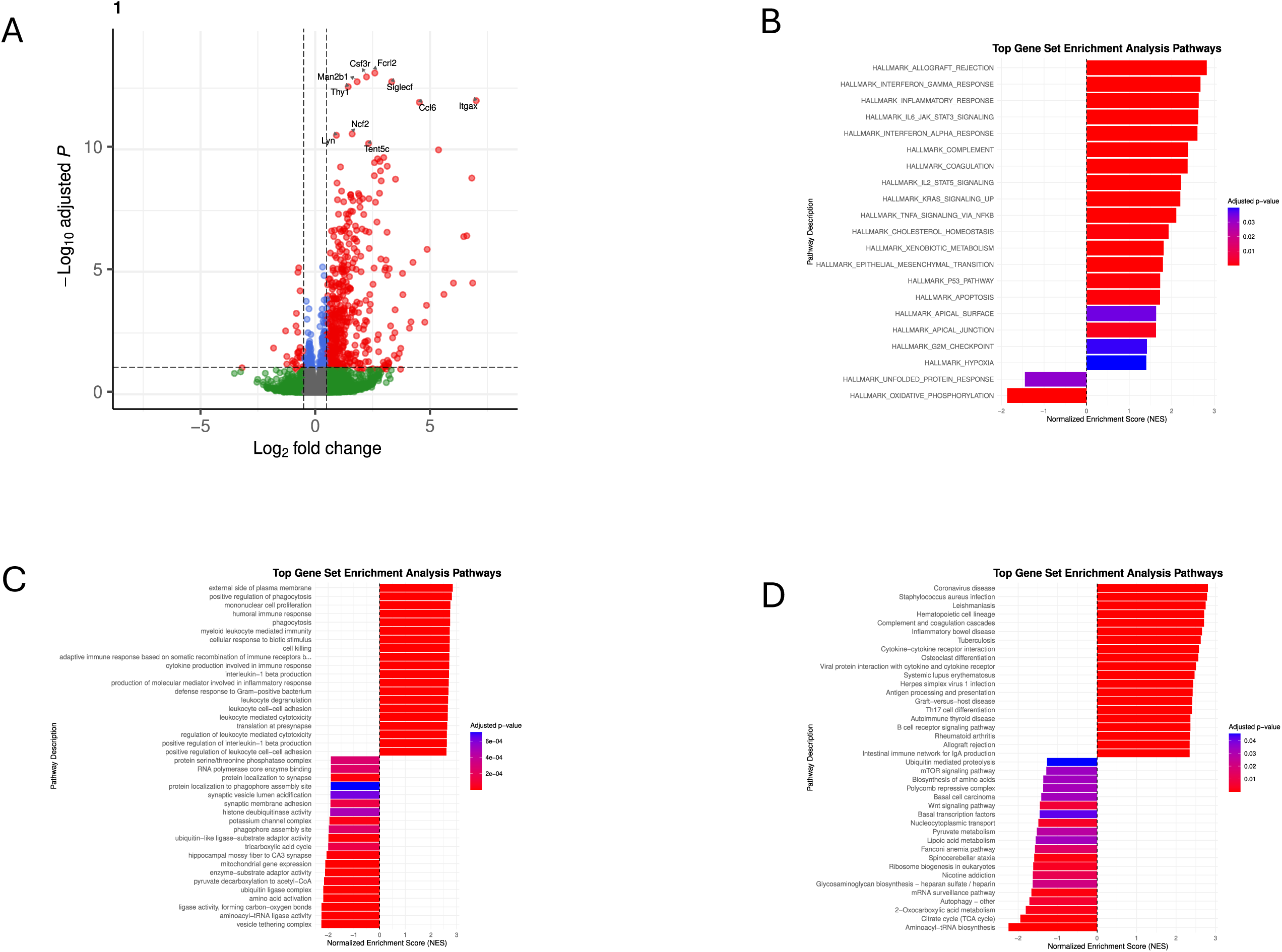
Age-specific transcriptomic alterations in the auditory cortex at 8-month. (A) Volcano plot showing gene-level differential expression between 5xFAD and wild-type (WT) mice. (B–D) Gene set enrichment analysis (GSEA) of (B) Hallmark, (C) Gene Ontology (GO), and (D) Kyoto Encyclopedia of Genes and Genomes (KEGG) pathways. For each gene-set collection, the 20 most positively enriched and 20 most negatively enriched pathways, ranked by normalized enrichment score (NES), are shown. Positive NES values indicate enrichment in 5xFAD mice, whereas negative NES values indicate enrichment in WT mice.

**Figure S2.**
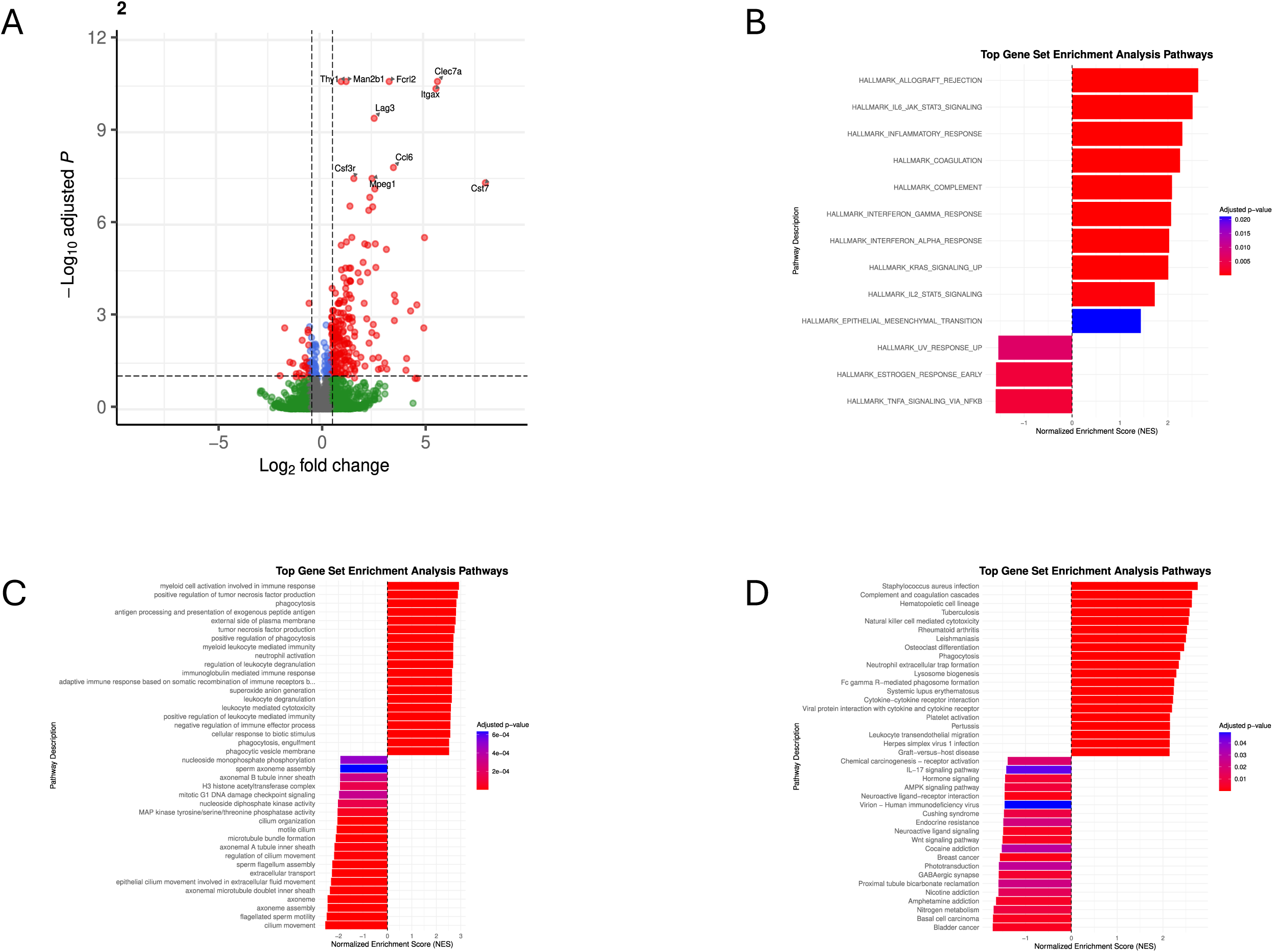
Age-specific transcriptomic alterations in the inferior colliculus at 8-onth. (A) Volcano plot showing gene-level differential expression between 5xFAD and wild-type (WT) mice. (B–D) Gene set enrichment analysis (GSEA) of (B) Hallmark, (C) Gene Ontology (GO), and (D) Kyoto Encyclopedia of Genes and Genomes (KEGG) pathways. For each gene-set collection, the 20 most positively enriched and 20 most negatively enriched pathways, ranked by normalized enrichment score (NES), are shown. NES directionality is defined as in Figure 2.

**Figure S3.**
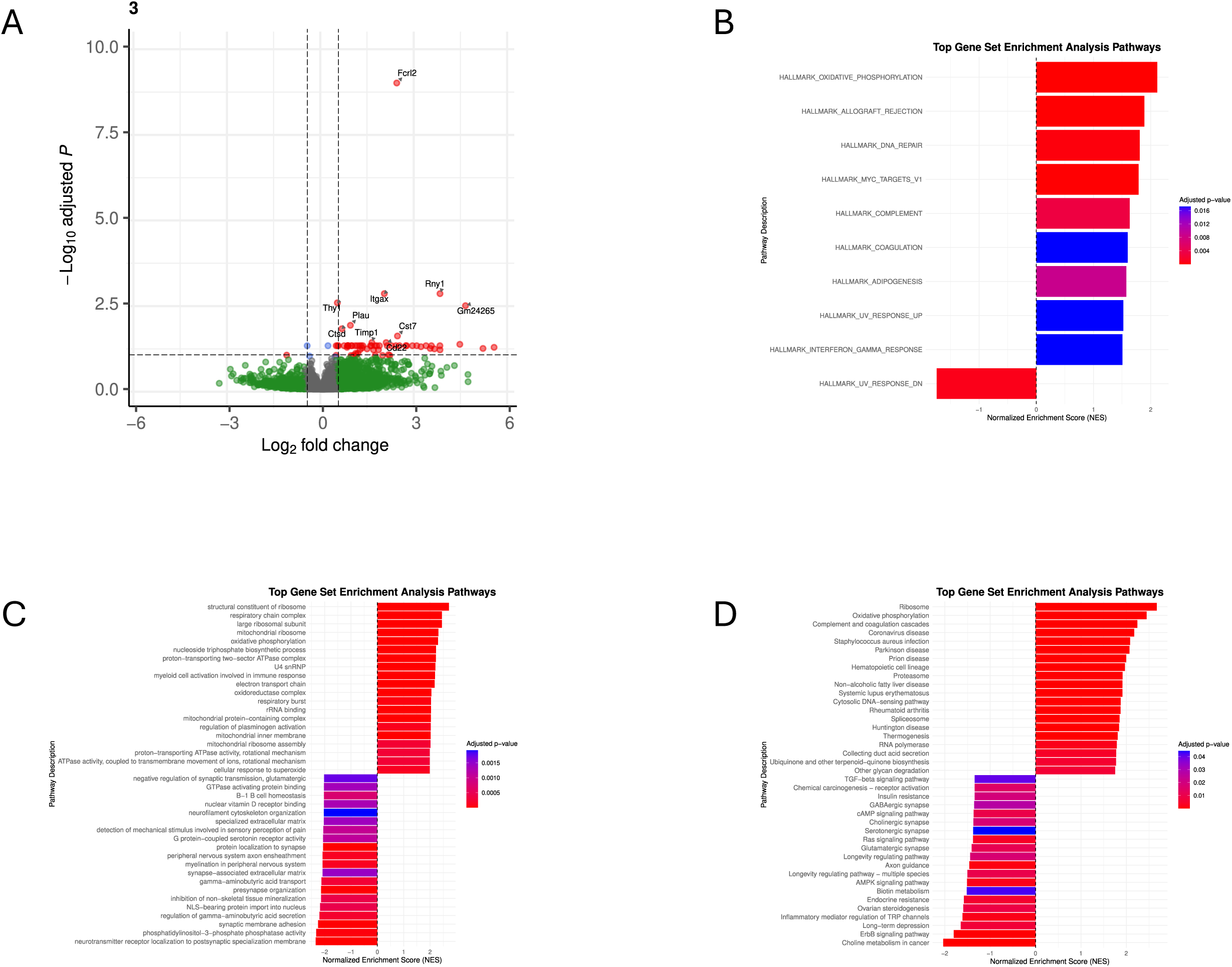
Age-specific transcriptomic alterations in the cochlear nucleus at 8-month. (A) Volcano plot showing gene-level differential expression between 5xFAD and wild-type (WT) mice. (B–D) Gene set enrichment analysis (GSEA) of (B) Hallmark, (C) Gene Ontology (GO), and (D) Kyoto Encyclopedia of Genes and Genomes (KEGG) pathways. For each gene-set collection, the 20 most positively enriched and 20 most negatively enriched pathways, ranked by normalized enrichment score (NES), are shown. NES directionality is defined as in Figure 2.

**Figure S4.**
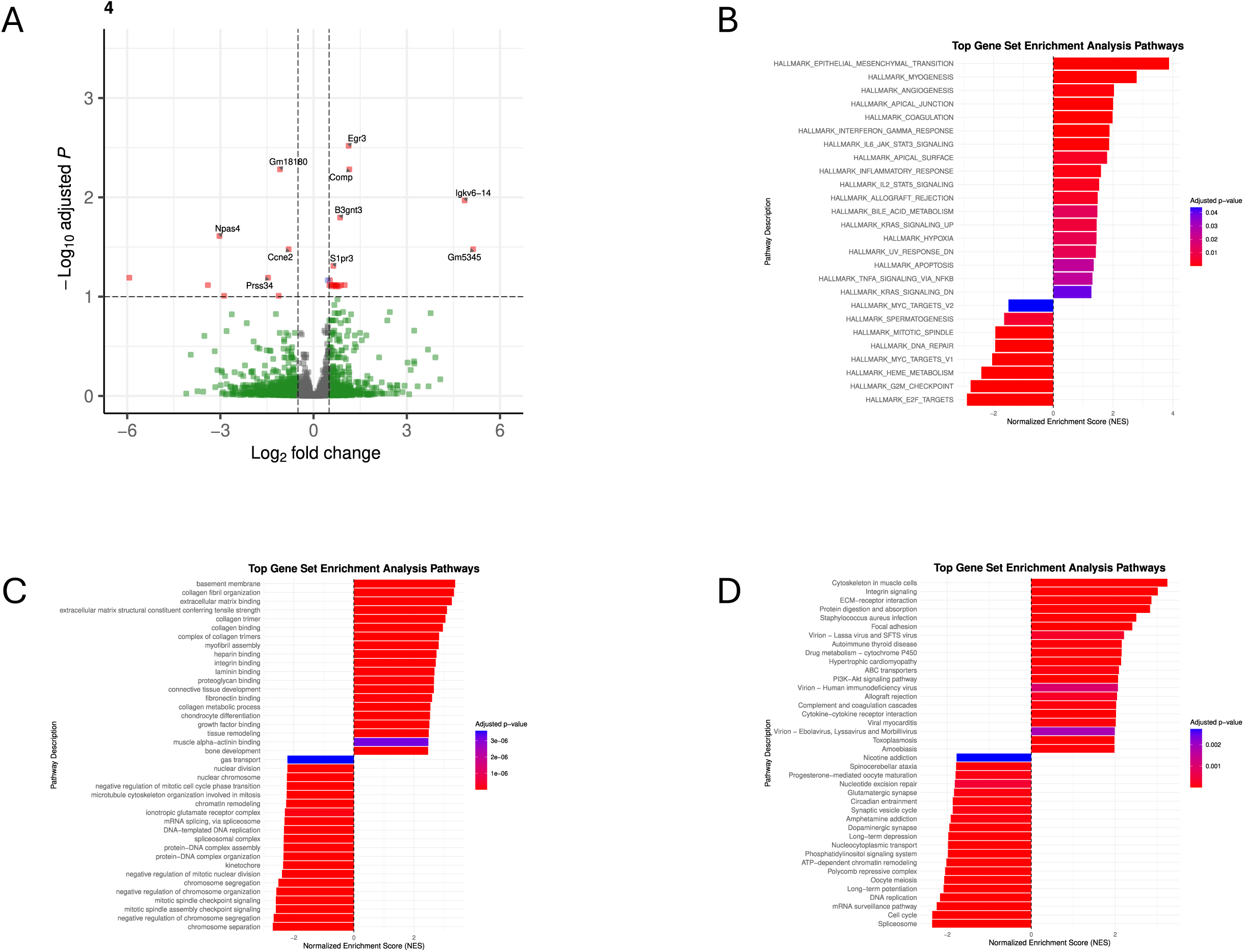
Age-specific transcriptomic alterations in the cochlea at 8-month. (A) Volcano plot showing gene-level differential expression between 5xFAD and wild-type (WT) mice. (B–D) Gene set enrichment analysis (GSEA) of (B) Hallmark, (C) Gene Ontology (GO), and (D) Kyoto Encyclopedia of Genes and Genomes (KEGG) pathways. For each gene-set collection, the 20 most positively enriched and 20 most negatively enriched pathways, ranked by normalized enrichment score (NES), are shown. NES directionality is defined as in Figure 2.

**Figure S5.**
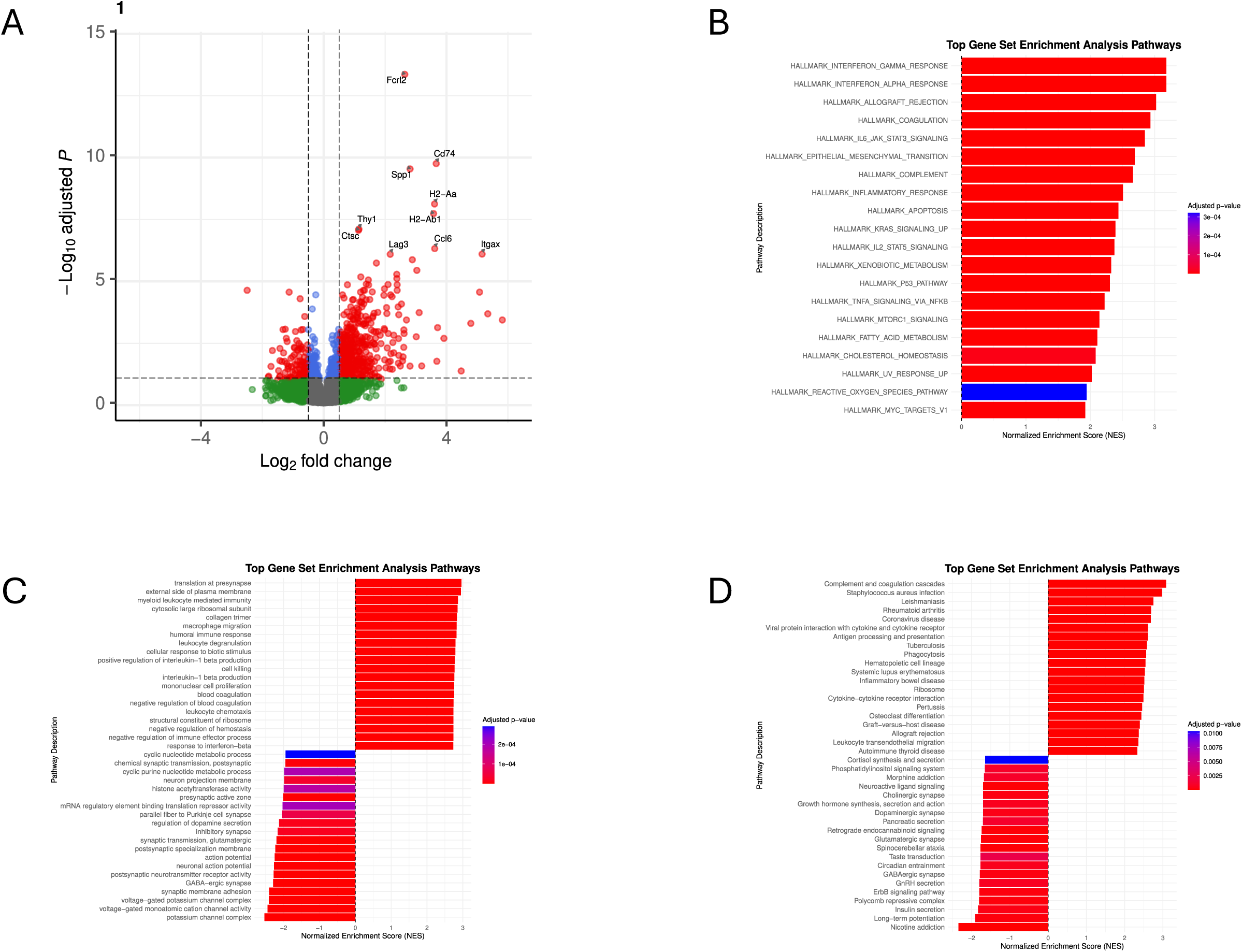
Age-specific transcriptomic alterations in the auditory cortex at 11-month. (A) Volcano plot showing gene-level differential expression between 5xFAD and wild-type (WT) mice. (B–D) Gene set enrichment analysis (GSEA) of (B) Hallmark, (C) Gene Ontology (GO), and (D) Kyoto Encyclopedia of Genes and Genomes (KEGG) pathways. For each gene-set collection, the 20 most positively enriched and 20 most negatively enriched pathways, ranked by normalized enrichment score (NES), are shown.Positive NES values indicate enrichment in 5xFAD mice, whereas negative NES values indicate enrichment in WT mice.

**Figure S6.**
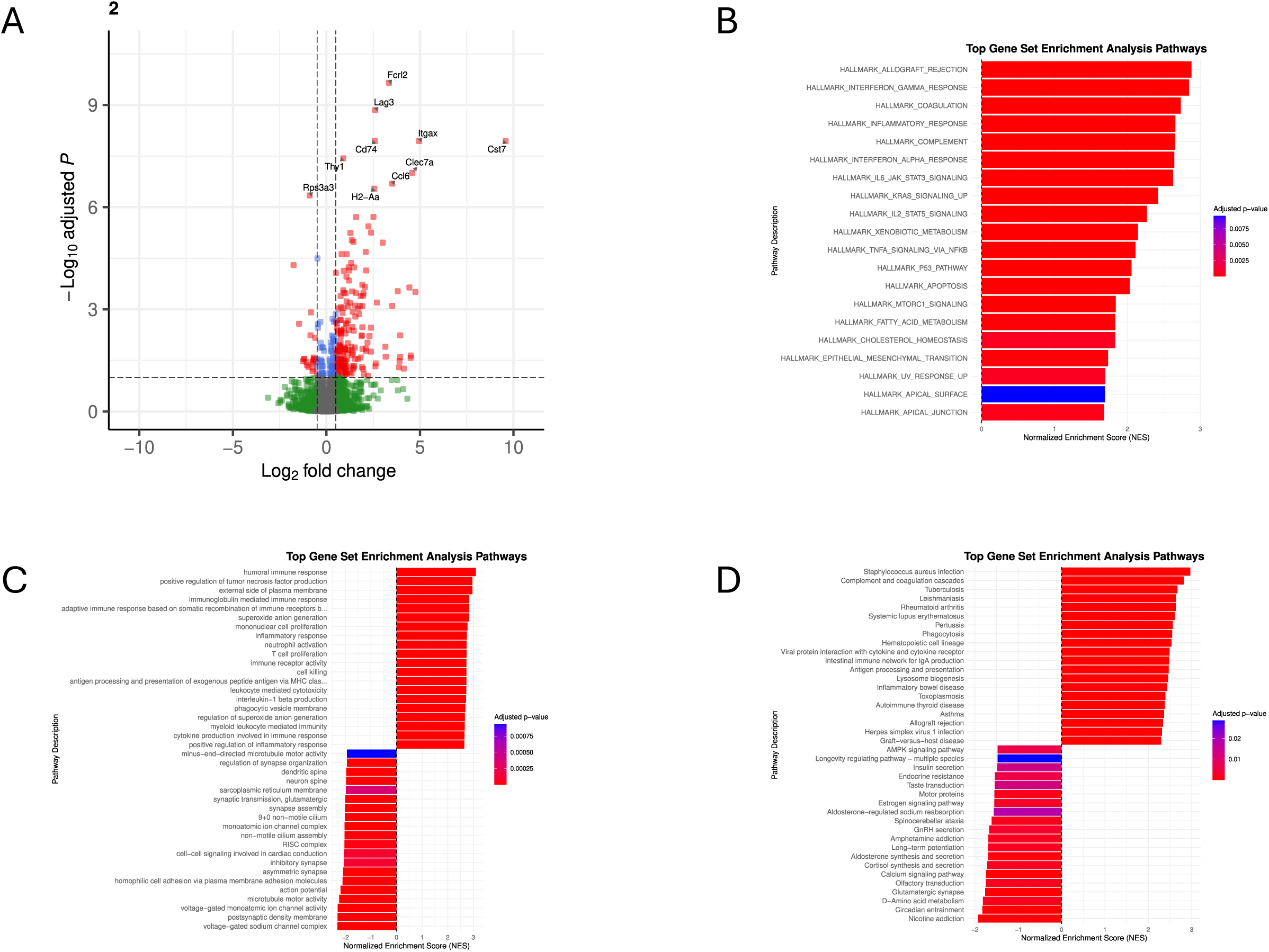
Age-specific transcriptomic alterations in the inferior colliculus at 11-onth. (A) Volcano plot showing gene-level differential expression between 5xFAD and wild-type (WT) mice. (B–D) Gene set enrichment analysis (GSEA) of (B) Hallmark, (C) Gene Ontology (GO), and (D) Kyoto Encyclopedia of Genes and Genomes (KEGG) pathways. For each gene-set collection, the 20 most positively enriched and 20 most negatively enriched pathways, ranked by normalized enrichment score (NES), are shown. NES directionality is defined as in Figure 2.

**Figure S7.**
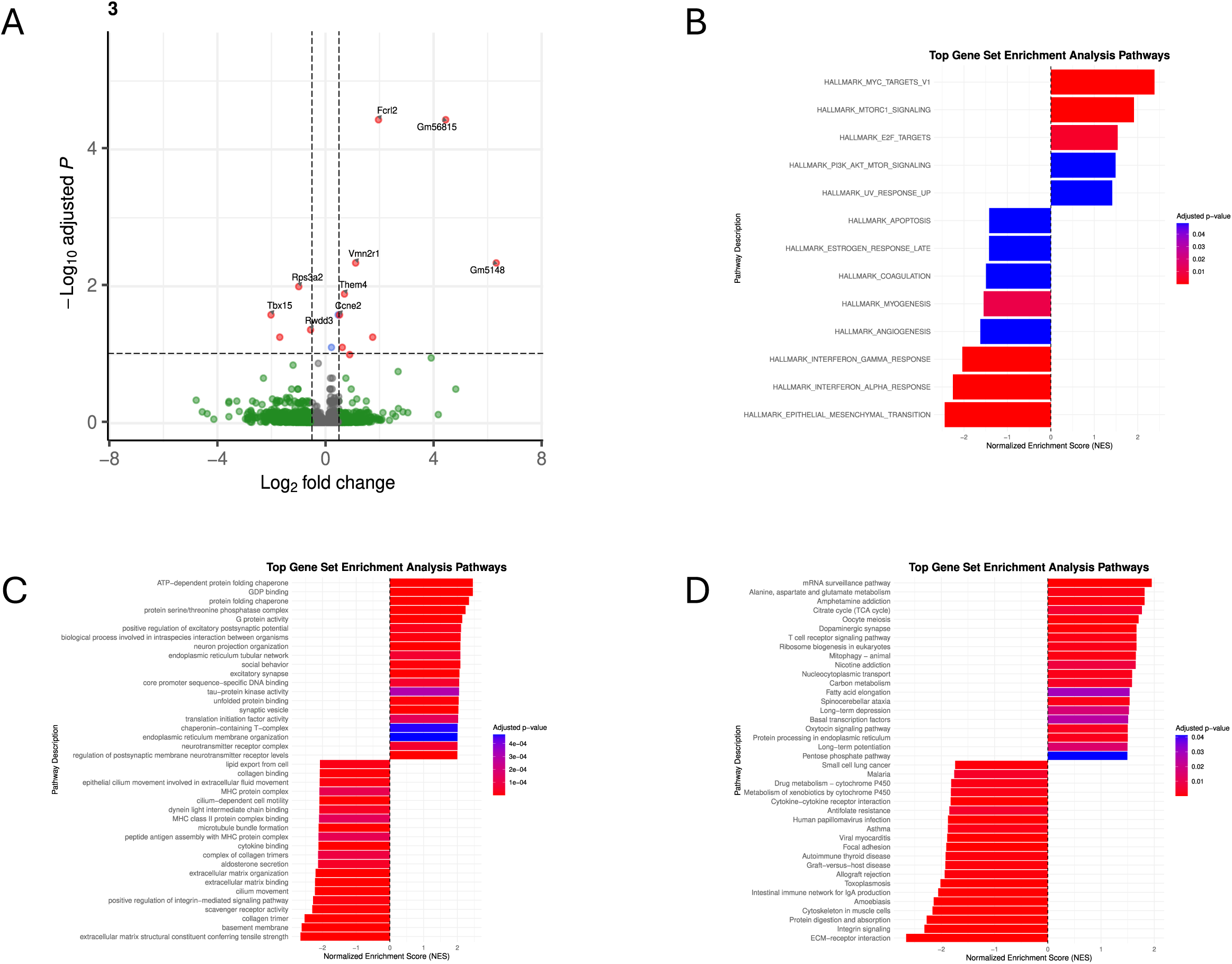
Age-specific transcriptomic alterations in the cochlear nucleus at 11-month. (A) Volcano plot showing gene-level differential expression between 5xFAD and wild-type (WT) mice. (B–D) Gene set enrichment analysis (GSEA) of (B) Hallmark, (C) Gene Ontology (GO), and (D) Kyoto Encyclopedia of Genes and Genomes (KEGG) pathways. For each gene-set collection, the 20 most positively enriched and 20 most negatively enriched pathways, ranked by normalized enrichment score (NES), are shown. NES directionality is defined as in Figure 2.

**Figure S8.**
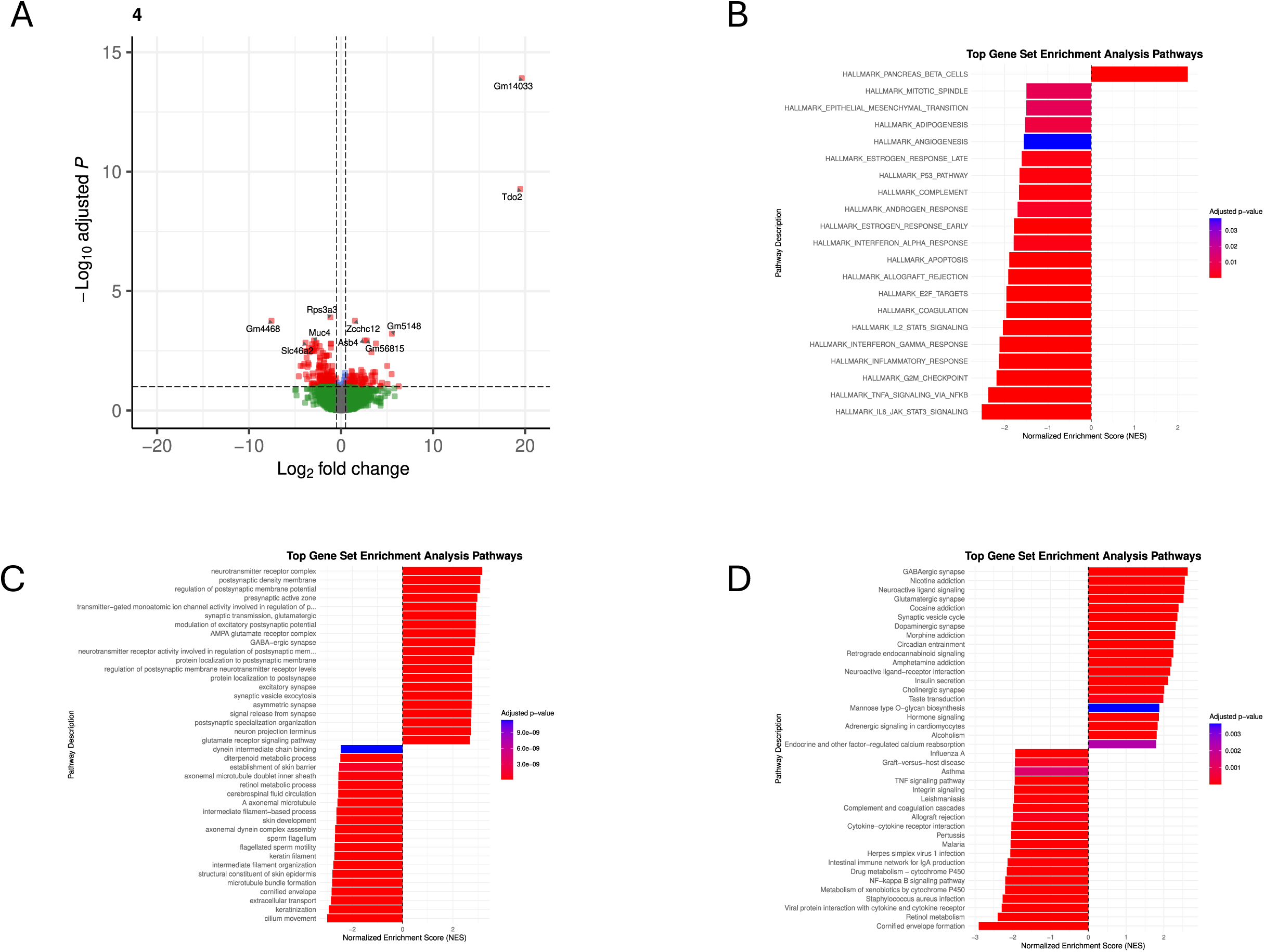
Age-specific transcriptomic alterations in the cochlea at 11-month. (A) Volcano plot showing gene-level differential expression between 5xFAD and wild-type (WT) mice. (B–D) Gene set enrichment analysis (GSEA) of (B) Hallmark, (C) Gene Ontology (GO), and (D) Kyoto Encyclopedia of Genes and Genomes (KEGG) pathways. For each gene-set collection, the 20 most positively enriched and 20 most negatively enriched pathways, ranked by normalized enrichment score (NES), are shown. NES directionality is defined as in Figure 2.

**Figure S9.**
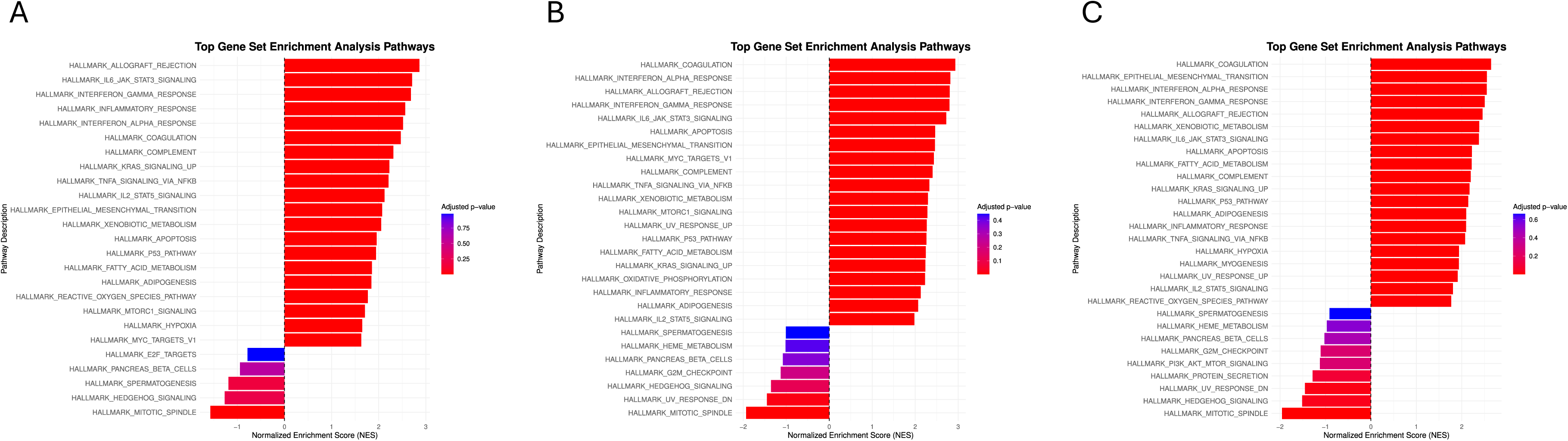
Age-dependent changes in Hallmark pathway enrichment in the auditory cortex relative to P50. Heatmaps show interaction normalized enrichment scores (NES) from Genotype × Age analyses comparing the 5xFAD-versus-WT effect at (A) 8 months, (B) 11 months, and (C) 16 months with the corresponding genotype effect at P50. For each comparison, the 20 pathways with the highest positive interaction NES and the 20 pathways with the strongest negative interaction NES are shown. A positive interaction NES indicates that the 5xFAD-versus-WT effect shifted in a positive direction relative to P50, whereas a negative interaction NES indicates a shift in the negative direction. Interaction NES values describe changes from the P50 genotype effect and do not necessarily indicate the direction of enrichment at the later age.

**Figure S10.**
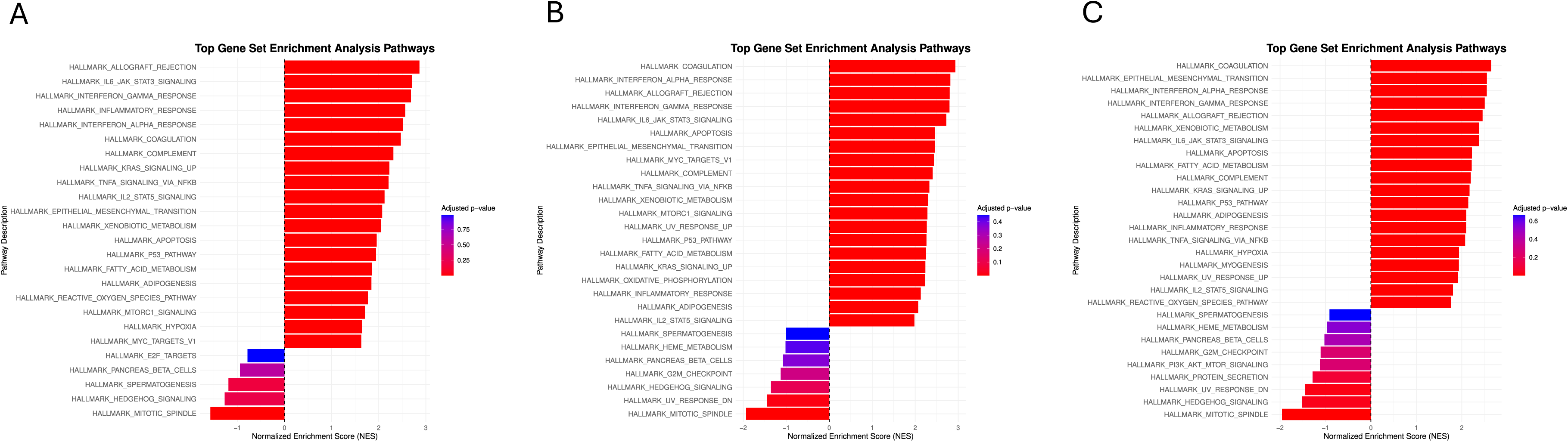
Age-dependent changes in GO pathway enrichment in the auditory cortex relative to P50.

**Figure S11.**
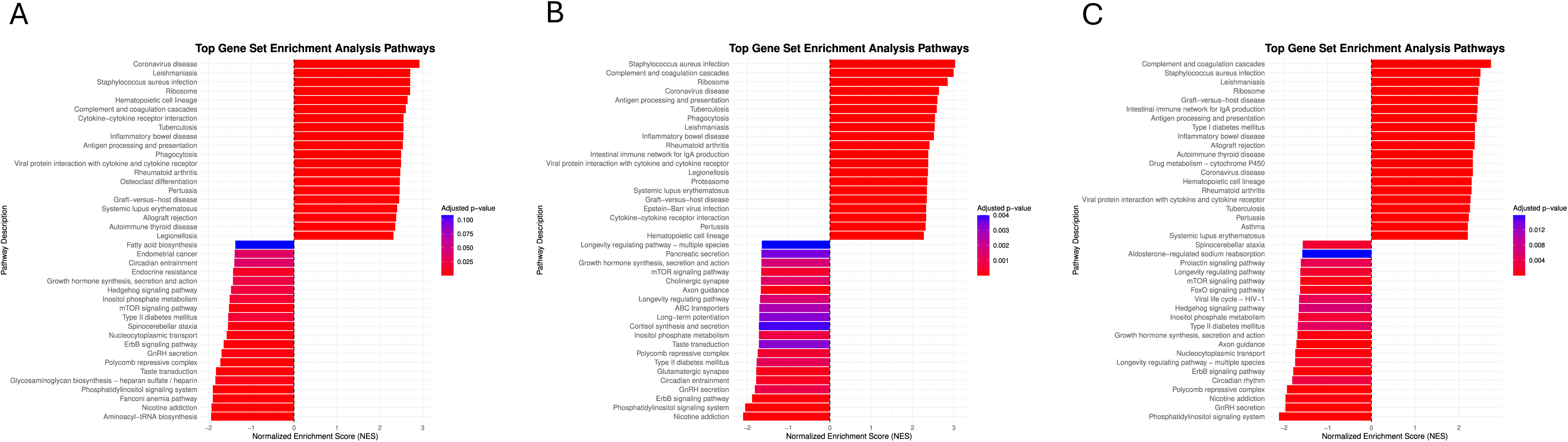
Age-dependent changes in KEGG pathway enrichment in the auditory cortex relative to P50.

**Figure S12.**
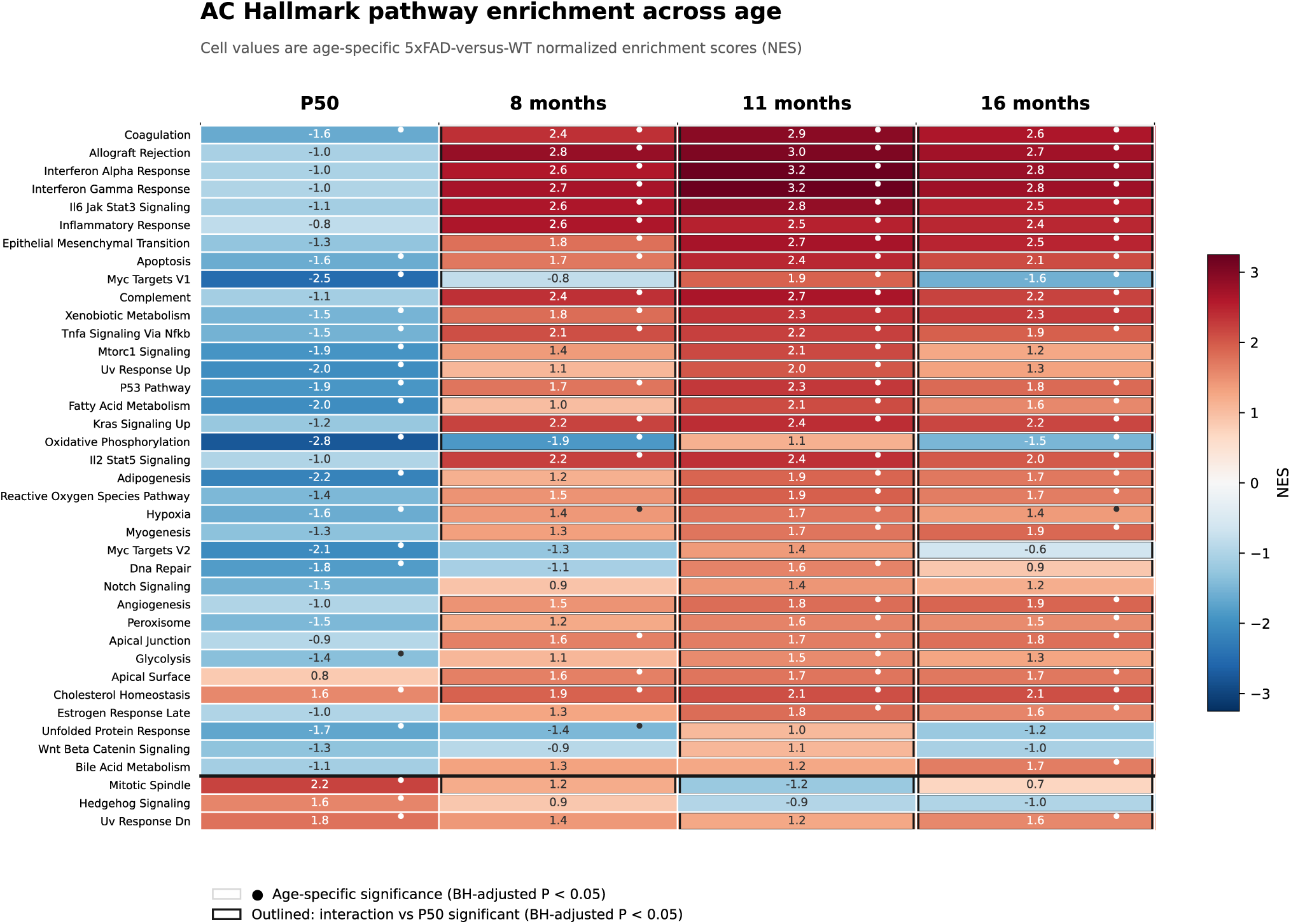
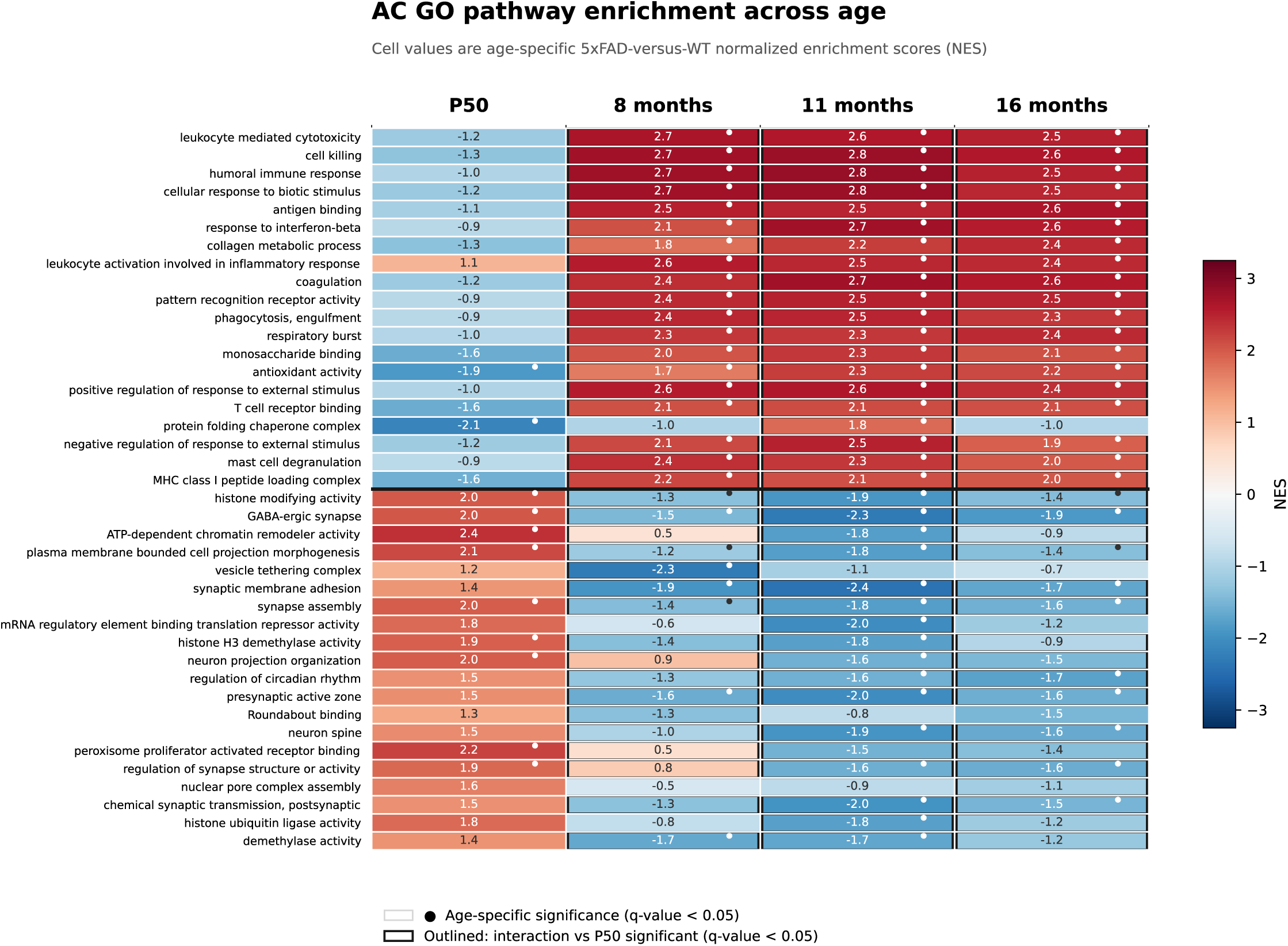
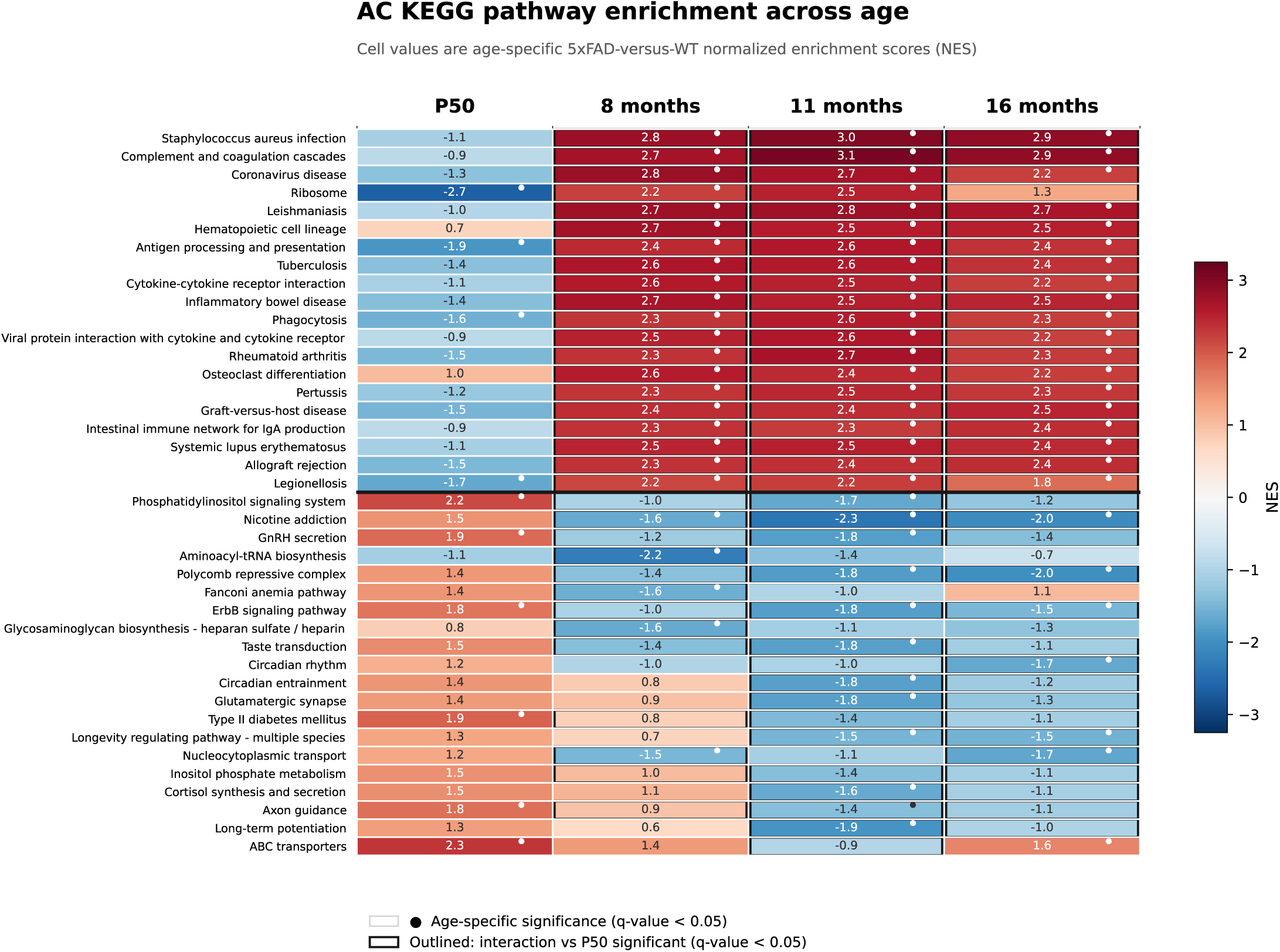
Age-dependent pathway enrichment in auditory cortex. Heatmaps show age-specific 5xFAD-versus-WT NES at P50, 8, 11, and 16 months. Hallmark includes pathways with a significant Genotype×Age interaction; KEGG and GO show the 20 strongest predominantly positive and 20 strongest predominantly negative interaction shifts. Dots mark significant age-specific enrichment and black outlines mark significant interactions relative to P50. Red and blue denote positive and negative age-specific enrichment. Interaction outlines indicate change from P50 and do not independently specify later-age enrichment. Significance was BH-adjusted P < 0.05 for Hallmark and q-value < 0.05 for KEGG and GO.

**Figure S13.**
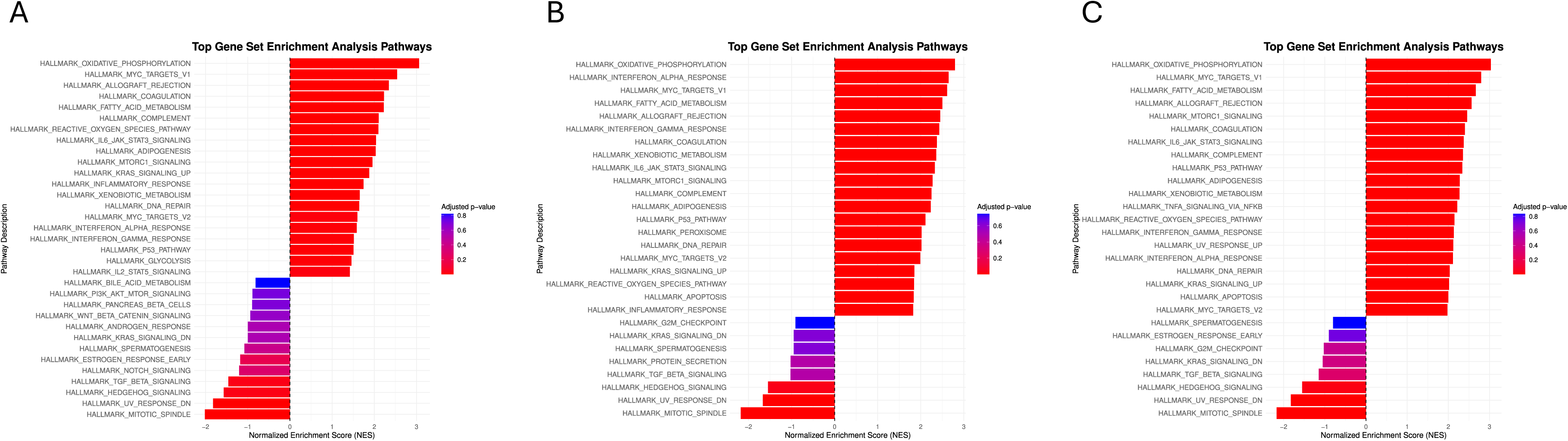
Age-dependent changes in Hallmark pathway enrichment in the inferior colliculus relative to P50.

**Figure S14.**
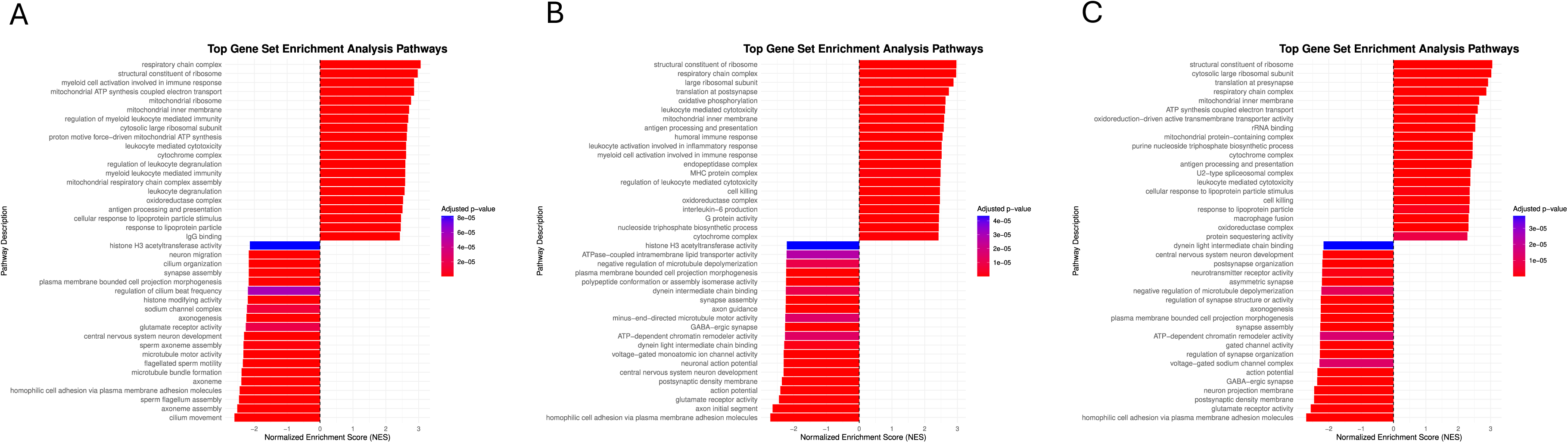
Age-dependent changes in GO pathway enrichment in the inferior colliculus relative to P50.

**Figure S15.**
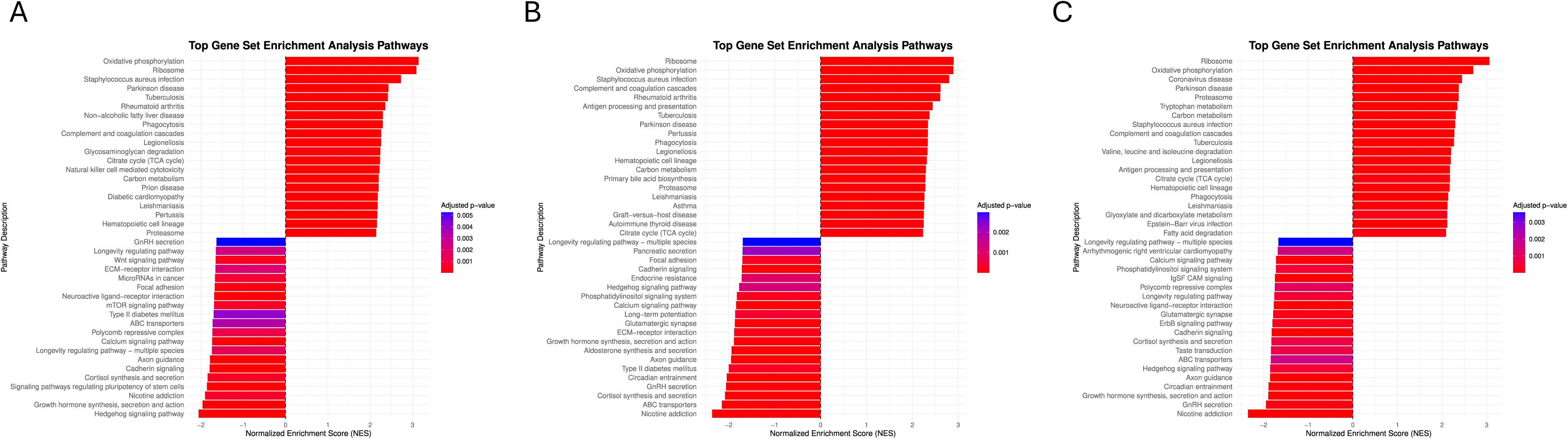
Age-dependent changes in KEGG pathway enrichment in the inferior colliculus relative to P50.

**Figure S16.**
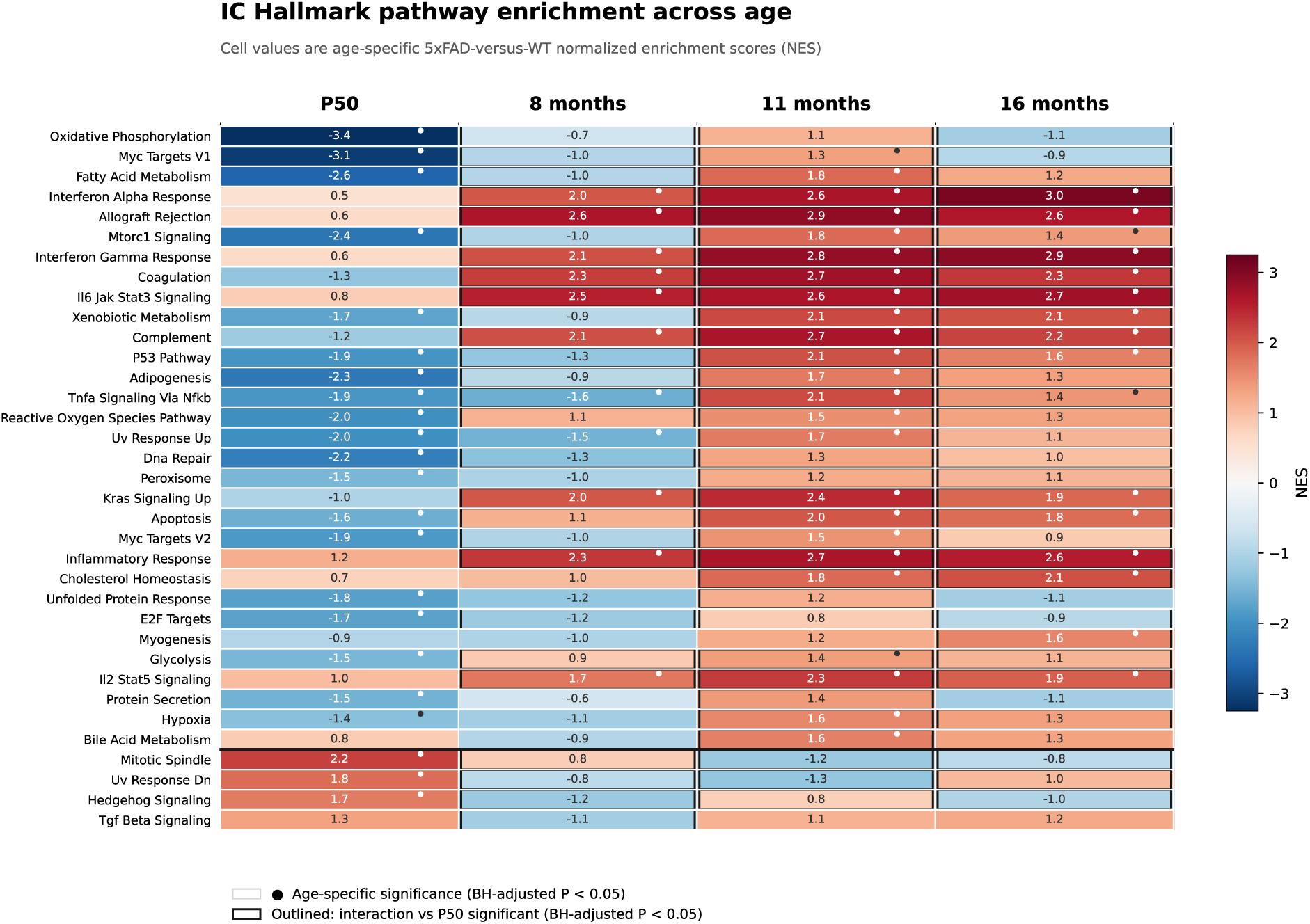
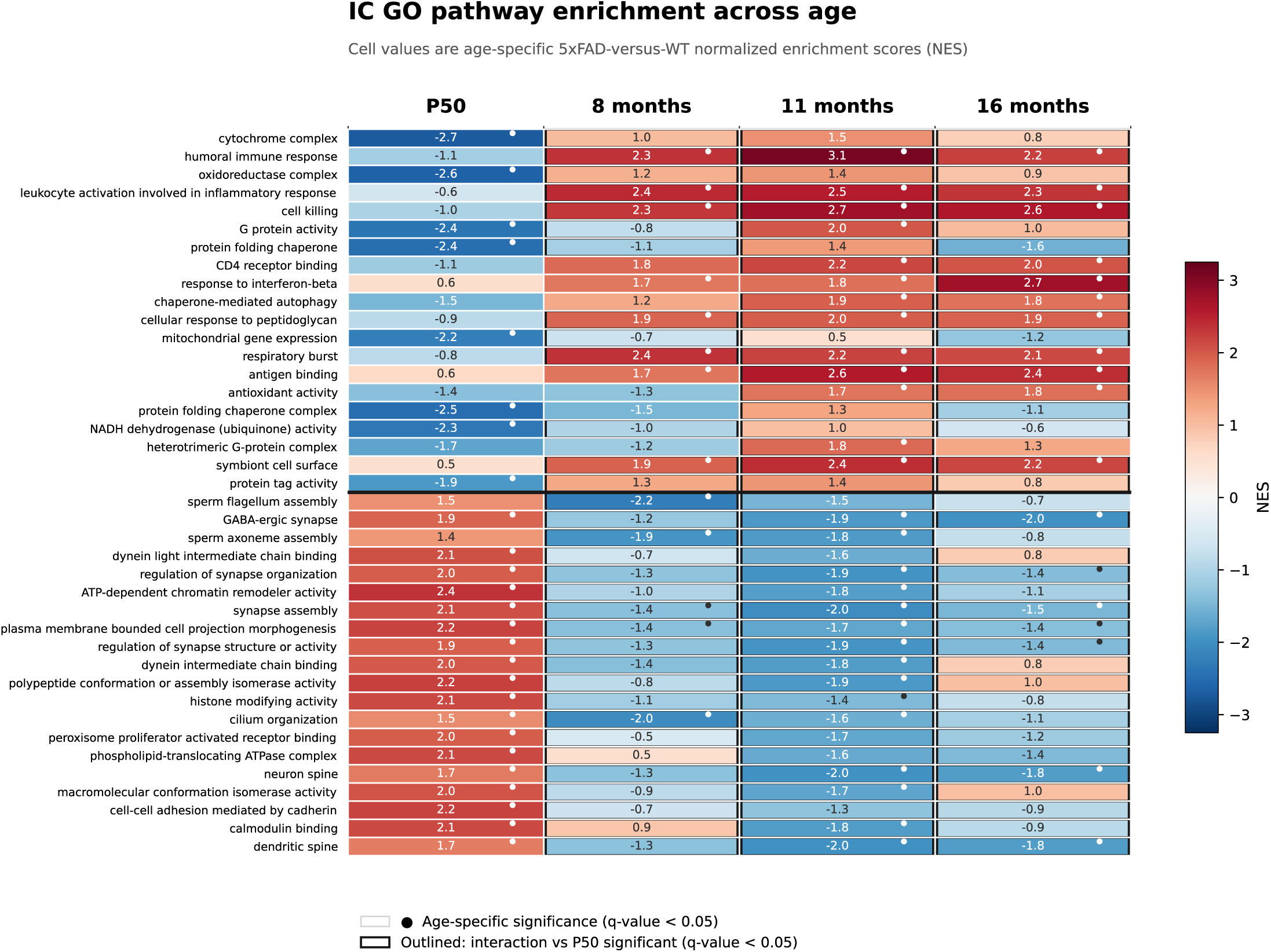
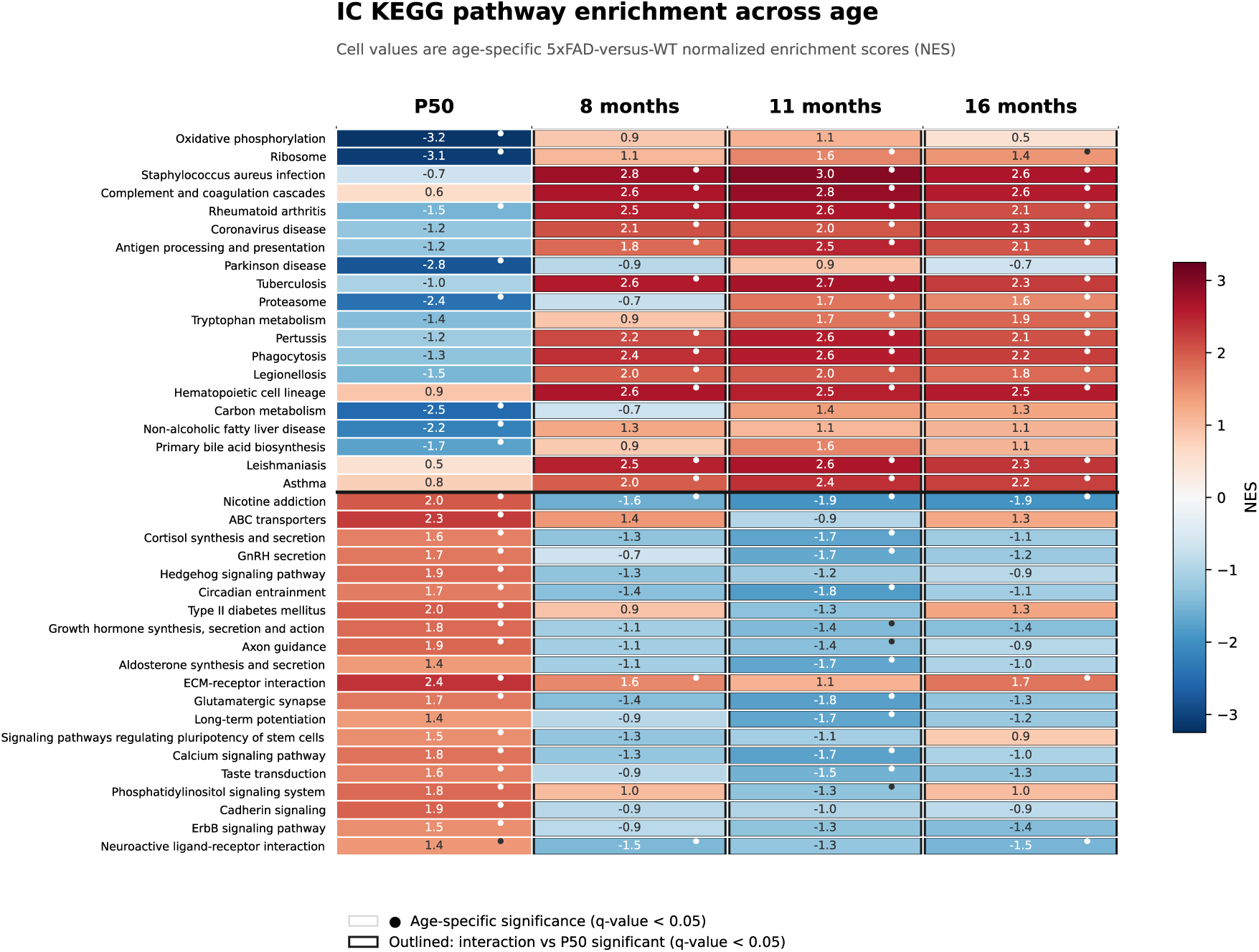
Age-dependent pathway enrichment in inferior colliculus. Display, selection, and significance conventions are as in Figure S12. Columns show P50, 8, 11, and 16 months; dots identify significant age-specific enrichment and black outlines identify significant Genotype×Age interactions relative to P50.

**Figure S17.**
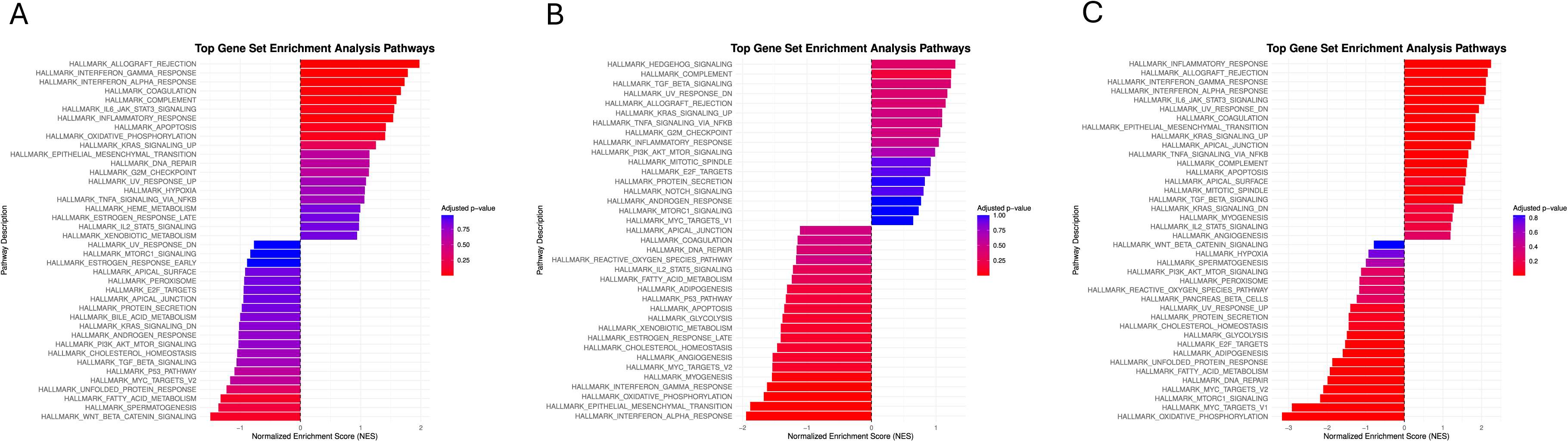
Age-dependent changes in Hallmark pathway enrichment in the cochlear nucleus relative to P50.

**Figure S18.**
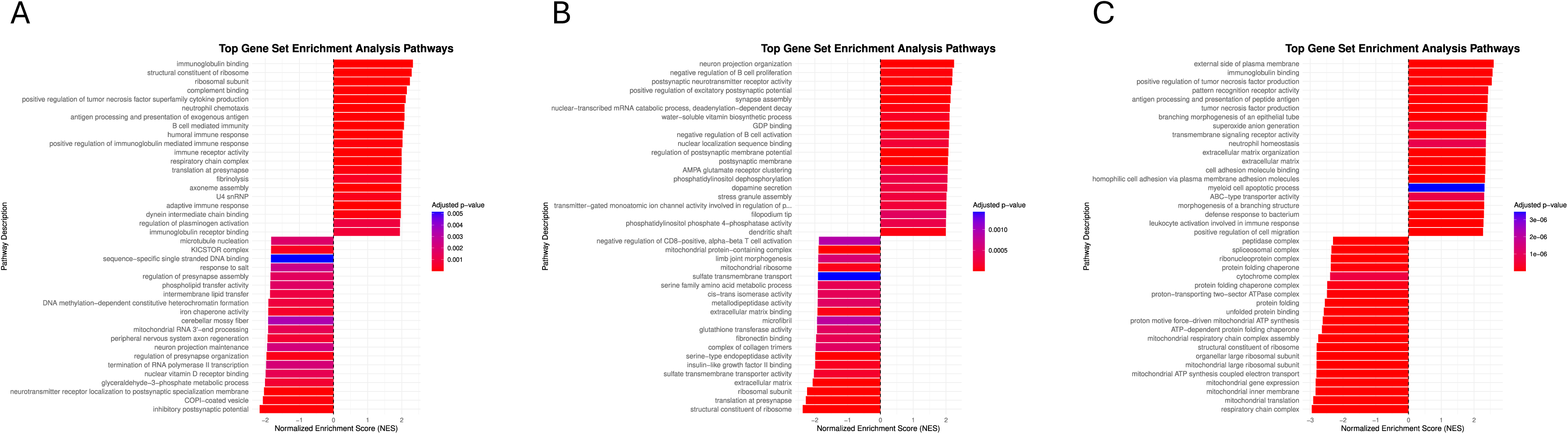
Age-dependent changes in GO pathway enrichment in the cochlear nucleus relative to P50.

**Figure S19.**
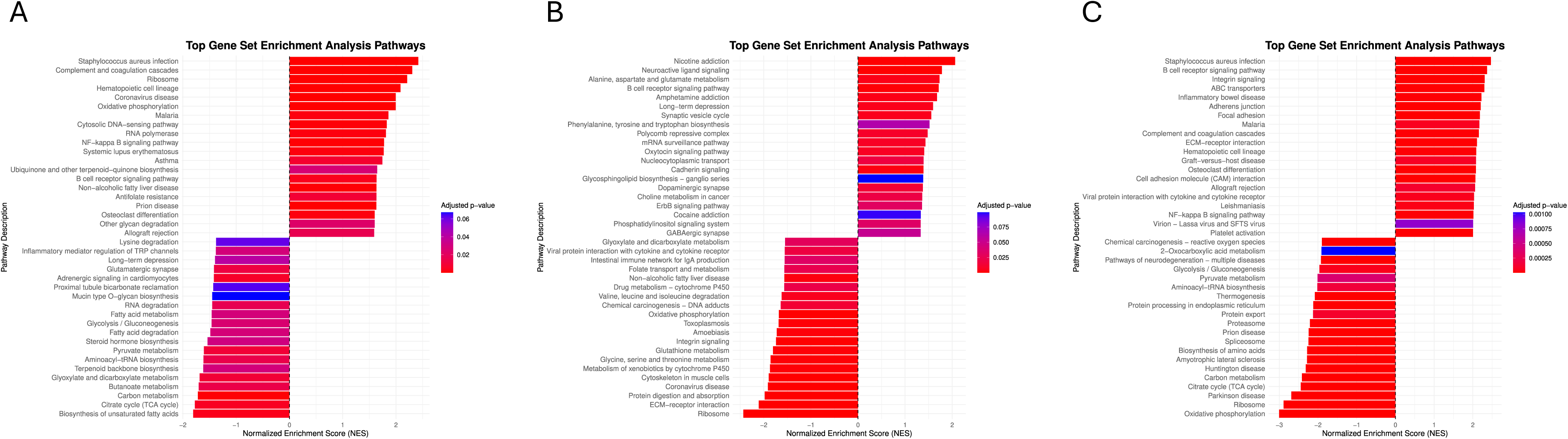
Age-dependent changes in KEGG pathway enrichment in the cochlear nucleus relative to P50.

**Figure S20.**
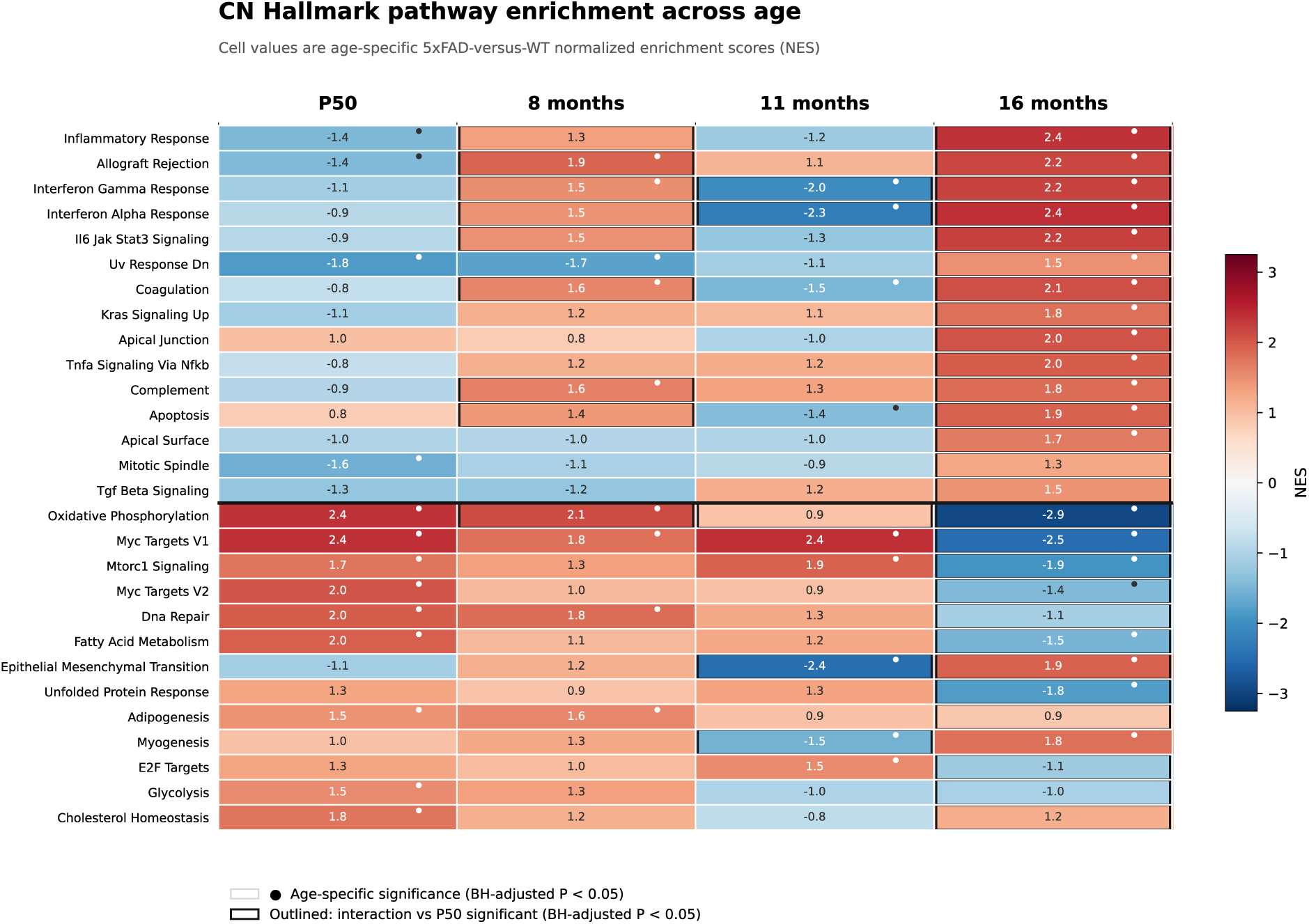
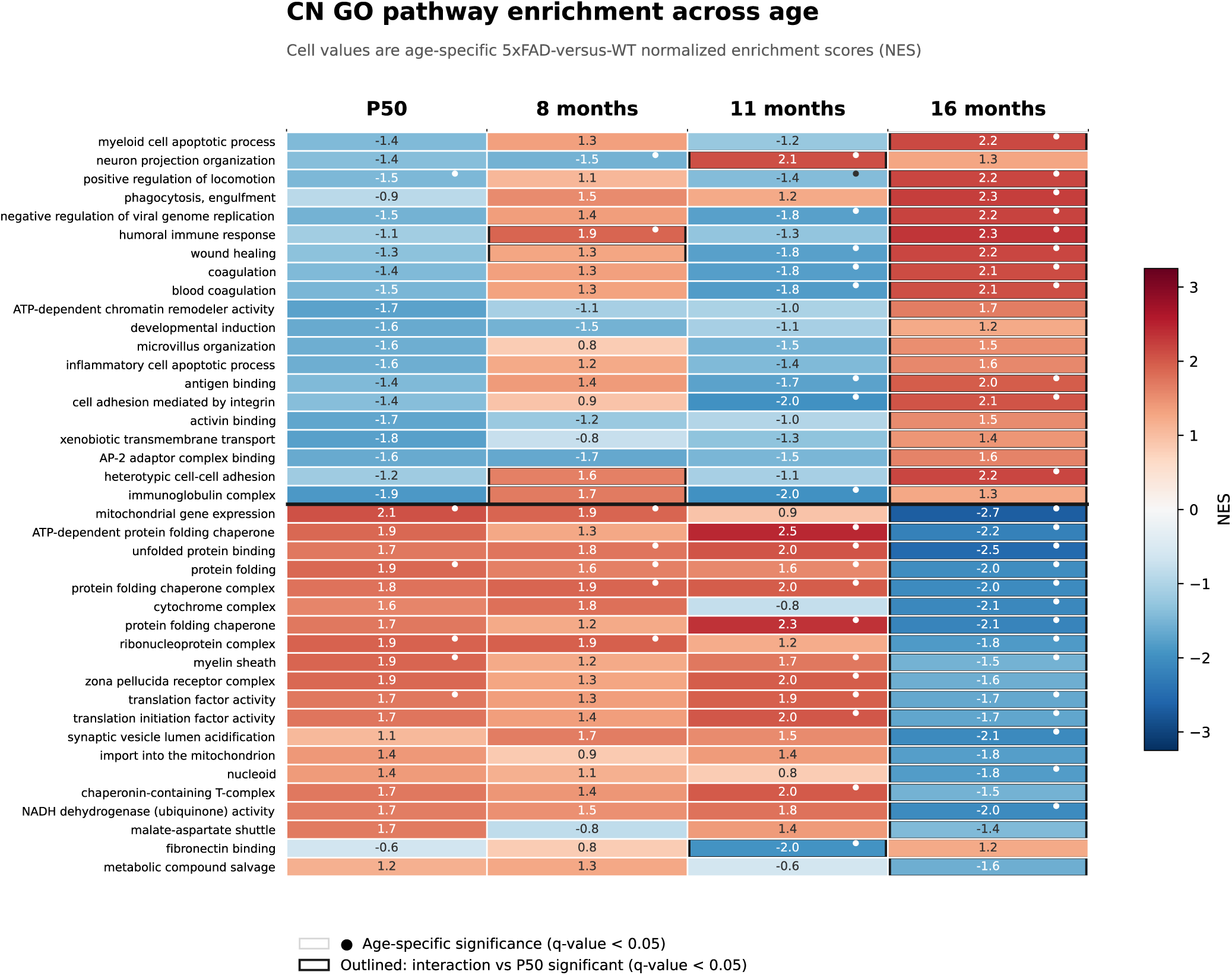
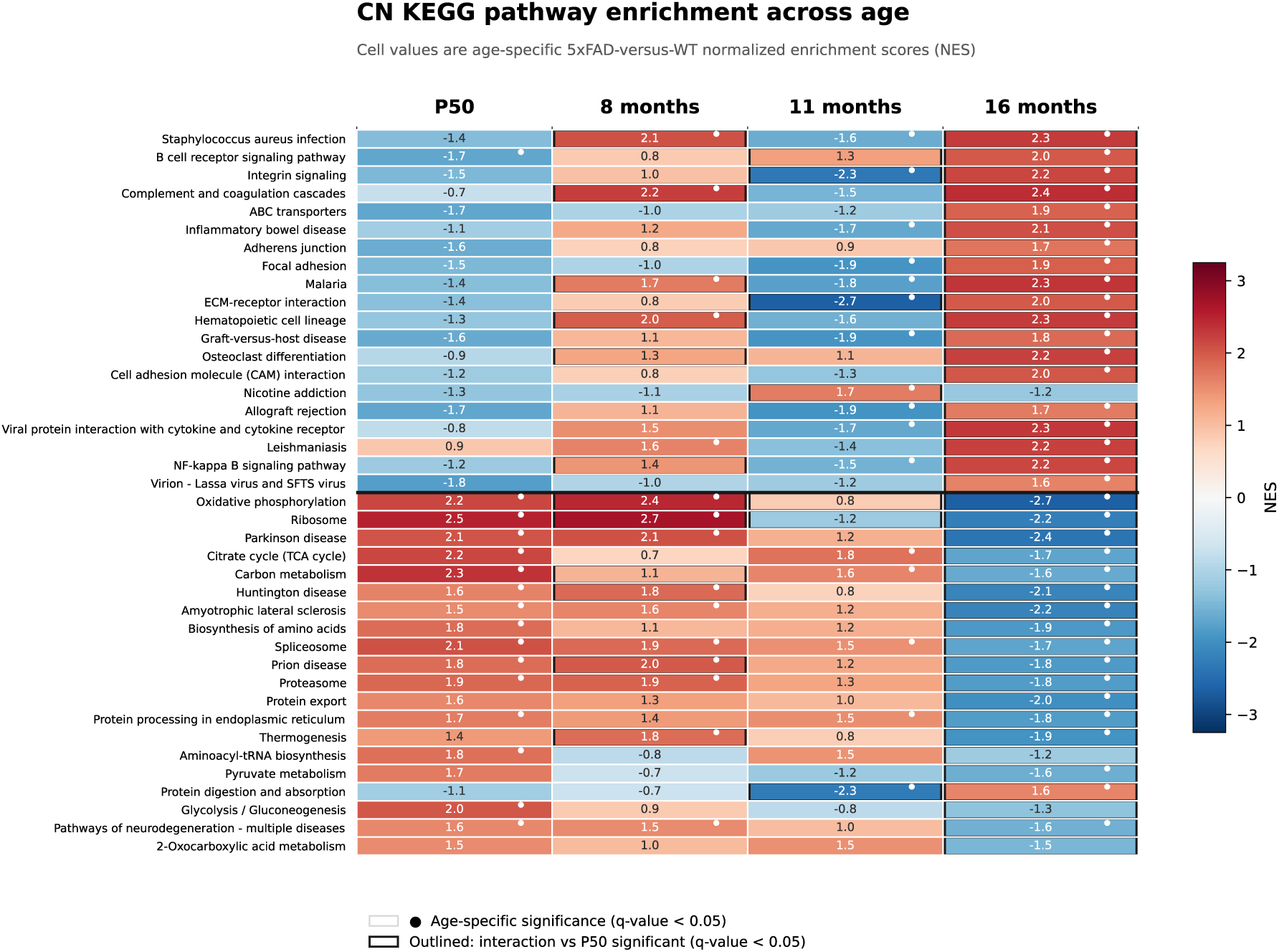
Age-dependent pathway enrichment in cochlear nucleus. Display, selection, and significance conventions are as in Figure S12. Columns show P50, 8, 11, and 16 months; dots identify significant age-specific enrichment and black outlines identify significant Genotype×Age interactions relative to P50.

**Figure S21.**
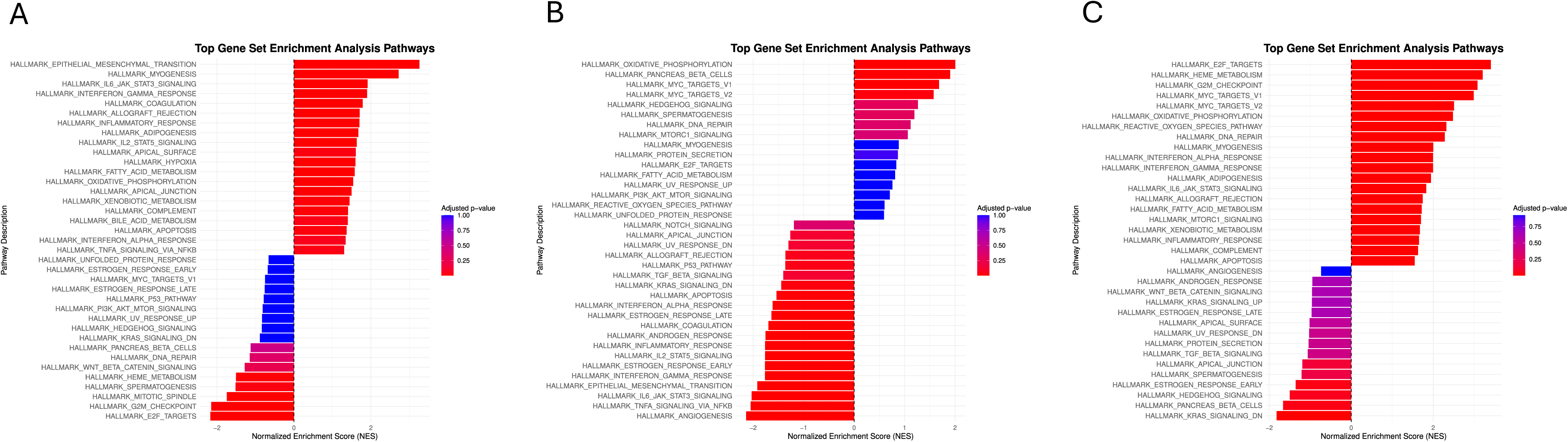
Age-dependent changes in Hallmark pathway enrichment in the cochlea i relative to P50.

**Figure S22.**
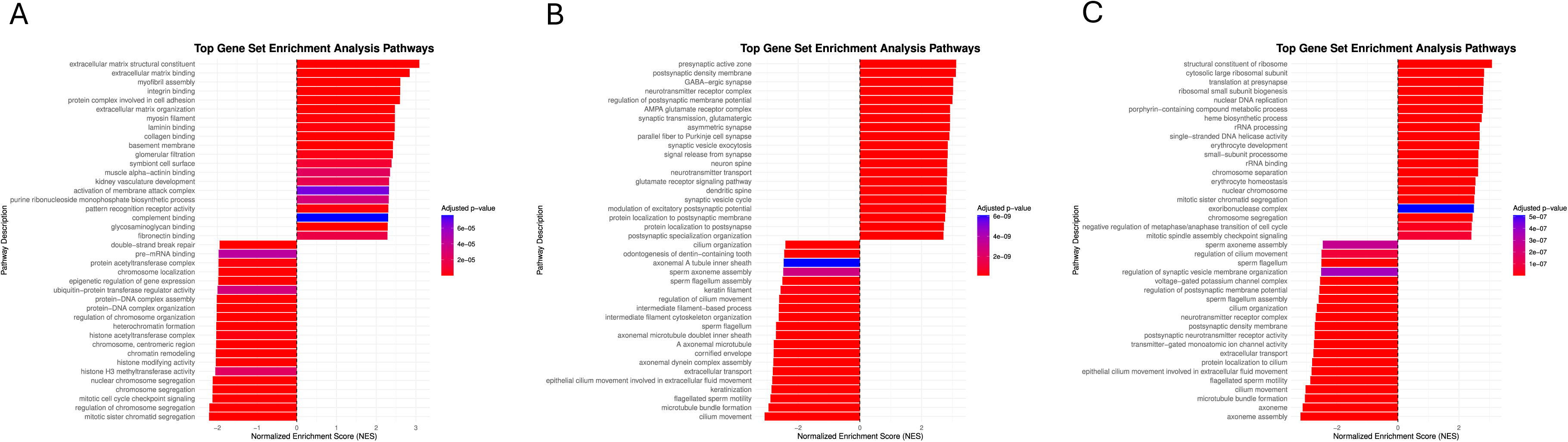
Age-dependent changes in GO pathway enrichment in the cochlea relative to P50.

**Figure S23.**
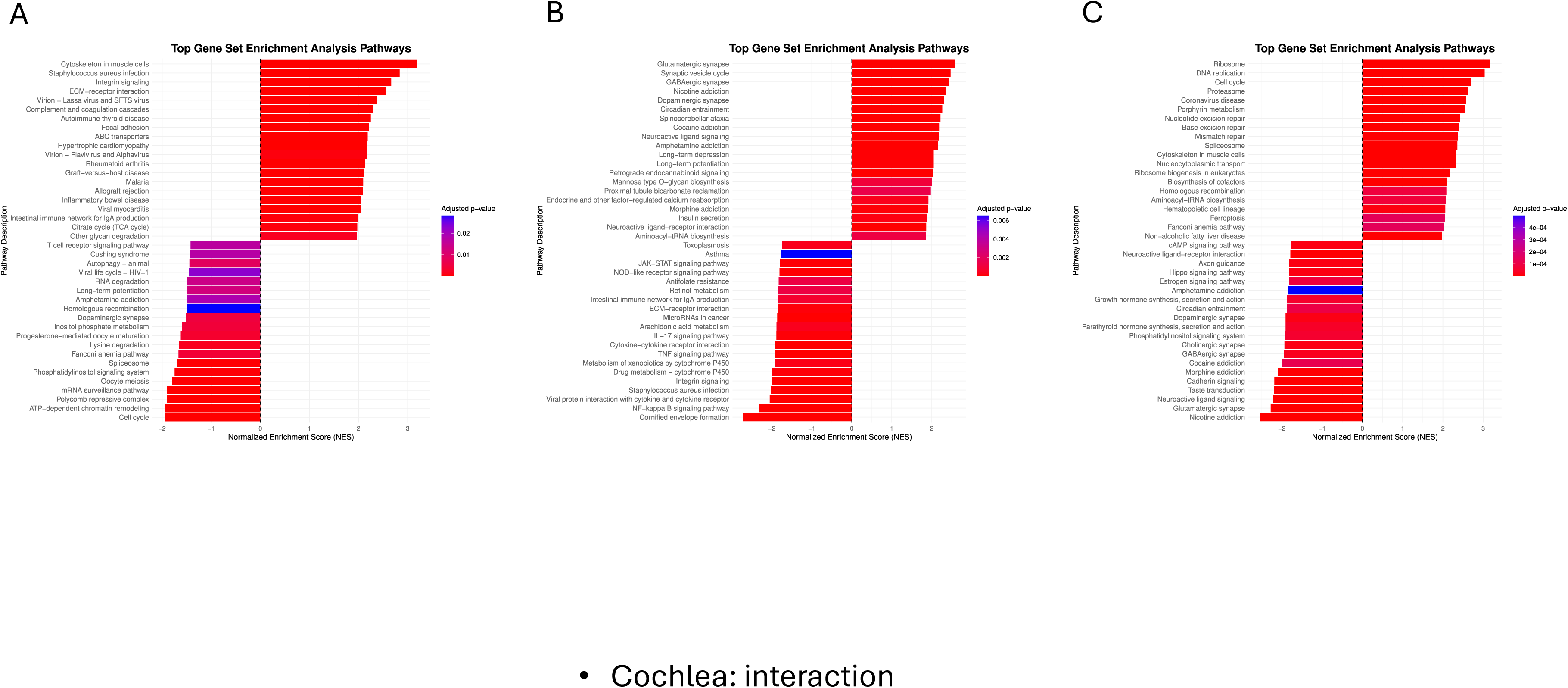
Age-dependent changes in KEGG pathway enrichment in the cochlea relative to P50.

**Figure S24.**
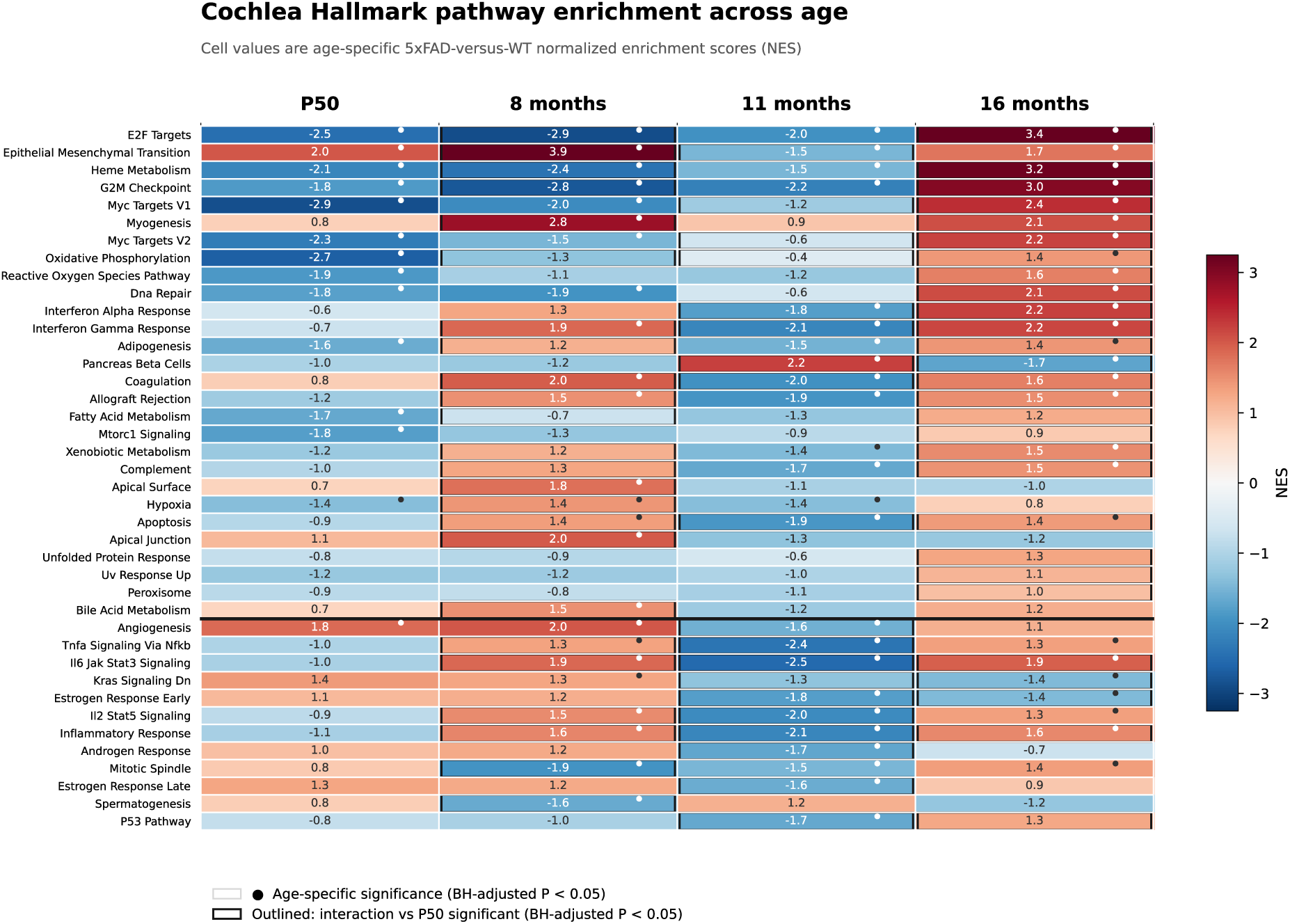
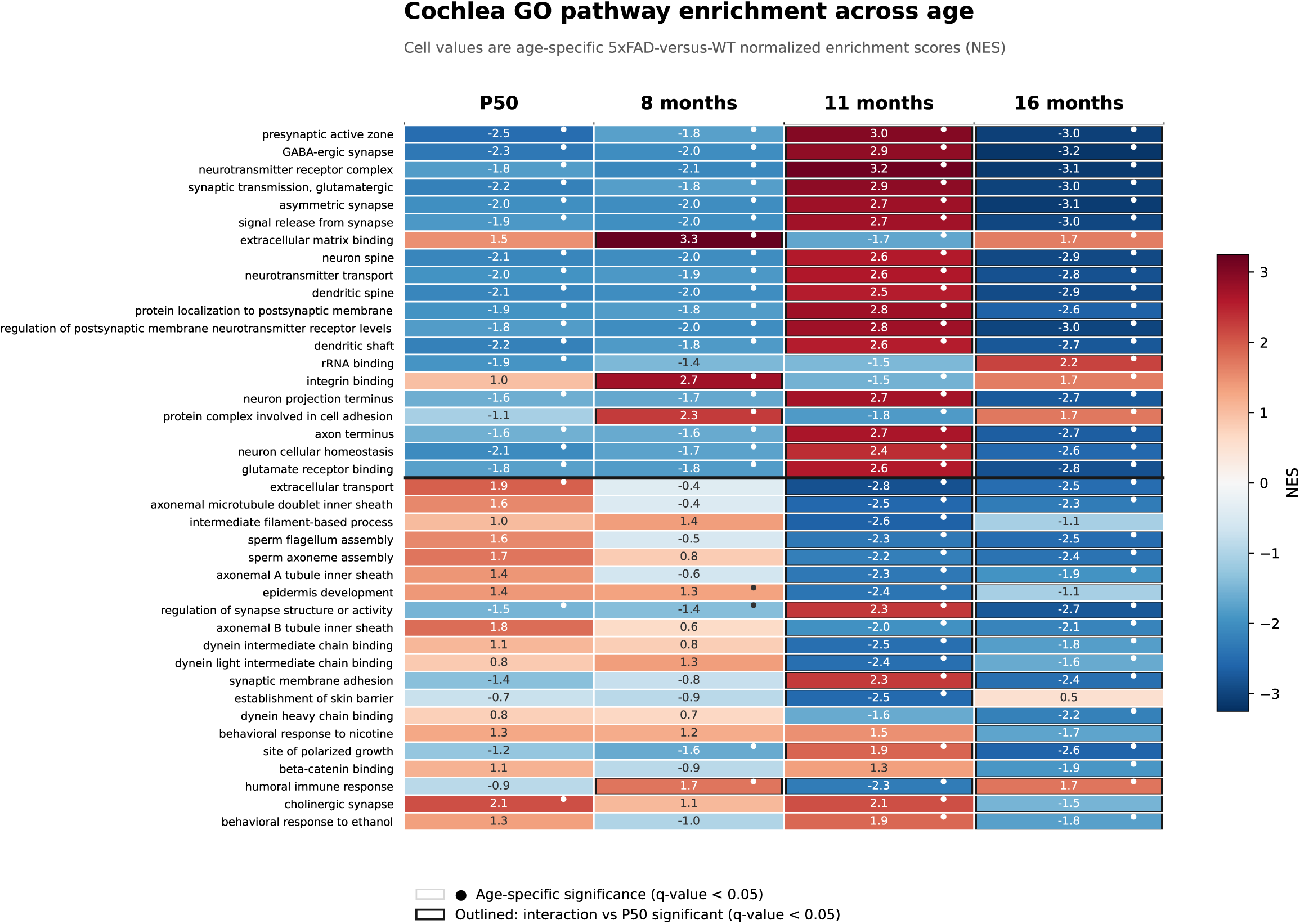
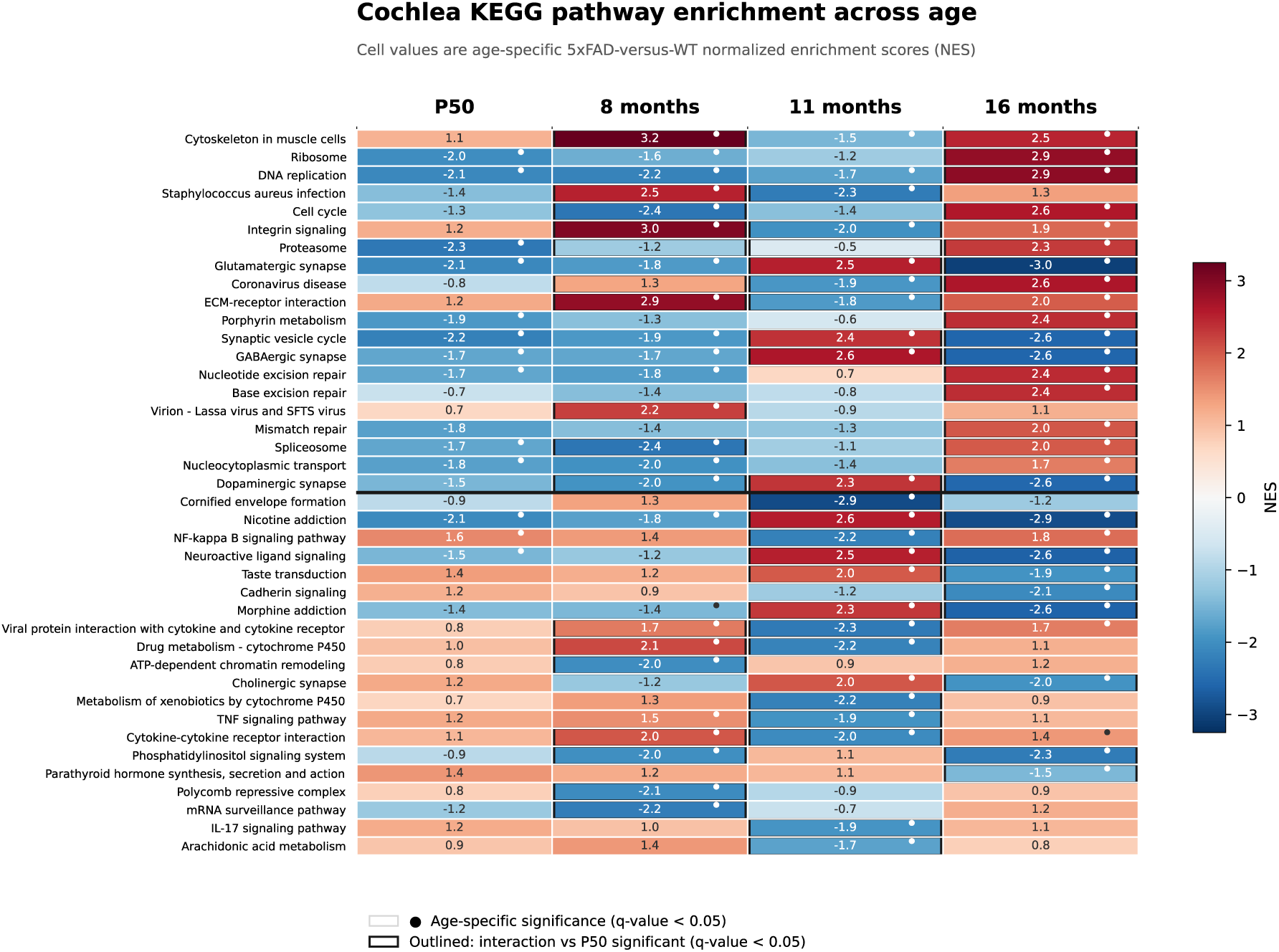
Age-dependent pathway enrichment in cochlea. Display, selection, and significance conventions are as in Figure S12. Columns show P50, 8, 11, and 16 months; dots identify significant age-specific enrichment and black outlines identify significant Genotype×Age interactions relative to P50.

**Figure S25.**
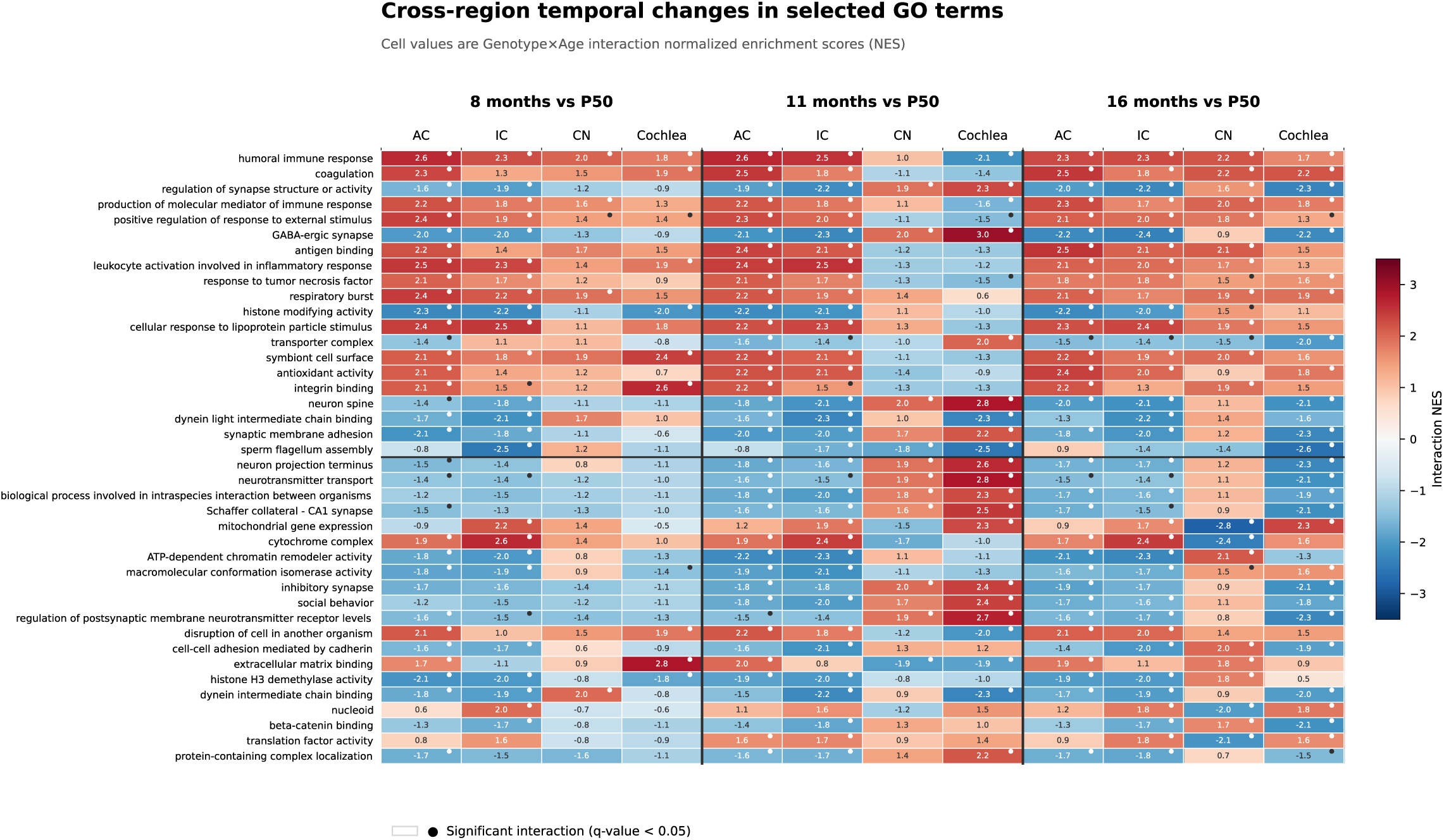
Cross-region temporal changes in selected GO pathways. The heatmap shows Genotype×Age interaction NES for AC, IC, CN, and cochlea at 8, 11, and 16 months relative to P50. The panel contains 40 exact GO identifiers: the 20 strongest broadly shared and 20 strongest divergent terms. Only identifiers retained after independent redundancy reduction in every regional age contrast were eligible. Columns are grouped by age, with regions ordered AC, IC, CN, and cochlea. Red and blue denote positive and negative interaction NES; dots mark q-value < 0.05. Interaction direction describes change from P50 rather than absolute later-age enrichment.

## References

Adaikkan C, Middleton SJ, Marco A, Pao PC, Mathys H, Kim DN, Gao F, Young JZ, Suk HJ, Boyden ES, McHugh TJ, Tsai LH. (2019) Gamma Entrainment Binds Higher-Order Brain Regions and Offers Neuroprotection. Neuron. 102(5):929–943.e8. doi: 10.1016/j.neuron.2019.04.011. PMID: 31076275

Bateman RJ, Xiong C, Benzinger TLS, et al. Clinical and biomarker changes in dominantly inherited Alzheimer’s disease. New England Journal of Medicine. 2012;367:795–804. doi:10.1056/NEJMoa1202753.

Buskila Y, Crowe SE, Ellis-Davies GCR. Synaptic deficits in layer 5 neurons precede overt structural decay in 5xFAD mice. Neuroscience. 2013;254:152–159. doi:10.1016/j.neuroscience.2013.09.016.

Dobin, A., Davis, C.A., Schlesinger, F., Drenkow, J., Zaleski, C., Jha, S., Batut, P., Chaisson, M., Gingeras, T.R., 2013. STAR: ultrafast universal RNA-seq aligner. Bioinformatics 29, 15–21. 10.1093/bioinformatics/bts635.

Durinck, S., Spellman, P.T., Birney, E., Huber, W., 2009. Mapping identifiers for the integration of genomic datasets with the R/Bioconductor package biomaRt. Nat Protoc 4, 1184–1191. 10.1038/nprot.2009.97

Gates GA, Anderson ML, McCurry SM, Feeney MP, Larson EB. Central auditory dysfunction as a harbinger of Alzheimer dementia. Archives of Otolaryngology–Head and Neck Surgery. 2011;137:390–395. doi:10.1001/archoto.2011.28.

Gates GA, Beiser A, Rees TS, D’Agostino RB, Wolf PA. Central auditory dysfunction may precede the onset of clinical dementia in people with probable Alzheimer’s disease. Journal of the American Geriatrics Society. 2002;50:482–488. doi:10.1046/j.1532-5415.2002.50114.x.

Hong S, Beja-Glasser VF, Nfonoyim BM, et al. Complement and microglia mediate early synapse loss in Alzheimer mouse models. Science. 2016;352:712–716. doi:10.1126/science.aad8373.

Iaccarino HF, Singer AC, Martorell AJ, Rudenko A, Gao F, Gillingham TZ, Mathys H, Seo J, Kritskiy O, Abdurrob F, Adaikkan C, Canter RG, Rueda R, Brown EN, Boyden ES, Tsai LH. (2016) Gamma frequency entrainment attenuates amyloid load and modifies microglia. Nature. 540: 230–235. doi: 10.1038/nature20587. PMID: 27929004

Kaylegian K, Stebritz AJ, Weible AP, Wehr M. 5XFAD mice show early onset gap detection deficits. Frontiers in Aging Neuroscience. 2019;11:66. doi:10.3389/fnagi.2019.00066.

Keren-Shaul H, Spinrad A, Weiner A, et al. A unique microglia type associated with restricting development of Alzheimer’s disease. Cell. 2017;169:1276–1290.e17. doi:10.1016/j.cell.2017.05.018.

Kong, Y., 2011. Btrim: a fast, lightweight adapter and quality trimming program for next-generation sequencing technologies. Genomics 98, 152–3. 10.1016/j.ygeno.2011.05.009

Lin FR, Metter EJ, O’Brien RJ, Resnick SM, Zonderman AB, Ferrucci L. Hearing loss and incident dementia. Archives of Neurology. 2011;68:214–220. doi:10.1001/archneurol.2010.362.

Liu Y, Fang S, Liu LM, Zhu Y, Li CR, Chen K, Zhao HB. Hearing loss is an early biomarker in APP/PS1 Alzheimer’s disease mice. Neurosci Lett. 2020 Jan 19;717:134705. doi: 10.1016/j.neulet.2019.134705. Epub 2019 Dec 20. PMID: 31870800; PMCID: PMC7004828.

Love, M.I., Huber, W., Anders, S., 2014. Moderated estimation of fold change and dispersion for RNA-seq data with DESeq2. Genome Biology 15, 550. 10.1186/s13059-014-0550-8

Martorell AJ, Paulson AL, Suk HJ, Abdurrob F, Drummond GT, Guan W, Young JZ, Kim DN, Kritskiy O, Barker SJ, Mangena V, Prince SM, Brown EN, Chung K, Boyden ES, Singer AC, Tsai LH. (2019) Multi-sensory Gamma Stimulation Ameliorates Alzheimer’s-Associated Pathology and Improves Cognition. Cell. 177: 256–271.e22. doi: 10.1016/j.cell.2019.02.014. PMID: 30879788.

Mei L, Liu LM, Chen K, Zhao HB. Early Functional and Cognitive Declines Measured by Auditory-Evoked Cortical Potentials in Mice With Alzheimer’s Disease. Front Aging Neurosci. 2021 Sep 13;13:710317. doi: 10.3389/fnagi.2021.710317. PMID: 34588972; PMCID: PMC8473830.

Na D, Zhang J, Beaulac HJ, et al. Increased central auditory gain in 5xFAD Alzheimer’s disease mice as an early biomarker candidate for Alzheimer’s disease diagnosis. Frontiers in Neuroscience. 2023;17:1106570. doi:10.3389/fnins.2023.1106570.

Oakley H, Cole SL, Logan S, et al. Intraneuronal beta-amyloid aggregates, neurodegeneration, and neuron loss in transgenic mice with five familial Alzheimer’s disease mutations: Potential Factors in Amyloid Plaque Formation. Journal of Neuroscience. 2006;26:10129–10140. doi:10.1523/JNEUROSCI.1202-06.2006.

O’Leary TP, Shin S, Fertan E, et al. Reduced acoustic startle response and peripheral hearing loss in the 5xFAD mouse model of Alzheimer’s disease. Genes, Brain and Behavior. 2017;16:554–563. doi:10.1111/gbb.12370.

Raglio, A., Bellandi, D., Baiardi, P., Gianotti, M., Ubezio, M. C., Zanacchi, E., Granieri, E., Imbriani, M., & Stramba-Badiale, M. (2015). Effect of active music therapy and individualized listening to music on dementia: A multicenter randomized controlled trial. Journal of the American Geriatrics Society, 63(8), 1534–1539.

Särkämö, T., Tervaniemi, M., Laitinen, S., Numminen, A., Kurki, M., Johnson, J. K., & Rantanen, P. (2014). Cognitive, emotional, and social benefits of regular musical activities in early dementia: Randomized controlled study. The Gerontologist, 54(4), 634–650.

Stevenson JS, Clifton L, Kuźma E, Littlejohns TJ. Speech-in-noise hearing impairment is associated with an increased risk of incident dementia in 82,039 UK Biobank participants. Alzheimer’s & Dementia. 2022;18:445–456. doi:10.1002/alz.12416.

Wang W, Huang R, Lv L, Ma X, Li Z, Zhang Y, Wu J, Wu S, Xu J, Hu Y, Turck CW, Li H, Hu X. (2026) Long-term effects of forty-hertz auditory stimulation as a treatment of Alzheimer’s disease: Insights from an aged monkey model study. Proc Natl Acad Sci USA. 2026 Jan 13;123(2):e2529565123. doi: 10.1073/pnas.2529565123. PMID: 41490490; PMCID: PMC12799102.

Weible AP, Stebritz AJ, Wehr M. 5XFAD mice show early-onset gap encoding deficits in the auditory cortex. Neurobiology of Aging. 2020;94:101–110. doi:10.1016/j.neurobiolaging.2020.05.013.

Weible AP, Wehr M. Amyloid pathology in the central auditory pathway of 5XFAD mice appears first in auditory cortex. Journal of Alzheimer’s Disease. 2022;89:1385–1402. doi:10.3233/JAD-220538.

Xu, S., Hu, E., Cai, Y., Xie, Z., Luo, X., Zhan, L., Tang, W., Wang, Q., Liu, B., Wang, R., Xie, W., Wu, T., Xie, L., Yu, G., 2024. Using clusterProfiler to characterize multiomics data. Nat Protoc 19, 3292–3320. 10.1038/s41596-024-01020-z

Zhai TY, Liang C, Chen J, Yang J, Kong Y, Zhu Y, Yu N, Zhao HB. Upregulation of ATP-purinergic P2×2 receptors in the cochlea over-amplifies hearing sensitivity leading to hyperacusis and attenuation by antagonists. bioRxiv [Preprint]. 2026 Jun 22:2026.06.17.733049. doi: 10.64898/2026.06.17.733049. PMID: 42395442; PMCID: PMC13320770.

Zhou Y, Song WM, Andhey PS, et al. Human and mouse single-nucleus transcriptomics reveal TREM2-dependent and TREM2-independent cellular responses in Alzheimer’s disease. Nature Medicine. 2020;26:131–142. doi:10.1038/s41591-019-0695-9.

